# Geospatial foundation models enable data-efficient tree species mapping in temperate mountain forests

**DOI:** 10.64898/2026.02.23.707022

**Authors:** James GC Ball, Jana Annika Wicklein, Zhengpeng Feng, Jovana Knezevic, Sadiq Jaffer, Anil Madhavapeddy, Clement Atzberger, Michele Dalponte, David Coomes

## Abstract

Accurate mapping of tree species from satellite data remains challenging in heterogeneous mountain forests due to environmental gradients, mixed stands, limited availability of high-purity training labels, and strong illumination-angle effects. Recent geospatial foundation models offer a new approach by learning generic, cloud-agnostic, information-rich representations from large multi-sensor archives suitable for a range of downstream tasks, but their ecological utility for species-level mapping remains incompletely understood. Here, we evaluate two geospatial foundation-model embeddings, AlphaEarth and Tessera, for tree species classification in the Trentino region of northern Italy, using parcel-level forest inventories as reference data (18 species and species groups). We compare their performance against conventional Sentinel-1+2 satellite composites across a series of controlled experiments examining classification accuracy, label efficiency, classifier complexity, robustness to label impurity, and temporal transferability. Foundation-model embeddings consistently outperform composite-based multispectral satellite baselines (weighted F1 = 0.83 vs. 0.80; macro F1 = 0.55 vs. 0.50), reaching near-asymptotic accuracy with as few as 5% of available training parcels and preserving ecologically meaningful structure aligned with functional and taxonomic groupings. However, realising this advantage requires a nonlinear classifier: a compact neural network provides better results than classic machine learning (i.e. Random Forest) and performs as well as deeper neural networks, while a linear classifier on foundation-model embeddings underperforms a neural network on conventional composites. Ancillary environmental covariates offer no additional classification benefit when added to embedding-based models. Classification accuracy remains robust to moderate levels of label impurity, allowing mixed parcels to be retained in the training dataset without substantial penalties, while training with parcel-level species proportions as soft labels achieves higher peak performance (macro F1 = 0.586 for Tessera, 0.589 for AlphaEarth) and lower Proportion L1 error than hard labels without requiring purity filtering, maximising the value of the full range of input data. However, temporal transfer across years reveals performance degradation, with weighted F1 declining by 9% for Tessera and 15% for AlphaEarth, and disproportionate losses for rare species. Overall, our results show that geospatial foundation models shift a primary bottleneck in species mapping from feature engineering toward the availability, quality, and temporal alignment of ecological reference data, while opening new opportunities for scalable biodiversity monitoring and the analysis of ecological change.

## 1. Introduction

Accurate, up-to-date regional maps of tree species are valuable for biodiversity assessment, carbon estimation, and evidence-based management, because species composition regulates productivity, disturbance susceptibility and resilience, as well as the delivery of key ecosystem services (Tilman et al., 2013; Brockerhoff et al., 2017; Schwaiger et al., 2018; Bai et al., 2024). This reinforces the need for species-resolved information in monitoring and policy (Thompson et al., 2009; Silva Pedro et al., 2015; Thrippleton et al., 2023; Germain and Lutz, 2024). Field inventories remain the most reliable source of species-level data, yet their sparse coverage and high per-unit cost of maintenance limit their utility at landscape and regional scales (McRoberts et al., 2010; Bergseng et al., 2015). National forest inventory programmes therefore increasingly integrate plot networks with airborne and satellite observations to expand spatial coverage cost-effectively, complementing traditional surveys with remote sensing products that capture broad environmental gradients and disturbance regimes (McRoberts and Tomppo, 2007; White et al., 2016; Lister et al., 2020; Immitzer and Atzberger, 2023). The need for such scalable approaches is especially acute in montane regions, where climate warming is driving elevational shifts of forest belts and composition, and amplifying disturbance dynamics – most notably drought, windthrow, and bark-beetle-linked mortality – across the European Alps and temperate Europe (Seidl et al., 2017; Senf and Seidl, 2021; Frei et al., 2023). The degree to which these rapid changes will impact ecosystem stability and local extinction remains uncertain (Scherrer et al., 2020), underscoring the urgency of developing robust, repeatable species mapping frameworks to monitor forest trajectories and support adaptive management in mountain ecosystems (European Commission, 2021; De Keersmaecker et al., 2024).

Traditional satellite-based approaches remain constrained in their ability to discriminate tree species reliably over large, heterogeneous landscapes (Fassnacht et al., 2016). Although multispectral sensors such as Landsat and Sentinel-2 provide several broad spectral bands, the provided data is often insufficient to detect subtle canopy variations associated with species-specific differences in leaf biochemistry and pigment concentrations (Dalponte et al., 2012; Fassnacht et al., 2016; Immitzer et al., 2019; Zhong et al., 2024). The deca-metric spatial resolution (10–30 m) further leads to mixed pixels that obscure individual crowns or fine-scale species mosaics characteristic of mountain forests (Xu et al., 2021).

Recent advances in geospatial foundation models (GFMs) offer a step change in the feasibility of large-scale tree species mapping. These models, trained once on massive multi-sensor archives using self-supervised objectives, learn information-rich embeddings that capture spectral and temporal features of the Earth’s surface (Xiao et al., 2025). Unlike traditional approaches that rely on handcrafted features, site-specific calibration, or extensive labelled datasets, GFMs distil geospatially relevant patterns directly from unlabelled imagery, allowing downstream classifiers to be trained efficiently on far smaller sets of local labels – one of the main bottlenecks in species-level remote sensing (Ball et al., 2024). Models such as AlphaEarth (Brown et al., 2025) and Tessera (Feng et al., 2025) exemplify this shift, providing latent representations derived from petabyte-scale, globally distributed, satellite time series that can be transferred to regional tasks even where labelled data are sparse or heterogeneous. Species identification using individual images is confounded by strong phenological variation (Andreatta et al., 2025; Dalponte et al., 2025), canopy understory variability (Rautiainen et al., 2011) and snow seasonality (Wang et al., 2023), meaning single-date imagery rarely captures optimal discriminative conditions. Multi-season time series, on the other hand, are cumbersome to process, involving vast arrays of data often impacted by irregular cloud cover (Wang et al., 2022). In principle, fusing complementary modalities – such as synthetic aperture radar and optical reflectance – can mitigate ambiguities (Blickensdörfer et al., 2024), but multimodal integration has historically required bespoke pre-processing, cross-sensor alignment/harmonisation, and task-specific modelling pipelines. Topographic effects introduce additional complexity: strong Bidirectional Reflectance Distribution Function (BRDF) effects result from variable illumination, shadowing, and slope-induced changes in reflectance and backscattering signals, reducing the transferability of signatures across terrain (Dong et al., 2020; Kellndorfer et al., 1998). As a result, models trained for one site or season often do not generalize to other areas (Tuia et al., 2016). These limitations have historically confined detailed tree species mapping to small study areas or a narrow set of species or species groups, hindering the development of robust, scalable classification frameworks.

Despite this promise, several fundamental questions about how abstract representations relate to ground reality remain unresolved. First, although foundation models have been applied to vegetation and forest monitoring, their use for tree species-level mapping from satellite imagery remains limited (Braham et al., 2025), and it is unclear whether their learned representations reliably encode species-discriminative information in structurally complex, heterogeneous forests in mountainous terrain (Brandt et al., 2025; Ishikawa et al., 2025; Jin and Wu, 2025). A central claim of foundation models is improved label efficiency, yet few studies quantify how performance scales with labelled data under a fixed evaluation design. In forest inventories – where species labels are costly, spatially uneven, and often aggregated at the stand or parcel level – it remains unclear how much labelled data are required for reliable species discrimination, or how additional sampling effort translates into performance gains (Brus et al., 2012; Ramezan et al., 2021). Second, the downstream classifier capacity needed to fully exploit the information encoded in GFM embeddings has not been systematically assessed. If embeddings already capture species-discriminative structure, simple classifiers may suffice; if not, increased complexity may be necessary to achieve strong performance and could change conclusions about representation quality. A further challenge arises from label noise and mixed-species stands, which are common in real inventories. Mono-specific (pure) parcels are often rare, and aggressive filtering inevitably trades label quality against data volume. This raises the open question of whether the structure of foundation-model representations can be leveraged to make use of noisy or aggregated labels – such as parcel-level species mixtures – to improve species discrimination, or whether such complexity yields limited practical benefit (Bolyn et al., 2022). Finally, operational forest monitoring requires models that generalise across both space and time. Although GFMs are trained on multi-year archives, evaluations typically focus on within-year performance, leaving the temporal stability of learned representations largely untested. It therefore remains unclear whether embeddings learned in one year retain species-discriminative power under interannual variation in phenology and acquisition conditions (Lisaius et al., 2026).

In this study, we evaluate whether globally pre-trained geospatial foundation models can serve - without any further fine tuning - as effective and data-efficient representations for regional-scale tree species mapping in complex mountain forests. Using forest cover in the Autonomous Province of Trento (Italy) as a demanding test case – characterised by steep environmental gradients, mixed stands, strong phenological variability, and detailed parcel-level forest inventories – we compare embeddings from two foundation models, AlphaEarth and Tessera, with conventional Sentinel-1+2 spectral composites. Our analysis focuses not on proposing a new classification algorithm, but on assessing the ecological information encoded in each representation. Specifically, we ask:

1. Do GFM embeddings encode species-discriminative information more effectively than conventional satellite composites?
2. Does increasing downstream classifier capacity lead to consistent performance gains when applied to geospatial foundation-model embeddings?
3. How sensitive is classification performance to training-data availability and label purity?
4. Do supplementary environmental variables provide additional predictive power beyond satellite-derived representations?
5. How does classification performance change under cross-year transfer?

Through this analysis, we test whether globally pre-trained geospatial foundation models can serve – without further fine-tuning – as a basis for regional, species-level biodiversity monitoring and ecosystem assessment. Using a temperate montane forest as a case study, we evaluate their ability to discriminate dominant tree species relative to conventional remote-sensing representations, and examine how performance depends on classifier capacity, training-data availability, label purity, and temporal transfer. To our knowledge, this is the first study to attempt to map detailed distributions from space using geospatial foundation models.

## 2. Methods

### 2.1. Conceptual Overview

This study evaluates whether geospatial foundation-model (GFM) embeddings provide ecologically meaningful and data-efficient representations for tree-species classification. We compare two globally pre-trained GFM embeddings (AlphaEarth; Brown et al., 2025 and Tessera; Feng et al., 2025) with conventional Sentinel-1/2 multispectral composites under controlled and comparable modelling conditions.

All training and prediction was conducted at the pixel level (while evaluation was also conducted at the parcel level), producing discrete species predictions compatible with existing wall-to-wall forest maps (e.g. European genus-level products; De Keersmaecker et al., 2024). Although sub-pixel or fractional approaches are increasingly used to represent mixed stands (e.g. Bolyn et al., 2022), we adopt a standard pixel-wise classification setting to minimise additional modelling assumptions and to keep the focus on representation choice and downstream model behaviour.

Classifier capacity is treated as a controlled experimental axis. Here, classifier capacity refers to the ability of a model to represent complex, nonlinear decision boundaries and higher-order feature interactions, governed by its functional form and number of learnable parameters. We evaluate a set of commonly used classifiers spanning increasing expressive capacity, from linear models and instance-based methods to ensemble and neural-network approaches. In particular, Random Forests (RFs) are used as a conservative and widely adopted baseline in ecological remote sensing, while multi-layer perceptrons (MLPs) represent a more flexible nonlinear alternative. This progression allows us to assess whether performance differences are primarily driven by representation quality or by downstream model capacity.

Model behaviour is assessed along five dimensions: (i) overall classification performance and the ecological structure reflected in predictions; (ii) label efficiency as training data availability is reduced; (iii) sensitivity to training-label purity and robustness to mixed parcels; (iv) the contribution of ancillary environmental covariates beyond satellite-derived representations; and (v) temporal generalisation under cross-year transfer. Results are presented by first introducing a reference configuration, followed by targeted analyses that vary one experimental factor at a time (ablation studies).

A schematic overview of the modelling pipeline is provided in Figure 1 and evaluation process in Figure 2. Detailed descriptions of individual components are presented in the following sections. Unless otherwise stated, results refer to an MLP classifier trained on 2018 embeddings using parcel-level cross-validation, with training labels filtered to ≥60 % dominant species cover.

**Figure 1:**
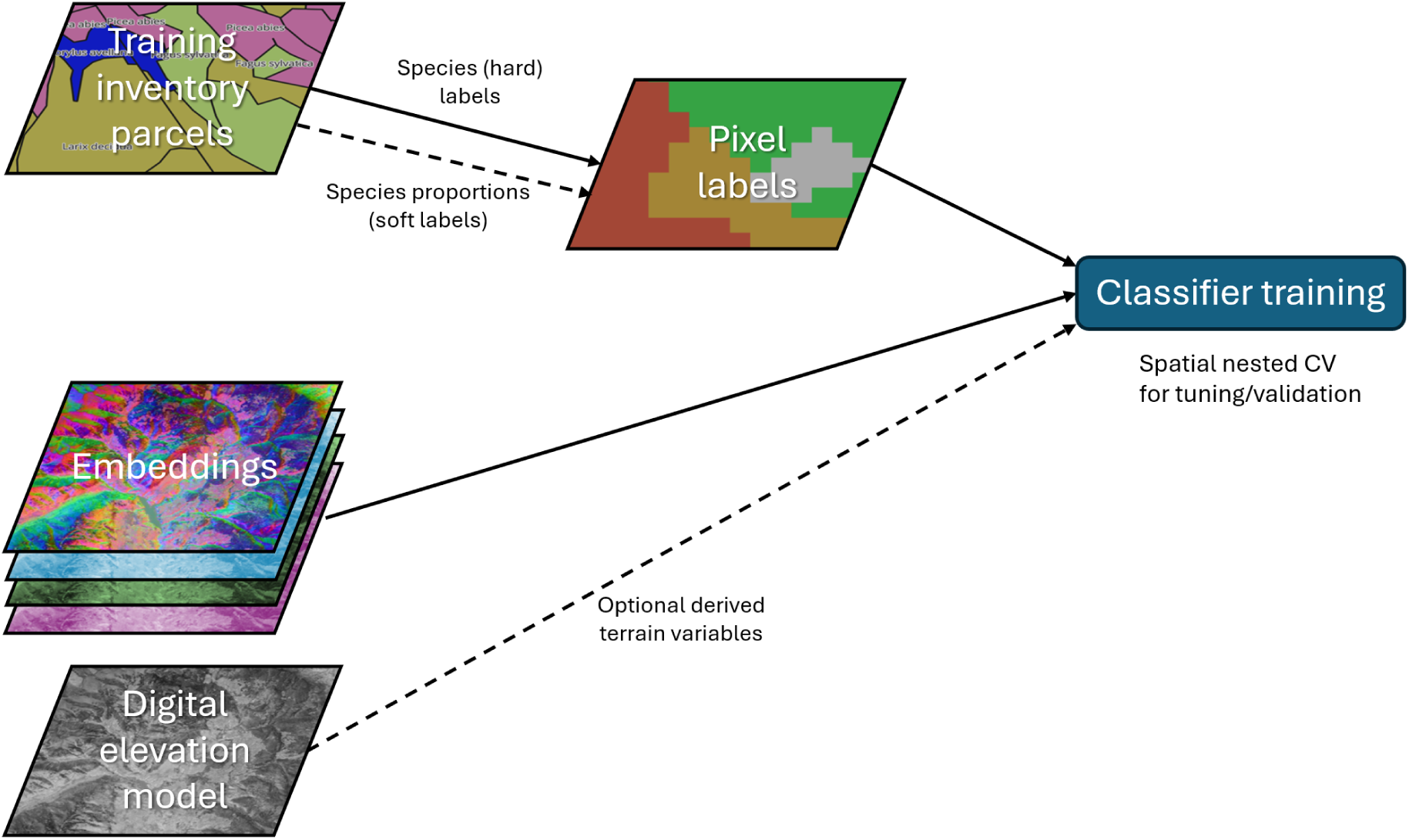
Conceptual overview of the species classification workflow. Parcel-level forest inventory data are rasterised to pixel-level labels or as mixed composition soft labels. Geospatial foundation-model embeddings provide pixel-level feature representations for classifier training. Optional terrain variables are included in ablation experiments.

**Figure 2:**
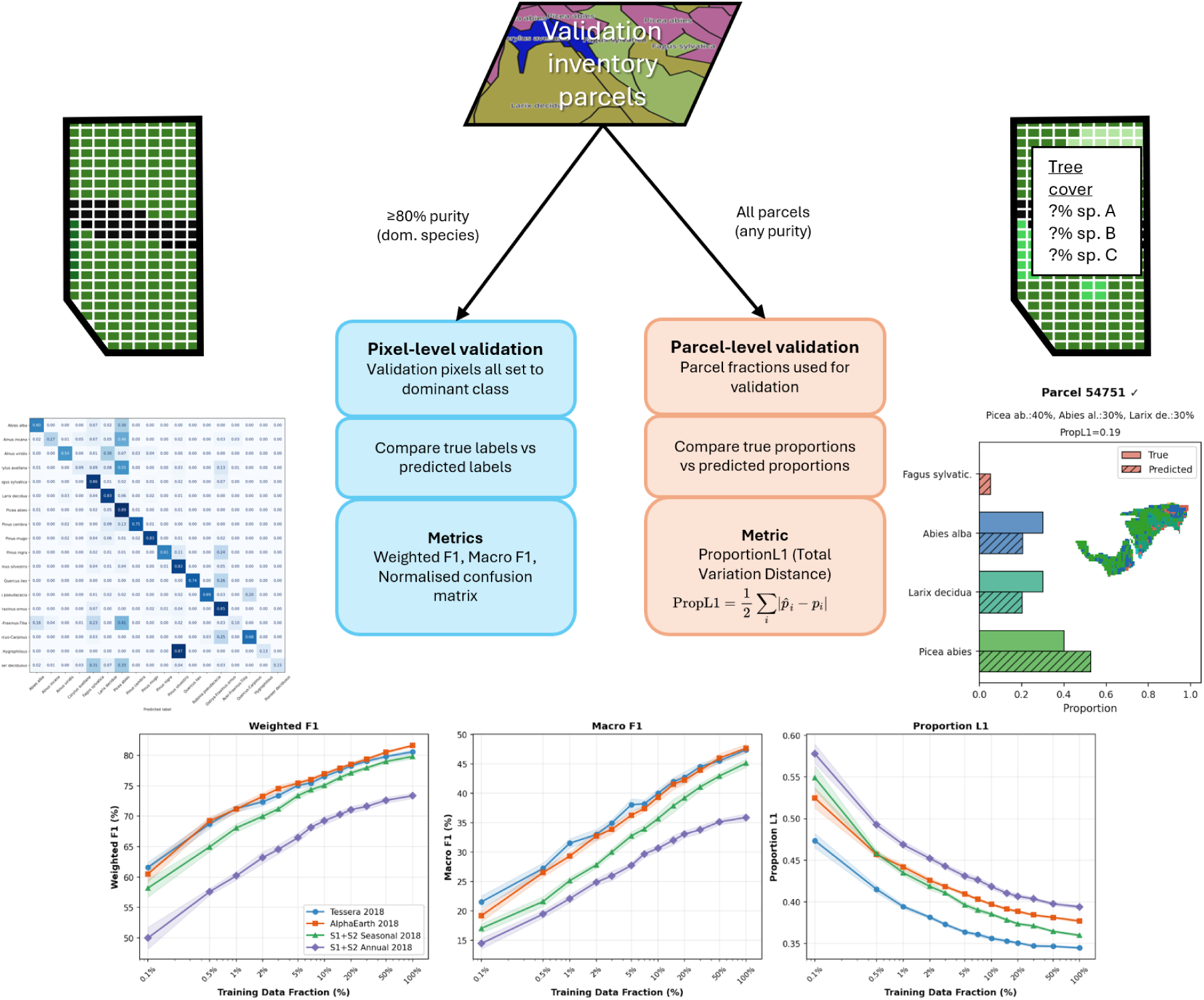
Schematic of the dual validation framework. Validation inventory parcels (top) are evaluated at two complementary levels. Left branch (pixel-level): parcels with dominant-species purity ≥ 80% are selected and all pixels are assigned the dominant species label; per-pixel predictions are compared against these labels to compute weighted F1, macro F1, and a normalised confusion matrix. Right branch (parcel-level): all parcels, including mixed-species stands, are used; pixel-level predictions are aggregated into predicted species proportions and compared against true inventory fractions using the Proportion L1 metric (Total Variation Distance). Black pixels represent filtered non-forest classes such as roads. Example outputs from each branch are shown at the bottom.

**Figure 3:**
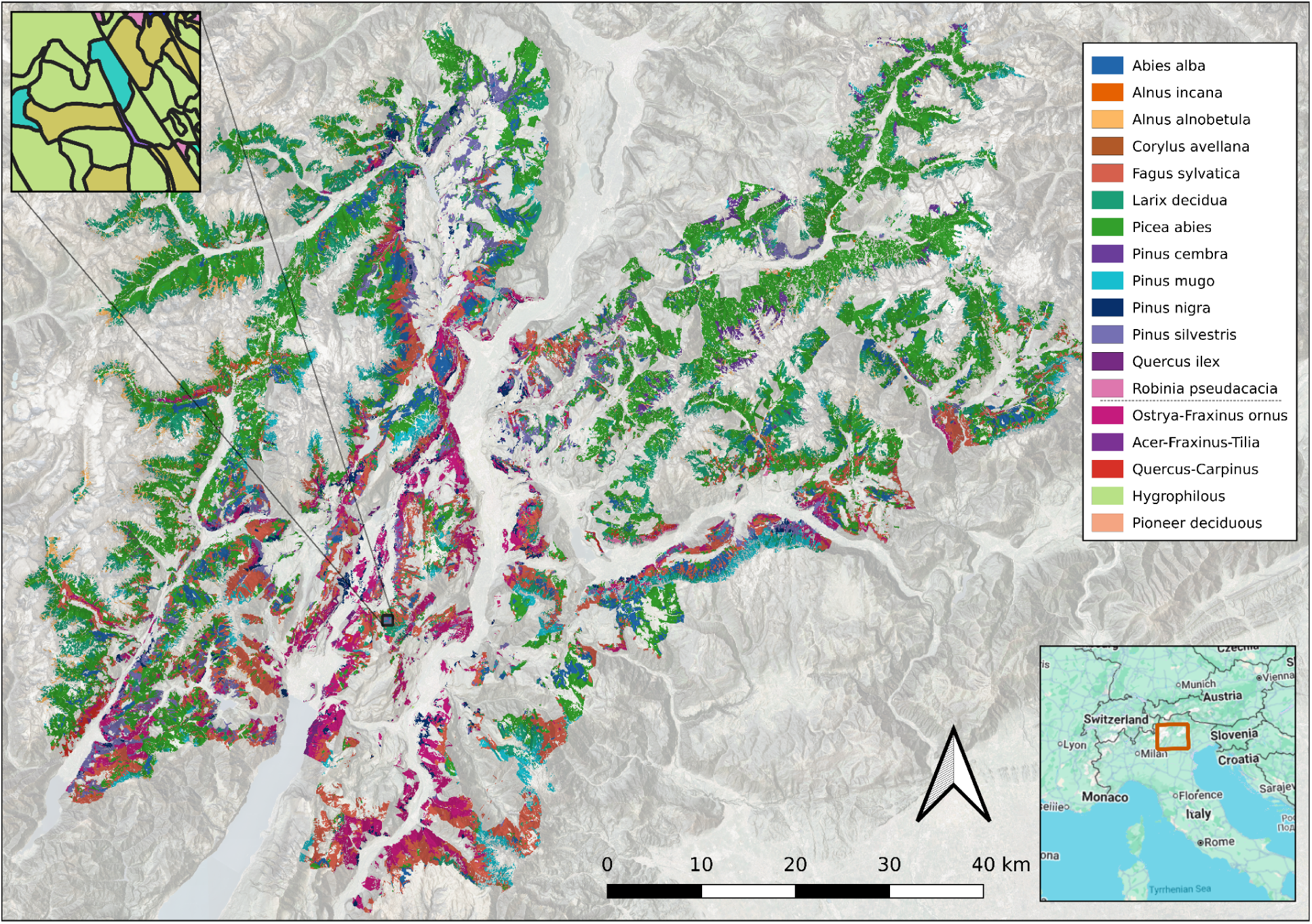
Trentino with the colours showing 18 labelled tree species and species group classes of the surveyed forest plots. The top-left inset shows a zoom on a dense section of the irregular forest inventory parcels which define the pixel-level species labels within them. The bottom-right inset shows the study region within its context of northern Italy.

### 2.2. Study region and forest inventory

#### 2.2.1. Study region: Trentino, Italy

The Autonomous Province of Trento (Trentino), Italy, provides a challenging and ecologically realistic test case for evaluating geospatial foundation models for species-level forest mapping. The region spans steep elevational, climatic, and biogeographic gradients over short distances, producing sharp transitions between distinct forest communities (Agnoletti and Biasi, 2013; Pedrotti, 2018). This fine-grained heterogeneity, coupled with strong phenological variation and complex illumination from rugged terrain, routinely undermines traditional optical satellite-based species classification (Weiss and Walsh, 2009; Hu et al., 2025). These challenges extend to radar-based observations, as mountainous terrain introduces pronounced layover, shadowing, and slope-orientation effects due to the oblique viewing geometry of SAR sensors. As a result, Trentino represents a stringent test of whether satellite-derived representations can encode species-discriminative information that is robust to terrain, forest structure, and mixed stand composition.

These gradients produce a pronounced elevational zonation of forest types. Forests dominate much of the landscape between roughly 600 and 2,200 m. Below ∼600 m, valley floors are largely dominated by agriculture, urban areas, and fragmented riparian woodlands. At lower montane elevations (600–1,200 m), stands are primarily composed of European beech (*Fagus sylvatica*), often mixed with oak, hornbeam, and chestnut on warmer, south-facing slopes. Moving upslope into the submontane–montane conifer belt (1,200–1,800 m), extensive Norway spruce (*Picea abies*) and silver fir (*Abies alba*) forests form some of the most productive timber zones in the Italian Alps. The subalpine belt (1,800–2,200 m) is characterised by mixed stands of European larch (*Larix decidua*) and stone pine (*Pinus cembra*), frequently interspersed with patches of spruce and open grassy clearings (Gasparini et al., 2022). Above the climatic treeline (typically >2,000–2,200 m), forests transition into alpine meadows, dwarf-shrub communities dominated by species such as Rhododendron ferrugineum, and extensive scree, cliff, and rock habitats.

#### 2.2.2. Inventory data

The region benefits from one of the most detailed and spatially complete forest surveys in Europe, enabling a controlled yet demanding evaluation. As reference data, we used the forest assessment plan (*piani di assestamento*) survey provided by the Forest Service of the Province of Trento (*Servizio Foreste Provincia autonoma di Trento*) collected from 2010-2021 (see Fig S.1 for collection by year). The inventory divides the regional forest area into management units (irregularly shaped polygons) that typically correspond to ownership parcels or structural boundaries such as rivers and roads. The dataset comprises 83,000 individual forestry units with a mean area of 2.89 ha and maximum area of 519 ha. Collectively, these units cover a total of 260,000 ha. For each unit, the dataset reports the tree species present and each species’ proportional cover within the upper canopy layer. The number of tree species per unit varies from a single species to fifteen, with a mean of 3.55 species. Parcels under 0.2 ha were excluded from the analysis as they were considered uninformative given pixel size and co-registration uncertainty.

In consultation with local forestry experts and using coexistence analysis (Jaccard similarity and hierarchical clustering; see S.2), we applied a merging scheme that groups less dominant tree species into ecologically meaningful community classes based on their co-occurrence patterns and habitat affinities resulting in 13 dominant species retained individually and five community groups (see Table 1 and Section S.2 for details).

**Table 1.**
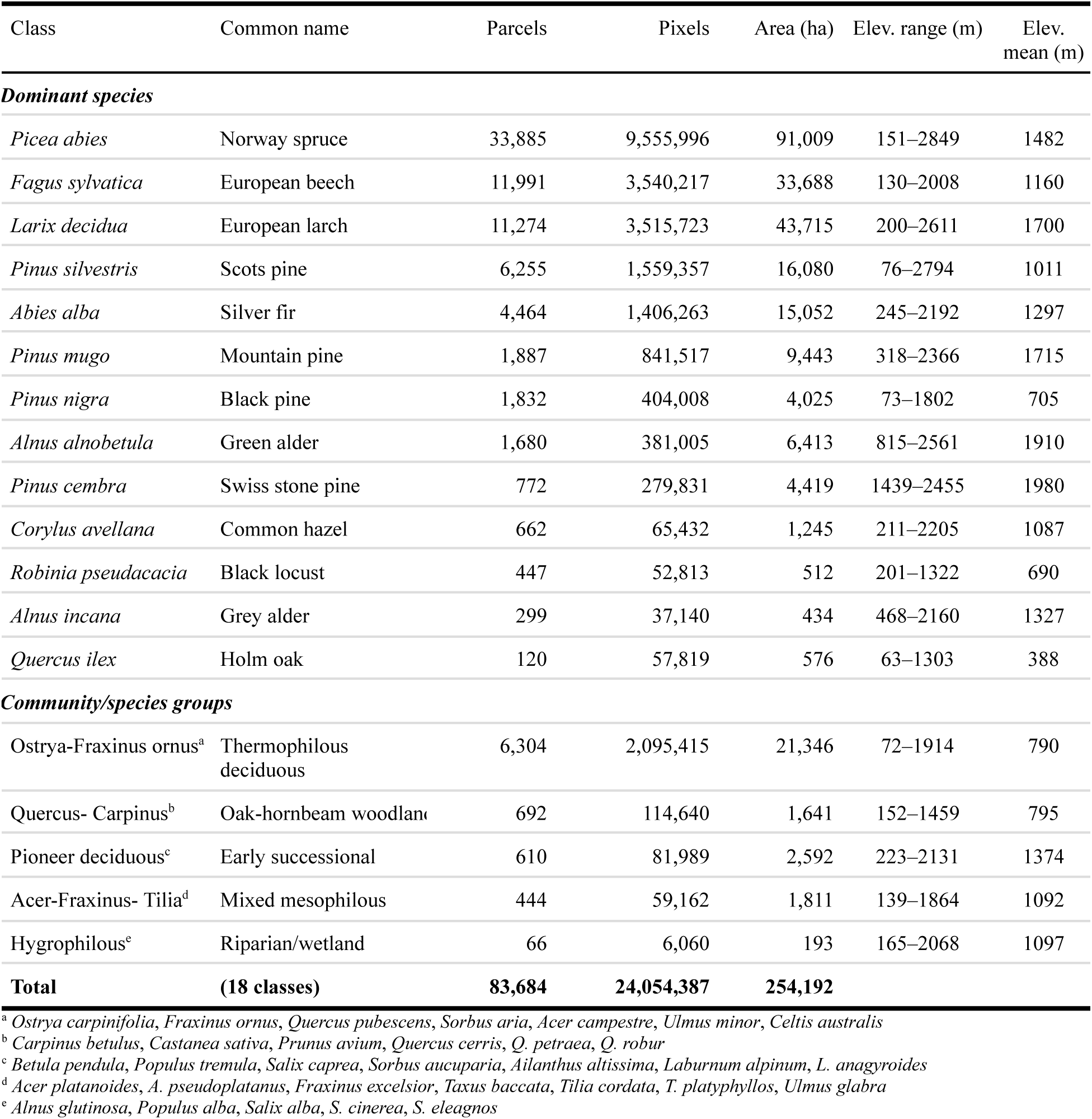
Target classes with training sample sizes and elevation distributions. The 18 classes comprise 13 dominant species retained as individual classes and 5 community groups formed by merging ecologically co-occurring minor species. Note that the pixel count is derived from pixels contained in parcels in which the species is dominant and area is calculated from the proportional cover of the forestry parcels, so correspondence is not exact.

| Class | Common name | Parcels | Pixels | Area (ha) | Elev. range (m) | Elev. mean (m) |
| --- | --- | --- | --- | --- | --- | --- |
| ***Dominant species*** |  |  |  |  |  |  |
| *Picea abies* | Norway spruce | 33,885 | 9,555,996 | 91,009 | 151–2849 | 1482 |
| *Fagus sylvatica* | European beech | 11,991 | 3,540,217 | 33,688 | 130–2008 | 1160 |
| *Larix decidua* | European larch | 11,274 | 3,515,723 | 43,715 | 200–2611 | 1700 |
| *Pinus silvestris* | Scots pine | 6,255 | 1,559,357 | 16,080 | 76–2794 | 1011 |
| *Abies alba* | Silver fir | 4,464 | 1,406,263 | 15,052 | 245–2192 | 1297 |
| *Pinus mugo* | Mountain pine | 1,887 | 841,517 | 9,443 | 318–2366 | 1715 |
| *Pinus nigra* | Black pine | 1,832 | 404,008 | 4,025 | 73–1802 | 705 |
| *Alnus alnobetula* | Green alder | 1,680 | 381,005 | 6,413 | 815–2561 | 1910 |
| *Pinus cembra* | Swiss stone pine | 772 | 279,831 | 4,419 | 1439–2455 | 1980 |
| *Corylus avellana* | Common hazel | 662 | 65,432 | 1,245 | 211–2205 | 1087 |
| *Robinia pseudacacia* | Black locust | 447 | 52,813 | 512 | 201–1322 | 690 |
| *Alnus incana* | Grey alder | 299 | 37,140 | 434 | 468–2160 | 1327 |
| *Quercus ilex* | Holm oak | 120 | 57,819 | 576 | 63–1303 | 388 |
| ***Community/species groups*** |  |  |  |  |  |  |
| Ostrya-Fraxinus ornus<sup>a</sup> | Thermophilous deciduous | 6,304 | 2,095,415 | 21,346 | 72–1914 | 790 |
| Quercus- Carpinus<sup>b</sup> | Oak-hornbeam woodland | 692 | 114,640 | 1,641 | 152–1459 | 795 |
| Pioneer deciduous<sup>c</sup> | Early successional | 610 | 81,989 | 2,592 | 223–2131 | 1374 |
| Acer-Fraxinus- Tilia<sup>d</sup> | Mixed mesophilous | 444 | 59,162 | 1,811 | 139–1864 | 1092 |
| Hygrophilous<sup>e</sup> | Riparian/wetland | 66 | 6,060 | 193 | 165–2068 | 1097 |
| **Total** | **(18 classes)** | **83,684** | **24,054,387** | **254,192** |  |  |

<sup>a</sup> *Ostrya carpinifolia*, *Fraxinus ornus*, *Quercus pubescens*, *Sorbus aria*, *Acer campestre*, *Ulmus minor*, *Celtis australis*
<sup>b</sup> *Carpinus betulus*, *Castanea sativa*, *Prunus avium*, *Quercus cerris*, *Q. petraea*, *Q. robur*
<sup>c</sup> *Betula pendula*, *Populus tremula*, *Salix caprea*, *Sorbus aucuparia*, *Ailanthus altissima*, *Laburnum alpinum*, *L. anagyroides*
<sup>d</sup> *Acer platanoides*, *A. pseudoplatanus*, *Fraxinus excelsior*, *Taxus baccata*, *Tilia cordata*, *T. platyphyllos*, *Ulmus glabra*
<sup>e</sup> *Alnus glutinosa*, *Populus alba*, *Salix alba*, *S. cinerea*, *S. eleagnos*

**Table 2.** Comparison of remote sensing predictors used to train the species classification models. For all layers, years 2018 (for training/validation) and 2019 (for transferability) were acquired which fall within the inventory collection period (2010-2021).

| Input | Features | Data type | Resolution | Modality |
| --- | --- | --- | --- | --- |
| AlphaEarth | 64 | GFM embeddings | 10 m | Optical + SAR |
| Tessera | 128 | GFM embeddings | 10 m | Optical + SAR |
| S1+S2 seasonal | 60 | Seasonal composite | 10 m | Optical + SAR |
| S1+S2 annual | 20 | Annual. composite | 10 m | Optical+ SAR |

### 2.3. Remote sensing inputs

#### 2.3.1. Geospatial foundation model embeddings

Geospatial foundation models produce embeddings, which are compact numerical representations of each pixel that summarise high-dimensional remote-sensing information – such as spectral signatures, temporal dynamics, and cross-sensor relationships – into a fixed-length vector (Xiao et al., 2025). These vectors inhabit a latent space, where proximity between points often corresponds to similarity in underlying surface properties according to the representation learned during self-supervised training. In this space, pixels with similar land-cover composition, phenology, or structure tend to cluster together, while dissimilar surfaces are separated by larger distances. Importantly, the latent space provides a continuous geometry in which operations such as measuring distances, identifying neighbourhoods, or anchoring representations to reference examples become meaningful.

**AlphaEarth Foundations (AEF)** is a global geospatial foundation model that produces annual 64-dimensional embeddings for every 10 m land pixel. It encodes multi-source remote sensing data from Sentinel-1 SAR, Sentinel-2 optical, and Landsat optical and thermal imagery. In addition to these AEF input sources, the approach uses additional target variables spanning a broader set of modalities, including LiDAR structure (GEDI), climate variables (ERA5), gravity fields (GRACE), and topography (DEM), implemented via implicit decoder objectives. Instead of using single images, representations are derived from satellite “video sequences” using a Space-Time-Precision (STP) encoder that supports continuous time, enabling the integration of irregularly sampled observations into a consistent temporal representation. During training, a student network is optimized to reproduce the embeddings of a teacher network via a consistency loss, encouraging robustness to missing or perturbed inputs, while a text-contrastive objective introduces a weak alignment with geolocated text descriptions. Together, these design choices aim to produce surface representations that are transferable across downstream tasks. The resulting embeddings are accessible via Google Earth Engine. Further details are provided in Brown et al. (2025).

**Tessera** is an open geospatial foundation model that generates annual 128-dimensional embeddings for every 10 m Sentinel pixel by jointly modelling optical and SAR satellite time series (Feng et al., 2025). Building on the concept of the spectral-temporal Barlow Twins (Lisaius et al., 2024), the model adopts a dual-encoder architecture in which annual time series of Sentinel-2 optical reflectance and Sentinel-1 SAR backscatter sequences are processed through separate encoders and fused into a unified representation. Tessera is trained on a global dataset using a self-supervised Barlow Twins objective (Zbontar et al., 2021; Lisaius et al., 2024) combined with sparse random temporal sampling, encouraging invariance to missing observations and irregular acquisition patterns. Robustness is further promoted through global shuffling, which reduces spatial autocorrelation, and mix-up regularization. The model learns characteristic spectral-temporal patterns of land surfaces without labelled or ancillary data. The resulting embeddings capture the temporal evolution of 10-band spectral signatures and SAR backscattering in VV and VH, and are precomputed as annual global rasters, accessible via the geotessera Python package^1^. For details, the reader is referred to Feng et al. (2025).

#### 2.3.2. Satellite composite baselines

##### Sentinel-2 (optical)

We used Sentinel-2 Level-2A surface reflectance imagery to generate annual and seasonal composite feature stacks for 2018. All scenes intersecting the study region were filtered to retain acquisitions with <60% scene-level cloud cover. Cloud/cirrus, cloud shadow, snow/ice, and no-data pixels were masked using the Sen2Cor Scene Classification Layer (SCL) and the QA60 cloud bitmask, and reflectance values were scaled to unitless surface reflectance (0–1) (see Gao et al., 2024).

To quantify the contribution of seasonal information in a conventional multi-sensor baseline, we produced two alternative Sentinel-2 representations: (i) an annual per-pixel median composite over all valid observations in the year, and (ii) seasonal composites computed as per-pixel medians within three phenologically meaningful periods—spring (March–May), summer (June–August), and autumn (September–November)—with winter excluded due to pervasive snow cover at higher elevations. Consistent with common practice in tree-species and forest type mapping, retaining seasonal structure aims to preserve phenological variation that is attenuated by annual aggregation.

All Sentinel-2 spectral bands (10 m, 20 m, 60 m; excluding atmospheric band B10) were retained. For each seasonal composite (and for the annual composite), we additionally computed NDVI, NDWI (McFeeters), and NBR.

##### Sentinel-1 (SAR)

To incorporate cloud-independent structural and moisture-sensitive information, we also derived Sentinel-1 GRD (IW mode) seasonal and annual features for 2018 using dual polarization VV and VH, along with the incidence angle band provided in the collection. The Earth Engine Sentinel-1 GRD collection is radiometrically calibrated and terrain corrected, with backscatter provided in decibels (dB). We formed seasonal (spring/summer/autumn) and annual Sentinel-1 composites by aggregating observations within the same date ranges as Sentinel-2. To reduce bias when compositing on a log scale, we computed medians in linear backscatter space and converted back to dB after aggregation (details in Section S.3). In addition to VV and VH backscatter, we derived simple polarization features (VV–VH in dB and VV/VH ratio) commonly used to capture vegetation structural differences.

##### Gridding and alignment

All composites were exported on a fixed 10 m UTM grid aligned exactly to the study region. Sentinel-2 bands at 20 m and 60 m were resampled to 10 m; Sentinel-1 features were exported on the same target grid. Full implementation details (mask classes, bitmasks, resampling choices, and export parameters) are provided in the Section S.3.

### 2.4. Label construction

Pixels within a forestry parcel were assigned the parcel’s dominant species label (determined from proportional inventory cover; see Fig. 1) for model training and testing. To improve the quality and quantity of the labeled pixels we applied some additional pre-processing steps (described below).

#### 2.4.1. Constraints of parcel-level labels

Forest inventory data in Trentino provide species composition at the level of management parcels rather than at the level of individual trees or pixels. Each parcel reports the proportional canopy cover of multiple species, often representing structurally and compositionally heterogeneous stands. As a result, direct use of parcel labels for pixel-level classification introduces several sources of ambiguity. Parcels frequently contain mixtures of species, parcel boundaries are irregular, and non-forest elements (e.g., roads, clearings, rocky outcrops) are not explicitly represented in the inventory.

Critically, parcel labels are not pixel-accurate ground truths. Treating them as such risks inflating apparent classification performance for dominant species while systematically penalising predictions of secondary species that are present but spatially intermixed. Conversely, discarding mixed parcels entirely would reduce the volume of usable training data and disproportionately exclude ecologically important stand types. Addressing this trade-off between label quantity and label contamination is central to enabling data-efficient species mapping from satellite imagery with wall-to-wall coverage using forest inventories as reference data. To address this issue, we propose a soft label training procedure described in S2.8.2.

#### 2.4.2. Land-cover masking

To ensure that species training and inference were restricted to ecologically plausible locations, we filtered non-forest areas during training label construction. Because inventory-derived species proportions refer exclusively to tree canopy cover (and exclude other ground cover such as roads or grassland), pixels corresponding to non-forest land-cover types (e.g. built surfaces, water bodies, bare ground, grassland, or persistent snow) were filtered to prevent implausible species assignments.

We used an independent, freely available, global land-cover product – the 10 m Global Land Cover 2020 (EU CLMS, 2020) – to identify forested pixels and gate species label assignment accordingly. Pixels classified as forest were retained as eligible for species inference, while non-forest pixels were assigned fixed non-tree classes to ensure consistent handling across all representations (see Section S.1.4 for the full filtering workflow).

### 2.5. Model selection

#### 2.5.1. Classifier architectures

We evaluated a small set of downstream classifier architectures to assess how classifier capacity influences the utility of different input representations. Our central premise is that geospatial foundation-model embeddings encode sufficiently rich spectral, structural, and phenological information to enable effective species discrimination without extensive fine-tuning or highly complex downstream models.

To probe this, we selected classifiers spanning a gradient of expressive capacity. Logistic Regression (LR; Hastie et al., 2009) provides a linear baseline, testing whether the embedding space is sufficiently structured for class separation using simple linear decision boundaries. K-Nearest Neighbours (kNN; (Cover and Hart, 1967) offers a non-parametric alternative that relies on local similarity in feature space without learning explicit decision boundaries; strong performance under kNN would indicate that ecologically similar species are clustered in the embedding space. Random Forests (RFs; (Breiman, 2001) serve as a widely adopted nonlinear baseline in ecological remote sensing, capable of modelling complex feature interactions through ensembles of axis-aligned decision trees. RFs are robust to noisy labels, mixed feature distributions, and class imbalance, and typically perform well with minimal tuning (Belgiu and Drăguţ, 2016), making them a practical reference point for applied mapping tasks. To assess whether additional nonlinear capacity yields meaningful performance gains beyond these baselines, we further evaluated multi-layer perceptrons (MLPs; Rumelhart et al., 1986). MLPs can learn smooth, high-dimensional nonlinear transformations of the input features, potentially exploiting structure in the embedding space that axis-aligned or instance-based methods cannot capture. Together, this set of classifiers defines a controlled gradient of downstream model capacity, enabling us to assess whether classification performance is primarily constrained by representation quality or by the expressive power of the classifier.

#### 2.5.2. Training and validation design

We employed a nested parcel-level cross-validation framework to tune hyperparameters while obtaining unbiased estimates of generalisation performance. The outer loop used five parcel-level folds to estimate test performance, while an inner loop tuned model hyperparameters using training data only. Hyperparameter optimisation was performed using Bayesian optimisation (Optuna; Akiba et al., 2019).

All experiments used a spatially explicit, parcel-aware partitioning strategy. Pixels belonging to the same forest inventory parcel were never split across training, validation, or test sets, preventing spatial leakage arising from within-parcel autocorrelation and providing a more realistic assessment of generalisation to unseen forest stands in the region. Preprocessing steps (including feature normalisation) were computed using training data only. After hyperparameter selection, models were retrained on the full outer-fold training set before evaluation on the held-out parcels. Performance is reported as the mean ± standard deviation across the five outer folds.

For all experiments, training parcels were restricted to those with a minimum dominant-species cover of 60% (aside from 2.8.2 where sensitivity to label noise was tested), while pixel-level evaluation metrics were computed on a higher-purity subset of parcels (≥80%) to ensure reliable reference labels. In contrast, the parcel-level species proportion error metric was computed across all validation parcels regardless of purity, reflecting performance in mixed-species stands (see Section 2.6). This spatial partitioning strategy, purity criteria, and evaluation protocol were held constant across all analyses, with only the input representation or classifier class varying between experiments.

To ensure reproducibility in downstream experiments (Sections 2.8-2.9), we fixed hyperparameters using the configuration from an outer fold that achieved optimal tuning performance while exhibiting representative architectural choices across folds.

#### 2.5.3. Model training and hyperparameter tuning

Hyperparameters for all models were selected using a composite validation metric designed to balance pixel-level classification accuracy with parcel-level compositional realism. Specifically, models were tuned to minimise the following objective:

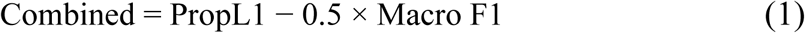

where PropL1 denotes the parcel-level species proportion error (Section 2.6.2) and Macro F1 denotes the macro F1 score (pixel level; see Section 2.6.3). Lower values of the combined metric indicate better overall performance.

This objective penalises models that achieve high pixel-level accuracy at the expense of unrealistic parcel-level species composition, while also avoiding solutions that recover aggregate species proportions but misclassify individual pixels. The weighting factor of 0.5 reflects a deliberate trade-off, such that a 1% improvement in Macro F1 offsets a 0.5% increase in parcel-level proportion error and was fixed across all experiments. This formulation encourages models that can detect less common species while predicting species in the correct proportions at the aggregate scale. This composite objective was used exclusively for hyperparameter selection and does not affect the evaluation metrics reported in Section 2.6.

#### 2.5.4. Class imbalance handling

The area covered by the different species is highly unbalanced (see Table 1). To mitigate severe class imbalance during model training, we applied parcel-aware downsampling of the most abundant species classes. Specifically, the three most common species by pixel count were downsampled within each training fold to match the pixel count of the fourth most abundant species (Fig. S.2).

Downsampling was performed in a parcel-stratified manner, such that retained pixels for each species were distributed across as many parcels as possible. This strategy preserves spatial and environmental variability within dominant species classes while reducing their numerical dominance during optimisation. Downsampling was applied during training only and did not affect validation or test data. The same class-balancing procedure was used across all experiments to ensure consistent and comparable optimisation.

### 2.6. Performance metrics and interpretation

We evaluated classification performance at both the pixel and parcel levels to capture complementary aspects of model behaviour relevant to ecological interpretation and forest management applications.

#### 2.6.1. Pixel-level classification metrics

To characterise pixel-level classification performance and the structure of model errors, we analysed class-level confusion matrices and F1-based metrics. Classification performance at the pixel level was assessed using the F1 score, defined as the harmonic mean of precision and recall for each class. These were assessed using parcels of a minimum 80% purity (cover of dominant class). To provide complementary perspectives on model performance, we computed both weighted and macro-averaged F1 scores across all species classes.

**Confusion matrices** report the frequency with which pixels belonging to a given reference species are assigned to each predicted species. They illustrate systematic misclassification patterns, including whether errors occur predominantly within functional or phylogenetic groups.

**Macro F1 (MF1)** computes the unweighted arithmetic mean of per-class F1 scores, treating all species equally regardless of their spatial prevalence. This metric is informative for evaluating model performance on rare or minority species, which may be of high ecological or conservation importance but contribute minimally to weighted metrics. A model achieving high macro F1 must perform well across the full range of species, including those with limited training samples.

**Weighted F1 (WF1)** computes the mean F1 score across classes, weighted by the number of pixels belonging to each class (i.e., class prevalence). This metric reflects overall classification accuracy and is dominated by performance on abundant species, making it sensitive to errors in the dominant land cover types that constitute the majority of the study area.

While WF1 reflects overall classification accuracy, MF1 is more sensitive to performance differences among less common species and was therefore emphasised in model tuning and when interpreting model behaviour.

#### 2.6.2. Parcel-level species proportion error

To evaluate model performance at a management-relevant spatial scale and on the full range of mixed parcels available, we quantified errors in predicted species composition at the parcel level using a parcel-level species proportion error metric (PropL1), defined as the Total Variation Distance between predicted and reference species-proportion distributions and mathematically equivalent to Bray–Curtis dissimilarity for proportional data (Gibbs and Su, 2002; Bray and Curtis, 1957). For each parcel, pixel-level predictions were aggregated to form a predicted species proportion vector, which was compared to inventory-derived reference proportions.

PropL1 is defined as:

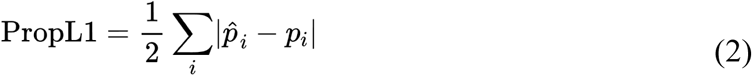

where ***p̂*_*i*_** and ***p_i_*** denote predicted and reference proportions for species *i*, respectively. This metric ranges from 0 (perfect agreement) to 1 (complete mismatch) and was computed across all validation parcels regardless of purity. For example, a PropL1 value of 0.3 corresponds to an average mismatch of 30% in parcel-level species composition, meaning that 30% of parcel area would need to be reassigned across species to match the reference distribution.

Unlike pixel-level accuracy metrics, which require assumptions about parcel purity and are therefore restricted to near-monospecific stands, PropL1 makes no assumptions about stand composition and does not require filtering or thresholding of the validation data. As a result, it provides a more robust and comprehensive assessment of model performance in heterogeneous landscapes.

### 2.7. Effect of downstream classifier capacity

To assess the extent to which classification performance is constrained by the quality of the input representations versus the capacity of the downstream model, we analyse how predictive performance varies across classifiers of increasing expressive power, while holding the training protocol, spatial cross-validation design, class-balancing strategy, and evaluation metrics fixed (Section 2.5).

For each input representation, we compare performance across linear, instance-based, tree-based, and neural network classifiers to determine whether discriminative information encoded in the representations is readily accessible to simple decision functions, or whether additional nonlinear capacity is required to exploit more complex structure in the embedding space. This analysis allows us to distinguish representation-limited from classifier-limited regimes and to assess whether gains from increased model capacity are consistent across representations or concentrated in specific cases.

Unless otherwise stated, all results in subsequent sections use the best-performing multi-layer perceptron configuration identified in this comparison, providing a consistent and competitive downstream model for representation-focused analyses.

### 2.8. Analyses of representation quality and supervision sensitivity

#### 2.8.1. Label availability and fraction-curve analysis

To quantify label efficiency, we conducted a fraction-curve analysis examining how classification performance scales with the amount of available training data. Models were trained using increasing fractions of the available training parcels, ranging from extremely limited supervision (0.1%) to near-complete data availability, while holding the evaluation protocol fixed.

Training parcels were selected using stratified random sampling at the parcel level to preserve class balance. Hyperparameters were fixed across all fractions, using values selected via nested cross-validation on the full training dataset (Section 2.5), such that fraction curves isolate the effect of label availability rather than changes in model configuration. At very low training fractions, class-balanced training sets were enforced to prevent dominance by the most abundant species.

At each fraction, models were trained and evaluated using 5-fold parcel-aware cross-validation repeated across three random seeds. Performance was assessed using both pixel- and parcel-level metrics (Section 2.6). Fraction curves were computed separately for foundation-model embeddings and satellite composites, enabling direct comparison of how rapidly performance saturates as labelled data increase.

#### 2.8.2. Sensitivity to label purity and weak supervision

To assess sensitivity to training-label uncertainty, we varied the minimum dominant-species cover threshold (purity) required for parcels included in model training. This analysis captures the trade-off between label noise and training data volume inherent in parcel-based forest inventories, where many stands contain mixed species compositions. For each input representation, models were trained using parcels exceeding a given purity threshold (30% dominance, ∼83k parcels, to 100% dominance, ∼4.9k parcels), while holding the overall training fraction constant. Pixel level validation was always performed on a high-purity subset of parcels (≥80% dominant species), independent of the training purity threshold. Parcel level (PropL1) was always performed across all purity thresholds.

To test whether parcel-level species composition information can mitigate label noise, we evaluated a soft-label training formulation in which pixel-level targets are defined by parcel-level species proportions. Under this formulation, each pixel is assigned a probabilistic target vector whose entries correspond to the fractional species composition of the parent parcel. This approach explicitly represents the uncertainty in pixel-level class membership induced by mixed-species parcels and enables weak supervision to be incorporated directly during model training. Neural network models were trained using a cross-entropy loss generalised to soft targets, encouraging predicted class probabilities to reflect observed species mixtures rather than enforcing hard assignments. Performance under soft-label training was compared to standard dominant-species labelling across the same purity thresholds, allowing us to assess whether weak, proportion-based supervision improves robustness to noisy training labels. An alternative strategy to refine input data based on label proportions was evaluated but not included here as it did not yield consistent improvements (see S.4).

#### 2.8.3. Ancillary environmental covariates

To evaluate whether explicitly providing terrain-derived environmental context improves classification performance beyond that encoded in the remote-sensing representations, we augmented the input feature space with terrain-derived covariates. Elevation data were obtained from the Shuttle Radar Topography Mission (SRTM) digital elevation model (30 m resolution; (Farr et al., 2007) and resampled to the satellite imagery grid.

From the resampled DEM, we derived elevation, slope, aspect, and topographic position index (TPI; Wilson and Gallant, 2000). Aspect was encoded using sine and cosine transformations to avoid angular discontinuities, resulting in five terrain-related input channels in total. Terrain covariates were concatenated with the corresponding remote-sensing features prior to model training.

All input features were standardised using z-scoring based on statistics computed from the training data only. Models incorporating terrain covariates were tuned using the same hyperparameter optimisation procedure and composite validation objective described in Section 2.5, ensuring that any performance differences reflect the contribution of the additional inputs rather than differences in model optimisation.

### 2.9. Temporal transfer experiments

To assess the temporal generalisability of the different input representations, we conducted a cross-year transfer experiment in which models trained on data from one year were applied to a subsequent year without retraining. This analysis evaluates the stability of learned representations under interannual variation in phenology and acquisition conditions.

Models were trained using embeddings from 2018 and evaluated on parcels from both 2018 (within-year validation) and 2019 (cross-year transfer). Based on the classifier capacity analysis in Section 2.7, we used the shallow multi-layer perceptron configurations identified as appropriate for exploiting the representations. Parcel assignments to training and evaluation folds were fixed across years, such that a given parcel was never used for training in one year and evaluation in another, preventing spatial overlap when assessing temporal transfer. The same parcel-aware partitioning strategy and purity thresholds described in Section 2.5 were applied, with pixel-level evaluation metrics computed on high-purity parcels to minimise label uncertainty.

No additional fine-tuning or domain adaptation was performed when transferring models across years. This design isolates the effect of temporal domain shift and provides a conservative assessment of how well classification boundaries learned on representations in one year transfer to subsequent observations. Performance was evaluated using the same pixel-level and parcel-level metrics as in the within-year analyses, enabling direct comparison of temporal and non-temporal generalisation behaviour.

This temporal transfer experiment serves as a stress test of representation robustness rather than a demonstration of optimal change detection and/or multi-year modelling strategies. As such, it highlights the extent to which current geospatial foundation-model embeddings encode temporally stable species-discriminative information under realistic monitoring conditions.

## 3. Results

### 3.1. Classification accuracy, label efficiency, and ecological structure

Across experiments, geospatial foundation-model embeddings outperformed conventional Sentinel-1+2 spectral composites for species-level tree classification. Using the best-performing MLP configuration for each representation, Tessera and AlphaEarth achieved similar overall accuracy (weighted F1 = 0.827 ± 0.010 and 0.833 ± 0.007, respectively) and higher macro F1 than Sentinel-1+2 baselines, indicating improved discrimination of less dominant species (Table 3). Sentinel-1+2 annual composites performed substantially worse across all metrics, consistent with temporally aggregated spectral features being insufficient for fine-grained species discrimination in heterogeneous mountain forests.

**Table 3.**
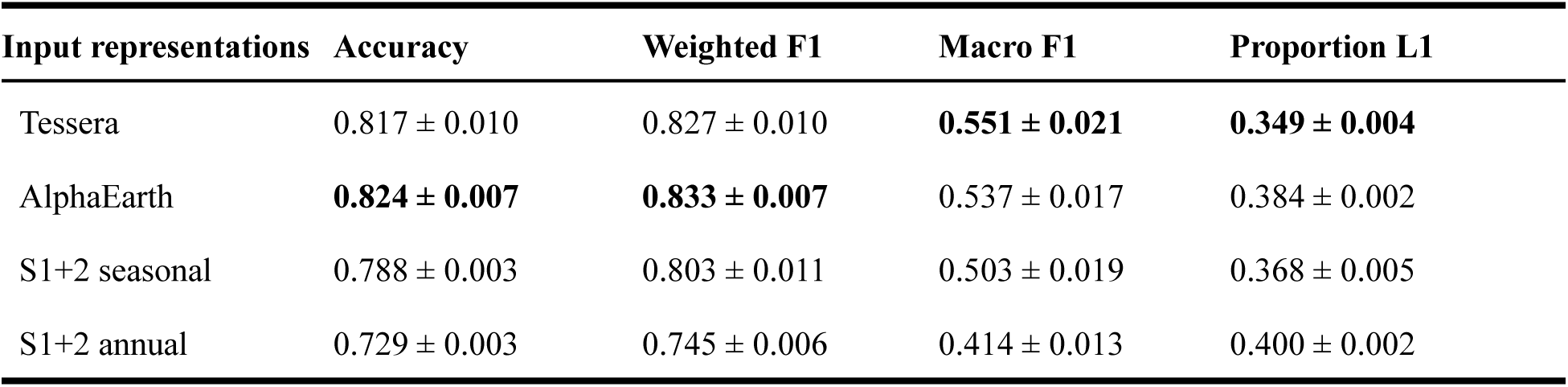
Classification performance of MLP models trained on different input representations, evaluated using 5-fold nested cross-validation.

| Input representations | Accuracy | Weighted F1 | Macro F1 | Proportion L1 |
| --- | --- | --- | --- | --- |
| Tessera | $0.817 \pm 0.010$ | $0.827 \pm 0.010$ | **$0.551 \pm 0.021$** | **$0.349 \pm 0.004$** |
| AlphaEarth | **$0.824 \pm 0.007$** | **$0.833 \pm 0.007$** | $0.537 \pm 0.017$ | $0.384 \pm 0.002$ |
| S1+2 seasonal | $0.788 \pm 0.003$ | $0.803 \pm 0.011$ | $0.503 \pm 0.019$ | $0.368 \pm 0.005$ |
| S1+2 annual | $0.729 \pm 0.003$ | $0.745 \pm 0.006$ | $0.414 \pm 0.013$ | $0.400 \pm 0.002$ |

#### Label efficiency

Figure 4 shows how performance scales with the fraction of labelled training parcels (100% corresponds to ∼43,834 parcels on average across folds). For weighted F1, both foundation-model embeddings exhibit strong label efficiency, reaching near-asymptotic performance with ≤5% of training parcels and converging rapidly as supervision increases. Tessera shows a small but consistent advantage at the lowest training fractions, while differences between Tessera and AlphaEarth diminish with additional training data. Sentinel-1+2 baselines lag behind the foundation models across all training fractions, with annual composites consistently underperforming seasonal composites.

**Figure 4.**
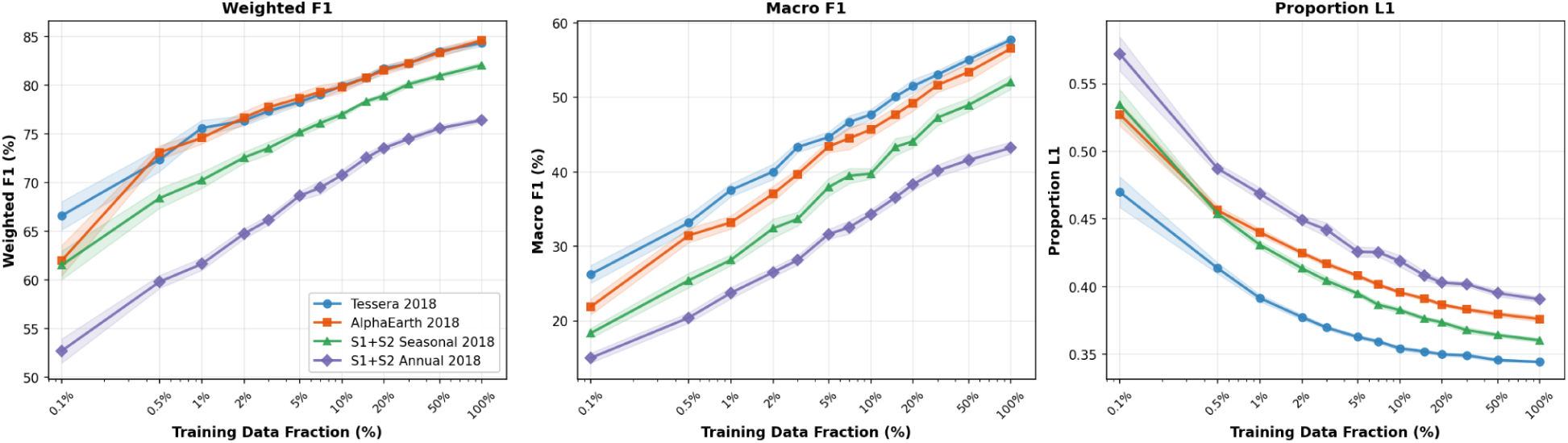
Label efficiency curves comparing Tessera 2018, AlphaEarth 2018 and Sentinel-1+2 seasonal and annual composites feature representations for multi-label forest species classification. Performance metrics are shown as functions of training data fraction for a multi-layer perceptron (MLP) classifier trained on forest parcel data from Trentino, Italy. (Left) Weighted F1-score, which accounts for class imbalance by weighting each class by its support in the test set. (Middle) Macro F1-score, computed as the unweighted mean of per-class F1-scores, providing equal importance to all species regardless of frequency. (Right) Proportion L1 error, measuring the mean absolute difference between predicted and true class proportions across the landscape (lower values indicate better preservation of species distributions). Shaded regions indicate 95% confidence intervals. Here, 100% of training data is ∼43,834 training parcels (or ∼5.9M training pixels).

Performance differences are more pronounced for macro F1, which is more sensitive to minority-class performance. Macro F1 increases more gradually with training fraction, indicating that continued gains primarily reflect improved discrimination of rarer species. Tessera achieves the highest macro F1 across most fractions, followed closely by AlphaEarth, while both Sentinel-1+2 variants show substantially weaker and more slowly saturating performance.

#### Parcel-level composition fidelity

Patterns in PropL1 differ from those observed for F1-based metrics. Tessera achieves the lowest proportion error across all training fractions, indicating the strongest preservation of parcel-level species composition. Notably, Sentinel-1+2 seasonal composites consistently outperform AlphaEarth on PropL1 despite lower pixel-level F1 scores, indicating that representations can differ in how well they preserve aggregate class frequencies even when their pixel-level discrimination differs. Overall, Tessera performs strongly on both discriminative accuracy and composition fidelity, whereas AlphaEarth prioritises discriminative accuracy and Sentinel-1+2 seasonal composites show relatively improved proportion calibration. Figure 5 shows pixel-level species predictions for three mixed validation parcels showing how predicted proportions correspond to ground truth proportions and the resulting PropL1 score.

**Figure 5.**
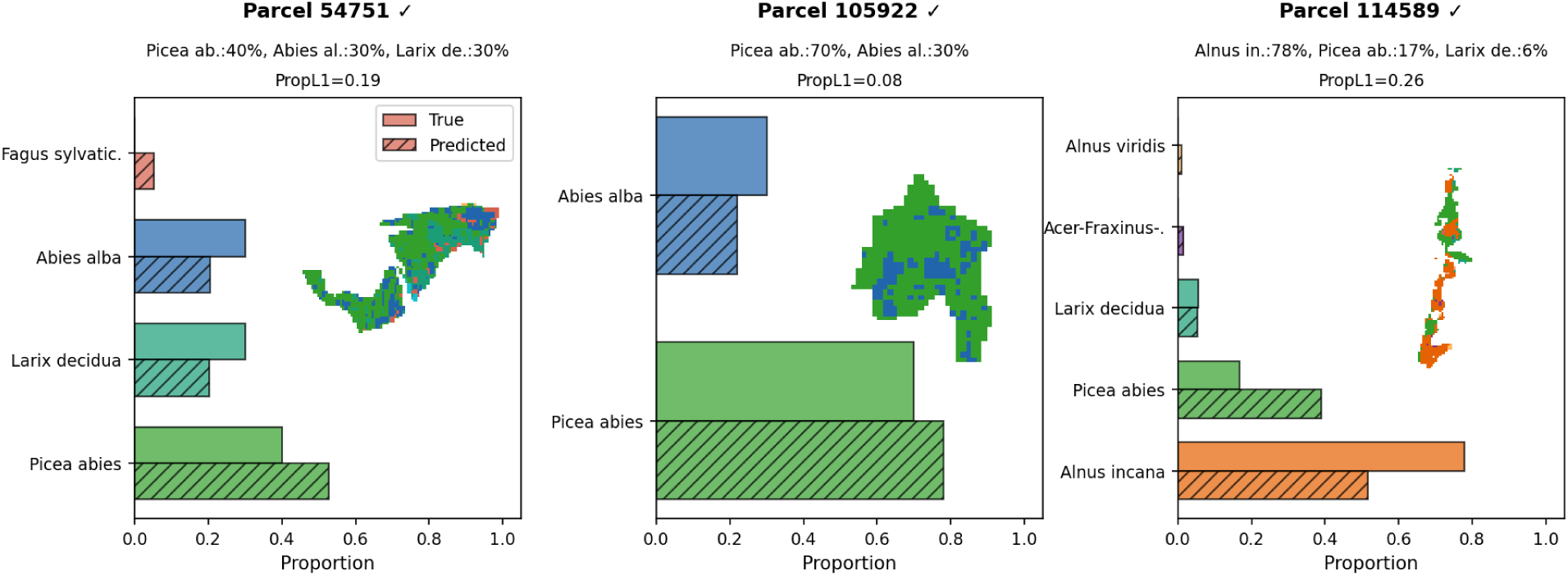
Pixel-level species predictions for three mixed validation parcels using the MLP classifier with Tessera embeddings. The bar chart compares true species proportions (solid bars) against predicted proportions aggregated from pixel classifications (hatched bars). For each parcel, the spatial prediction map (inset) shows individual pixel classifications colored by species. The Proportion L1 score (PropL1) quantifies the total variation distance between true and predicted proportions, ranging from 0 (perfect match) to 1 (complete mismatch). The tick (✓) or cross (✗) indicates whether the model correctly identified the dominant species at the parcel level. Species colours are consistent across maps and bars.

#### Embedding-space structure

To examine the ecological structure underlying these performance gains, we visualised Tessera embeddings in three dimensions using UMAP (Figure 6). The low-dimensional manifold appears to be organised primarily along broad environmental gradients. Elevation emerges as a dominant organising axis spanning both coniferous and broadleaf taxa, while separation by functional type is clearly evident. At finer taxonomic scales, genera and species occupy consistent regions of the manifold, with within-genus separation aligned with known ecological and elevational niches. Ecologically broad genera (e.g. *Pinus*) show broader dispersion, consistent with wide environmental amplitude.

**Figure 6:**
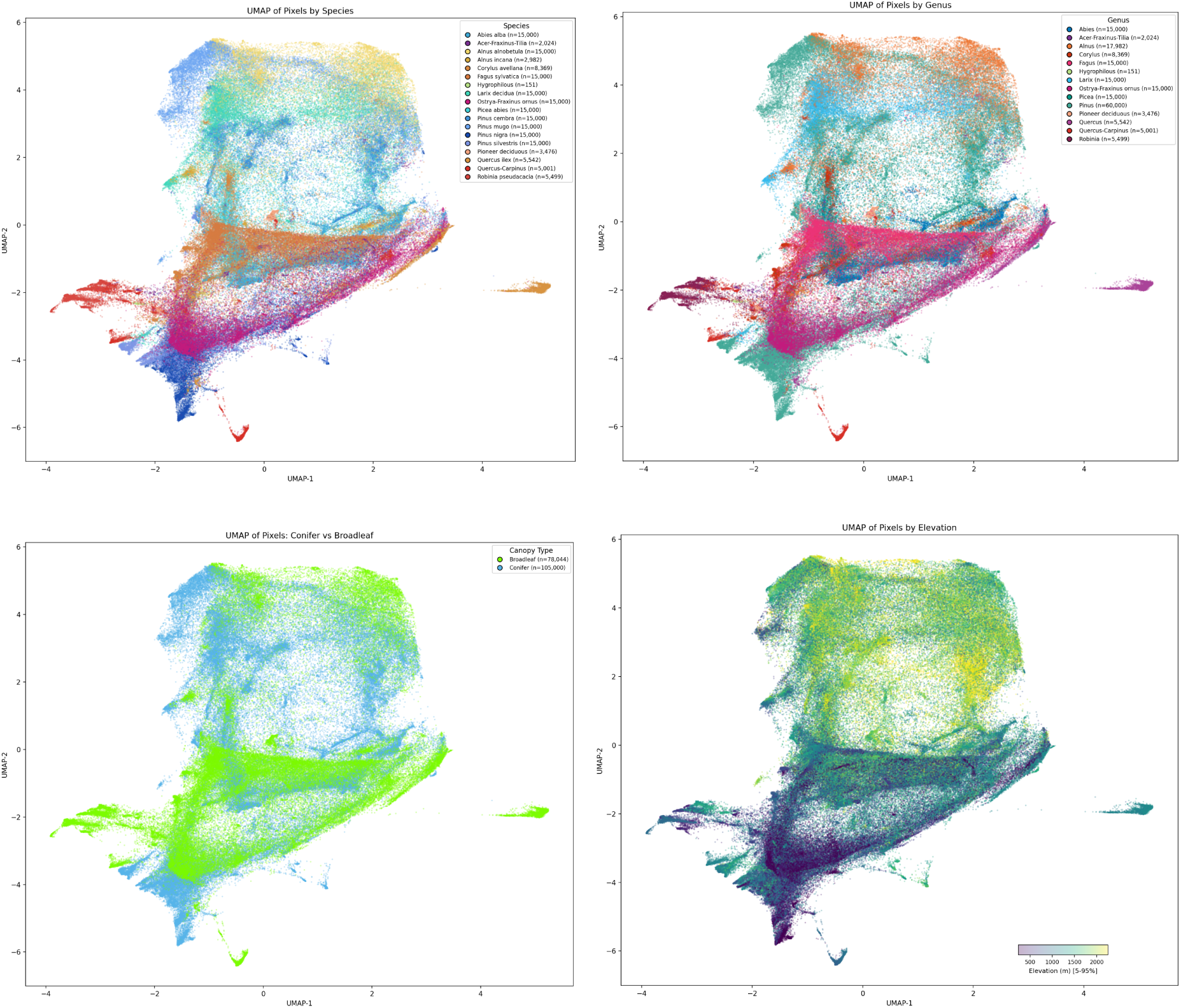
UMAP representations of Tessera embeddings coloured by species (or species groups), genus, broadleaf/conifer functional types and elevation. Point ordering done by the 3rd UMAP dimension to give the effect of a z-axis and demonstrate separation in this 3rd dimension. A plot with each species group plotted separately is available in Fig S.20. UMAP projections coloured by individual species (pre-merge) are provided in Figs S.13–S.15.

#### Error structure

The row-normalised confusion matrix for the Tessera + MLP model exhibits a coherent block-diagonal structure (Figure 7), indicating strong separability between classes. Several taxa show high recall, including Norway spruce (*Picea abies*; 0.93), European beech (*Fagus sylvatica*; 0.85), Scots pine (*Pinus sylvestris*; 0.82), mountain pine (*Pinus mugo*; 0.81), and European larch (*Larix decidua*; 0.79). Misclassifications are concentrated within functional groups and within genera: silver fir (*Abies alba*) is frequently confused with Norway spruce (*Picea abies*; 0.57), and moderate confusion occurs among pine species (e.g. *P. nigra*, *P. sylvestris*, *P. cembra*). Broadleaf taxa show more heterogeneous performance; European beech (*Fagus sylvatica*) is among the best-classified deciduous species, whereas sweet chestnut (*Castanea sativa*) and common hazel (*Corylus avellana*) show lower recall with predictions dispersed across other deciduous and mixed classes (largely aligned with the availability of training data). Mixed broadleaf assemblages (e.g. oak–hornbeam, Quercus–Carpinus; hop-hornbeam–manna ash, Ostrya–Fraxinus ornus) are frequently confused with one another. This may partly reflect genuine ecological and spectral overlap among co-occurring deciduous taxa, but also the internal heterogeneity introduced by merging spectrally distinct species into composite classes – a pragmatic choice that increases sample size at the cost of class coherence. The weak classification of hygrophilious and confusion with *Pinus silvestris* is likely a result of very few training examples. Overall, errors are strongly structured by phylogenetic relatedness, functional type, and stand composition, consistent with embeddings encoding ecologically meaningful variation even where species-level discrimination is challenging.

**Figure 7:**
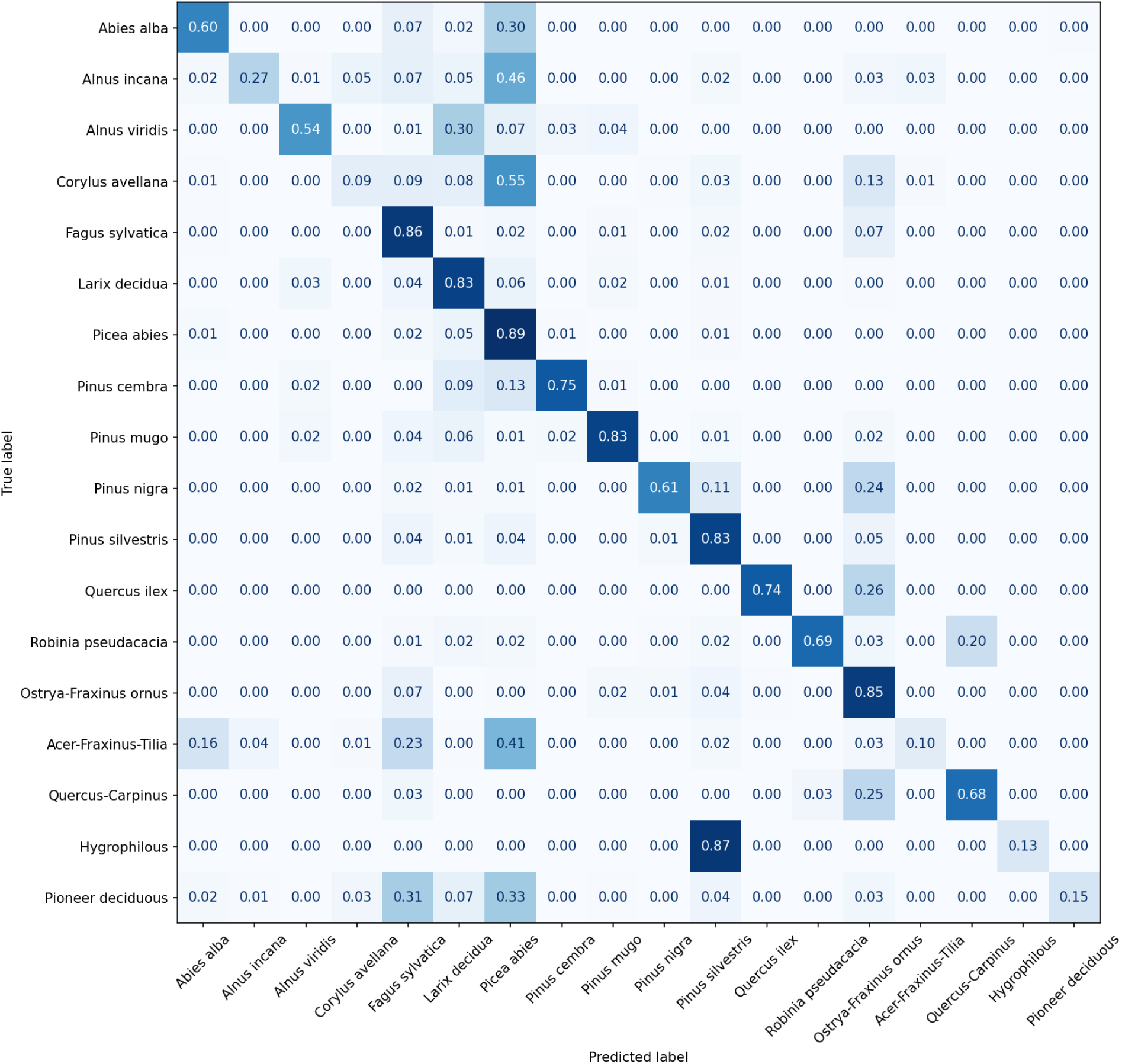
Row-normalised confusion matrix for pixel-level species classification on validation parcels using MLP training on Tessera embeddings. Eighteen species and merged species groups are shown.

**Figure 8:**
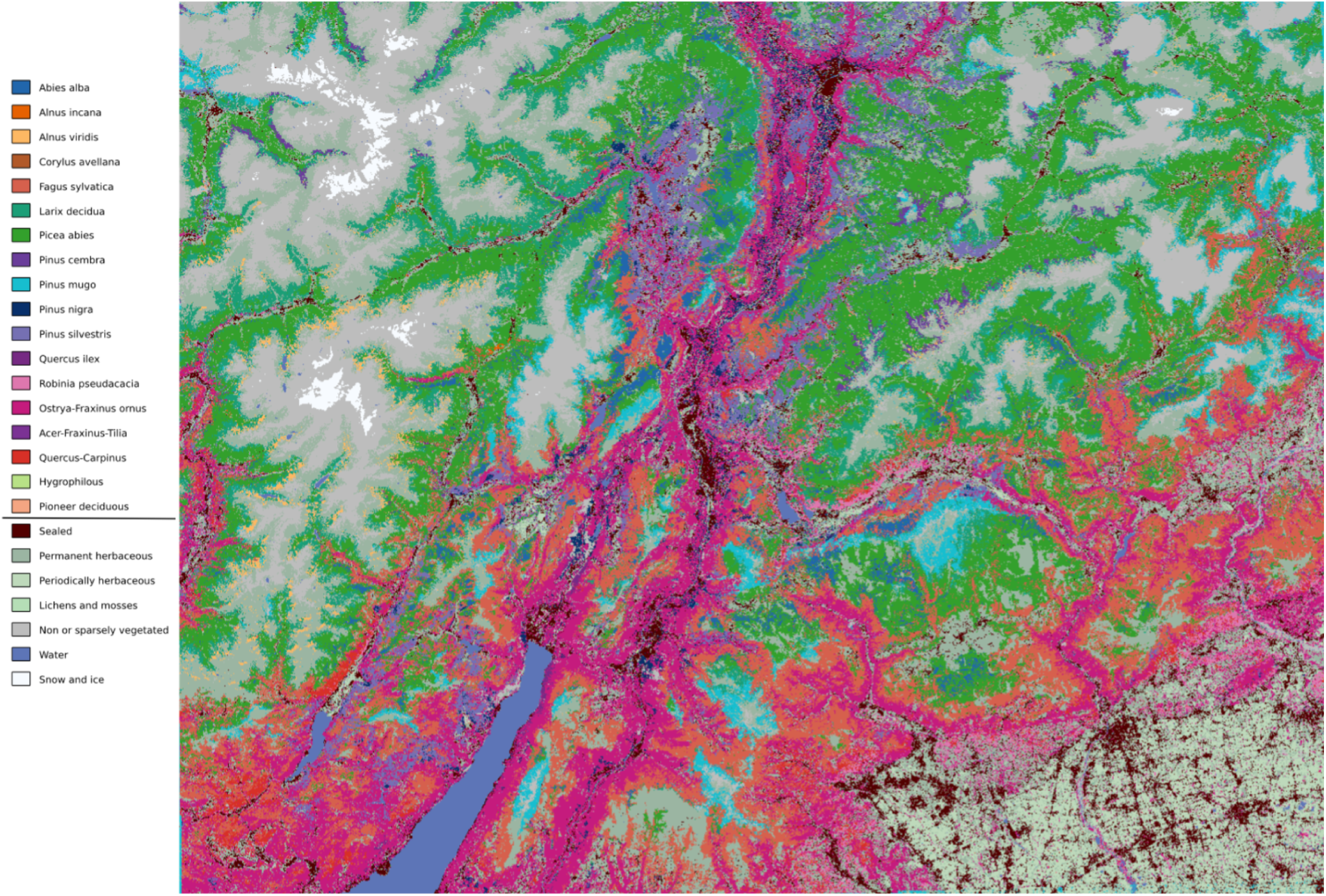
Wall-to-wall species composition map of Trentino at 10 m resolution. Forest pixels are classified into 18 tree species and species groups by an MLP trained with soft labels (KL-divergence loss) on Tessera foundation-model embeddings, using provincial forest-inventory polygons as reference data. Non-forest land cover (sealed surfaces, herbaceous, bare ground, water, snow/ice) is taken directly from the Copernicus Land Monitoring Service CLC+ Backbone 2018 product. Full raster predictions are given as a supplementary file.

### 3.2. Effect of downstream classifier capacity

To assess the sensitivity of classification performance to downstream model capacity, we evaluated a set of commonly used classifiers spanning increasing expressive power (Figure 9; full numerical results in Table S.8 and Fig S.10). Both classifier choice and input representation affected performance, with their relative importance depending on the metric considered.

**Figure 9.**
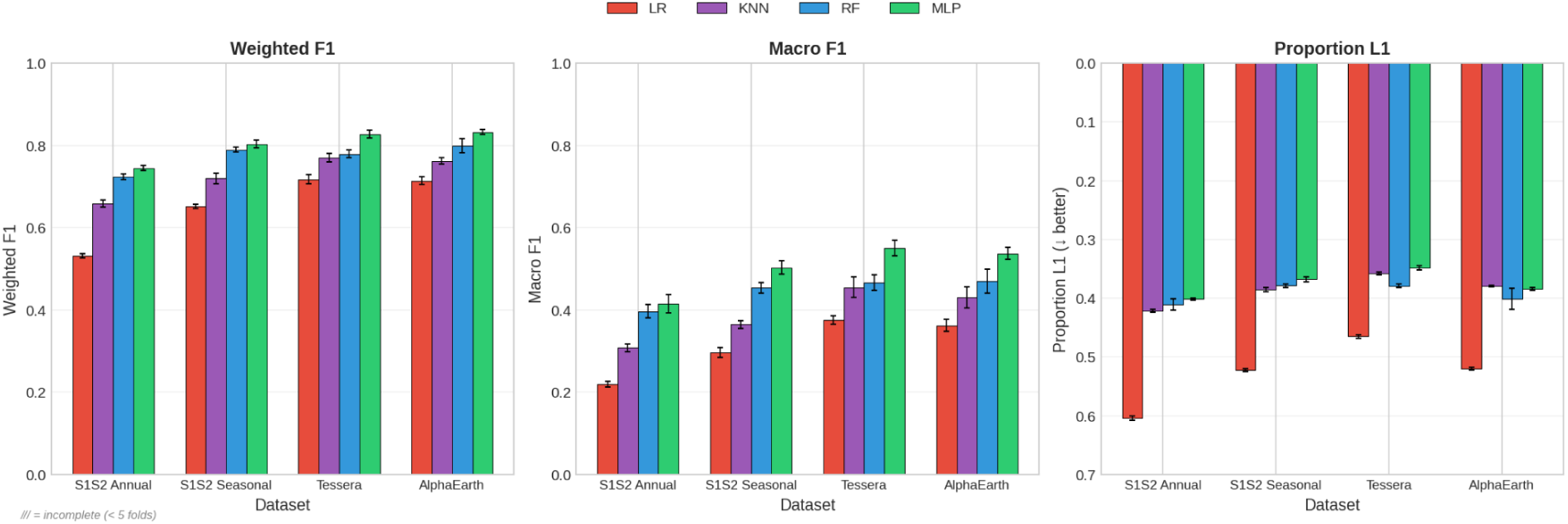
Comparison of classification performance across four model architectures and input feature sets using 5-fold nested cross-validation with class-balanced training data. Metrics shown are (a) Weighted F1-score, (b) Macro F1-score, and (c) Proportion L1 error (a lower score is better which is why the y-axis is inverted). Models compared: Logistic Regression (LR), K-Nearest Neighbors (KNN), Random Forest (RF), and Multi-Layer Perceptron (MLP). Input features: Sentinel-1/2 annual composite, Sentinel-1/2 seasonal composite, Tessera, and AlphaEarth embeddings. Error bars indicate standard deviation across folds.

For all input representations, MLPs achieved the highest performance, followed by Random Forest and k-nearest neighbour classifiers, while linear models performed considerably worse. Using Tessera embeddings, macro F1 increased from 0.38 (LR) to 0.55 (MLP), a gain of +0.17; weighted F1 increased from 0.72 to 0.83 (+0.11). AlphaEarth embeddings showed comparable gains. The effect of switching input representation while holding the classifier fixed was smaller for macro F1: with an MLP, moving from S1+2 seasonal composites to Tessera improved macro F1 by +0.05 and weighted F1 by +0.02. Thus, for macro F1 – the metric most sensitive to minority-species discrimination – classifier choice had a larger effect than representation choice. A linear classifier applied to foundation-model embeddings performed below a seasonal S1+2 composite paired with an MLP.

The practical consequence is that nonlinear classifiers are needed to unlock the discriminative information encoded in foundation-model embeddings. However, once a nonlinear classifier is used, further increases in capacity yield diminishing returns: the gain from RF to MLP (+0.08 macro F1 on Tessera) was modest, and hyperparameter optimisation typically favoured compact neural architectures – shallow MLPs (one or two hidden layers) with strong regularisation. Increasing network depth or complexity beyond this did not yield systematic performance gains. Performance gains from increased classifier capacity were most pronounced for macro F1, indicating improved discrimination of rarer species, whereas weighted F1 and parcel-level proportion error showed smaller relative improvements.

Together, these results indicate that both representation quality and classifier capacity contribute to species classification performance. Foundation-model embeddings provide a higher performance ceiling, but realising that ceiling requires at least a moderately nonlinear classifier. Once that threshold is met, further gains from increased model complexity are bounded, and simple, computationally efficient per-pixel classifiers are sufficient.

### 3.3. Do environmental covariates help?

We tested whether supplementing foundation-model embeddings with terrain-derived covariates improves species classification performance. Across all evaluated metrics, the inclusion of terrain-derived covariates produced no meaningful performance gains relative to using embeddings alone (Tables 4-5). For both Tessera and AlphaEarth embeddings trained on 2018 data, weighted F1, macro F1, and parcel-level proportion error (PropL1) remained effectively unchanged, with differences consistently <0.5% across metrics.

**Table 4:** Performance results for MLP classifier with Tessera 2018 embeddings and terrain-derived covariates.

| Metric | Embeddings Only | +Geofeatures (DEM) | Difference |
| --- | --- | --- | --- |
| Accuracy | 81.70% | 81.73% | +0.03% |
| Weighted F1 | 82.72% | 82.74% | +0.02% |
| Macro F1 | 55.10% | 54.76% | -0.34% |
| Prop. L1 | 0.349 | 0.350 | +0.002 |

**Table 5:** Performance results for MLP classifier with AlphaEarth 2018 embeddings and terrain-derived covariates.

| **Metric** | **Embeddings Only** | **+Geofeatures (DEM)** | **Difference** |
| --- | --- | --- | --- |
| Accuracy | 82.36% | 82.15% | -0.21% |
| Weighted F1 | 83.25% | 83.15% | -0.10% |
| Macro F1 | 53.75% | 53.98% | +0.23% |
| Prop. L1 | 0.384 | 0.384 | -0.000 |

Across both representations, adding terrain-derived covariates neither improved overall classification accuracy nor systematically benefited minority species or parcel-level composition estimates. Performance differences were small relative to cross-validation variability, indicating that any additional information provided by these covariates was not exploited by the downstream models.

### 3.4. Sensitivity to label purity and weak supervision

We evaluated sensitivity to training-label purity by varying the minimum dominant-species threshold required for parcels included in training, while holding the overall training fraction constant (Figure 10; Tables S.10–S.11). Pixel-level classification performance was comparatively robust across a broad range of purity thresholds. Weighted F1 increased slightly from low to intermediate purities (peaking around ∼70–80%) and then declined sharply when training was restricted to near-monocultures (≥90–100%), consistent with a reduction in available training data. Macro F1 followed a similar pattern but showed greater sensitivity to high-purity filtering, indicating that aggressively excluding mixed parcels disproportionately degrades performance for less common species.

**Figure 10.**
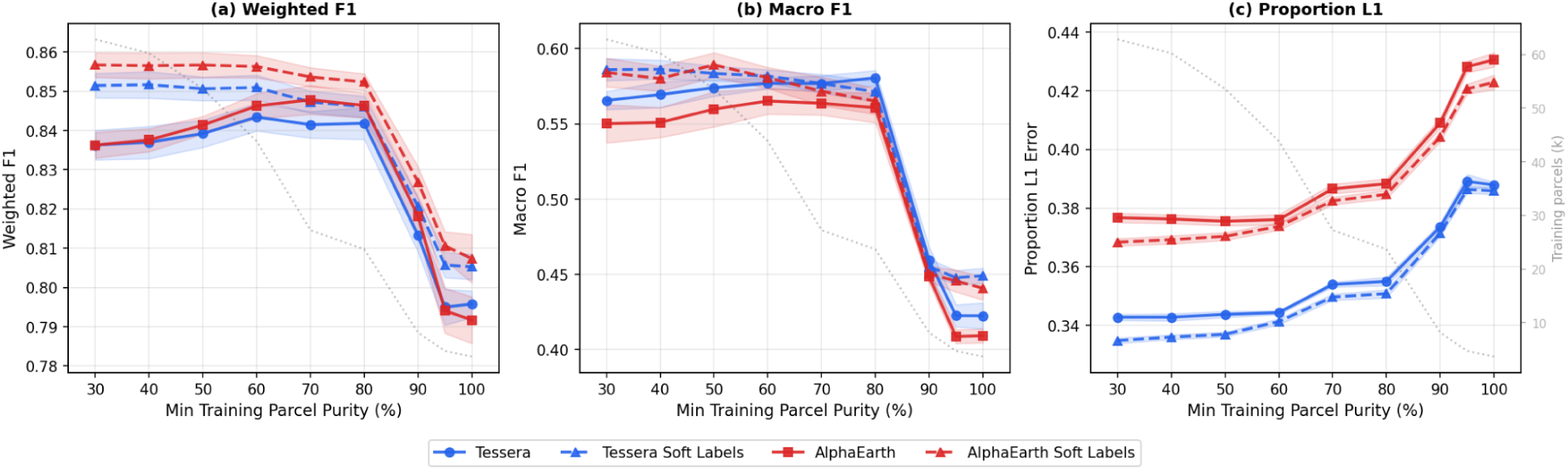
Effect of training parcel purity threshold on MLP classification performance. Comparison of model performance when training data is restricted to parcels exceeding progressively higher purity thresholds, evaluated on the Tessera and AlphaEarth label sources. Solid lines denote hard-label training (argmax class assignment); dashed lines denote soft-label training (proportion targets). (a) Weighted F1 score, measuring overall pixel-level classification accuracy. (b) Macro F1 score, the unweighted average across classes, reflecting performance on rare land cover types. (c) Proportion L1 error, measuring the absolute difference between predicted and true class proportions aggregated across the study area. Shaded ribbons indicate 95% confidence intervals from 5-fold cross-validation repeated with 3 random seeds. Pixel-level validation (weighted and macro F1) is performed exclusively on parcels with ≥80% purity, whereas parcel-level validation (Proportion L1) is performed across parcels of all purity levels. The dotted grey line shows the number of available training parcels (right axis) at each purity threshold.

Parcel-level composition fidelity exhibited the opposite dependence. Proportion error (PropL1) was lowest when mixed parcels were retained and increased steadily as stricter purity thresholds were applied, rising markedly at ≥90%. Thus, while high-purity filtering may marginally improve dominant-class assignment, it degrades the ability to recover realistic within-parcel species composition.

To test whether aggregate parcel-level species proportions could be leveraged, we trained MLP classifiers using soft labels in which each pixel received the full parcel-level species composition vector as its training target, optimised with KL-divergence loss (see Section S.6). Evaluated at 30% purity (the lowest threshold, maximising training volume), soft-label training achieved peak macro F1 of 0.586 for Tessera and 0.589 for AlphaEarth, exceeding the respective hard-label peaks of 0.581 and 0.565 obtained at 80% and 60% purity. In both cases, soft labels reached their best performance at low purity thresholds (30–50%) using all available training data, while hard labels required filtering to high-purity parcels. Proportion L1 error — measuring the fidelity of predicted species proportions at the landscape scale — improved consistently under soft-label training, decreasing by 0.006–0.008 across the stable purity range for both embeddings (e.g. 0.335 vs. 0.343 for Tessera and 0.368 vs. 0.377 for AlphaEarth at 30% purity). These gains were disproportionately concentrated in less common species.

Across the purity curve, the soft-label advantage was largest at low purity thresholds where mixed parcels dominate, and narrowed at intermediate purity (70–80%) where dominant-species labels are already largely correct (Figure 10). The pattern was consistent across both foundation models, with AlphaEarth showing a larger macro F1 advantage (+0.03 at 30% purity) than Tessera (+0.02), suggesting that soft labels may partially compensate for weaker embedding-level species discrimination. At extreme purity thresholds (95–100%), where training data becomes severely limited, soft labels again outperformed hard labels for both embeddings, suggesting that distributional supervision degrades more gracefully under data scarcity. Proportion L1 error showed a consistent advantage for soft labels across the full purity range for both Tessera and AlphaEarth, reflecting improved recovery of landscape-scale species proportions.

Taken together, these results show that foundation-model embeddings are tolerant to moderate label impurity in dominant-species training labels and that retaining mixed parcels does not substantially penalise pixel-level classification performance. Soft-label training provides targeted benefits for improving discrimination of less common species and, to a lesser extent, for recovering parcel-level species composition in heterogeneous stands.

### 3.5. Temporal transferability

To assess temporal generalisation, we trained MLP classifiers on 2018 embeddings and evaluated performance on both held-out 2018 parcels (within-year validation) and parcels from 2019 (cross-year transfer), using identical parcel-level partitions and high-purity validation criteria in both cases (Table 6; Figure 11; see also Figs S.21-S.22). This design isolates temporal domain shift while preventing spatial leakage between years.

**Fig 11:**
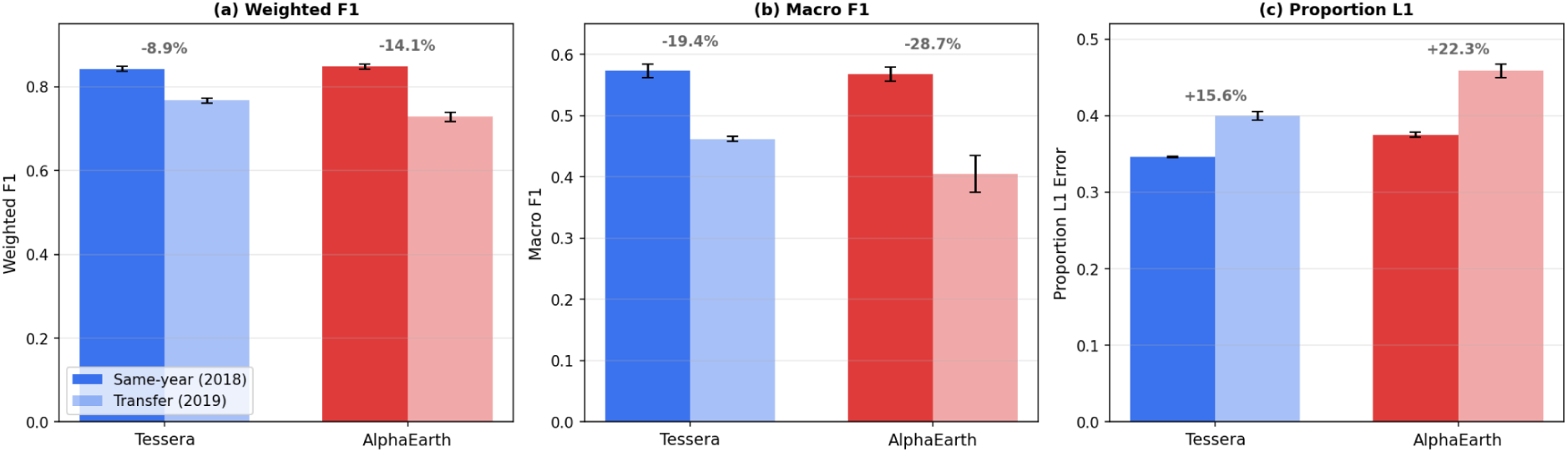
Temporal transfer performance for Tessera and AlphaEarth embeddings. MLP classifiers trained on 2018 data were evaluated on same-year holdout parcels (dark bars) and 2019 transfer data (light bars). (a) Weighted F1 score. (b) Macro F1 score, which weights all species equally regardless of prevalence. (c) Proportion L1 error, measuring species composition estimation accuracy across all parcels. Percentages above bars indicate relative performance degradation from 2018 to 2019. Error bars show ±1 standard deviation across 5 cross-validation folds.

**Table 6:** Temporal transfer performance. MLP classifiers trained on 2018 embeddings evaluated on held-out 2018 parcels (within-year) and 2019 parcels (cross-year transfer). Values are mean ± s.d. across 5-fold cross-validation.

| Input features | Weighted F1 |  | Macro F1 |  | PropL1 |  |
| --- | --- | --- | --- | --- | --- | --- |
|  | 2018 | 2019 | 2018 | 2019 | 2018 | 2019 |
| Tessera | $0.806 \pm 0.005$ | $0.731 \pm 0.004$ | $0.470 \pm 0.019$ | $0.408 \pm 0.008$ | $0.345 \pm 0.003$ | $0.393 \pm 0.005$ |
| AlphaEarth | $0.815 \pm 0.008$ | $0.692 \pm 0.011$ | $0.475 \pm 0.017$ | $0.354 \pm 0.009$ | $0.377 \pm 0.002$ | $0.452 \pm 0.011$ |

Within-year performance was strong for both foundation-model representations, confirming that classifiers trained on 2018 embeddings generalise well spatially under consistent observation conditions. When transferred to 2019 embeddings without retraining, performance declined for both models, indicating sensitivity to interannual variation in phenology, acquisition conditions, and/or management effects. However, the magnitude of degradation differed substantially between representations.

Tessera embeddings demonstrated greater temporal stability across all metrics. Weighted F1 decreased by 9.3% relative to within-year performance, compared to a 15.1% decline for AlphaEarth. Differences were more pronounced for macro F1, which weights all species equally: Tessera exhibited a 13.1% reduction, whereas AlphaEarth declined by 25.5%. Parcel-level composition error (PropL1) increased by 14.1% for Tessera and 19.9% for AlphaEarth under cross-year transfer.

The disproportionately large drop in macro F1 relative to weighted F1 for both models indicates that minority species are disproportionately affected by temporal domain shift, likely because their narrower spectral-temporal signatures and smaller training samples leave less margin for interannual variation. Nonetheless, Tessera consistently retained a greater proportion of its within-year performance, suggesting that its purely remote-sensing–driven self-supervised training may capture spectral–temporal features that are comparatively more invariant across years.

Overall, these results indicate that while both foundation-model embeddings support strong within-year classification, temporal robustness remains limited, and interannual domain shift represents a significant constraint for operational species mapping. Among the evaluated representations, Tessera exhibits comparatively greater stability under cross-year transfer, though neither model achieves fully temporally invariant performance.

## 4. Discussion

This study shows that geospatial foundation-model embeddings enable accurate and label-efficient tree species mapping in a highly heterogeneous alpine landscape, consistently outperforming conventional Sentinel-1+2 composites. Below, we discuss each experimental finding in turn: the nature and magnitude of the performance advantage (Section 4.1), the limited role of classifier complexity (4.2) and environmental covariates (4.3), the implications of soft-label supervision (4.4), temporal transferability as a remaining challenge (4.5), the broader operational context (4.6), limitations (4.7) and future directions (4.8).

### 4.1. Why foundation-model embeddings outperform multispectral baselines

The superior performance of geospatial foundation-model embeddings relative to conventional, seasonal multispectral Sentinel-1+2 composites reflects a difference in how ecological information is represented. Traditional species-mapping approaches rely on hand-crafted spectral features derived from single dates or seasonal summaries, which are only weak proxies for the structural, phenological, biochemical and functional traits that differentiate tree species in heterogeneous forests. In contrast, foundation models are trained to integrate spectral information across time and sensor modalities, producing representations that integrate over cloud gaps, atmospheric variability, and irregular acquisition patterns while preserving ecologically meaningful signals (Xiao et al., 2025). Although recent standardised benchmarks show that GFMs do not consistently outperform supervised baselines across all Earth observation tasks (Marsocci et al., 2025), our results demonstrate that species-level forest classification in complex terrain is a domain where their advantages are pronounced.

The label efficiency observed here – with foundation-model embeddings reaching near-asymptotic performance using ≤5% of available training parcels – suggests that discriminative structure closely aligned with species-relevant traits is already present in the representation rather than being learned during downstream training. Notably, neither weighted nor macro F1 fully plateaus across the range of training fractions: weighted F1 continues to rise with a shallow gradient, while macro F1 climbs more steadily. This pattern confirms that discrimination of rare species remains the limiting constraint on overall performance, since additional training examples continue to improve rare-species classification long after common species are well resolved. This interpretation is reinforced by the UMAP projections of the embedding space, which reveal coherent organisation along environmental gradients, functional types, and taxonomic groupings without any task-specific supervision, consistent with representations that encode ecological structure rather than merely grouping pixels by spectral similarity.

These results compare favourably with the broader tree species classification literature. Reviews of satellite-based species mapping have noted that most studies use fewer than ten classes and often achieve overall accuracies in the range of 70–85% with conventional features (Fassnacht et al., 2016; Zhong et al., 2024). Previous work in the same study region using fused multispectral and hyperspectral airborne data achieved strong discrimination among dominant conifers but required data sources with limited spatial and temporal coverage (Dalponte et al., 2012), while national-scale Sentinel time-series approaches in comparable European forests have reported lower accuracies at coarser taxonomic resolution (Blickensdörfer et al., 2024). Here, we classify 18 species and species groups – a finer scheme – and achieve a weighted F1 of 0.83 using only 10 m resolution Sentinel-derived embeddings, without airborne or hyperspectral data. This level of performance in a topographically complex montane landscape with strong illumination and BRDF effects suggests that foundation-model representations overcome several of the key limitations historically associated with satellite-based species discrimination, including the need for carefully engineered multi-temporal feature sets and sensor-specific tuning (Immitzer et al., 2019). By contrast, multispectral composites collapse temporal variation into summary statistics, obscuring phenological dynamics critical for species discrimination (Xu et al., 2021). Richer multi-temporal feature sets can partially compensate, but at considerable processing overhead and sensitivity to cloud cover (Wang et al., 2022) – challenges that foundation models sidestep by learning directly from complete, multi-year image collections.

An interesting nuance is the pattern of relative advantage between the two foundation models investigated in our study. Recent comparative evaluations of GFM embeddings for tree species tasks have generally treated them as interchangeable feature extractors (Ishikawa et al., 2025), but our results reveal a more differentiated picture. Tessera outperforms AlphaEarth on macro F1 (0.551 vs. 0.537) and parcel-level proportion error (PropL1 = 0.349 vs. 0.384), while AlphaEarth achieves marginally higher weighted F1 (0.833 vs. 0.824; weighted F1 was not prioritised in classification tuning). This divergence is not straightforwardly explained by embedding dimensionality, since Tessera’s advantage is concentrated in minority-species discrimination and compositional fidelity rather than in overall pixel accuracy. Instead, the two models’ architectural differences offer a more compelling explanation. AlphaEarth operates at the patch level and is trained using auxiliary targets spanning multiple Earth-system modalities, including lidar structure, climate, and topography. This design intends to prioritise robustness and generality across tasks and spatial contexts, but may over-smooth fine-grained spectral–phenological variation within parcels, explaining its weaker recovery of within-parcel species proportions. Tessera, by contrast, operates at the pixel level and is trained purely from remote-sensing time series using a self-supervised Barlow Twins objective (Lisaius et al., 2024). Its representations may therefore preserve canopy-trait variation that is particularly informative for differentiating co-occurring species and recovering realistic species mixtures at the parcel scale. This trade-off highlights that no single foundation-model representation is currently universally optimal: embeddings tuned for broad environmental generalisation may sacrifice local discriminative power, while models focused on sensor-derived signals may excel in region-specific ecological tasks and, as shown in our temporal transfer results, can retain greater stability across years.

The ecological coherence of the error structure provides further evidence that foundation-model representations encode biologically meaningful variation. Misclassifications are not randomly distributed but are concentrated along taxonomic and functional axes: silver fir (Abies alba) is predominantly confused with Norway spruce (Picea abies), consistent with the two genera sharing similar crown architecture, shade tolerance, and co-occurrence in the same montane conifer belt – a confusion pair that has been identified as a persistent challenge in the same region using both airborne and spaceborne sensors (Dalponte et al., 2012, 2025). Pine species (*P. sylvestris, P. nigra, P. cembra*) form a distinct confusion cluster, reflecting their shared needle morphology and overlapping spectral signatures despite occupying different elevational niches. Among broadleaf taxa, the merged community groups (e.g. oak–hornbeam, hop-hornbeam–manna ash, maple–ash–lime) are frequently confused with one another but rarely with coniferous classes, indicating that the embeddings reliably separate functional types even where species-level resolution remains limited. These patterns suggest that the representations capture a hierarchical ecological signal – functional type, genus, and species – and that residual errors largely reflect genuine biochemical–structural similarity among co-occurring taxa rather than arbitrary noise. For forest management, the practical consequence is that the most frequent misclassifications occur between species that often share similar silvicultural treatment, limiting the operational impact of these errors.

Taken together, these findings indicate that the performance advantage of foundation models stems primarily from a more expressive and ecologically aligned representation of the landscape itself, rather than from downstream modelling choices.

### 4.2. Classifier capacity: necessary but quickly saturating

A natural question arising from the strong performance of foundation-model embeddings is whether additional gains can be achieved through increased classifier capacity. Our results show that classifier choice matters – a linear classifier applied to foundation-model embeddings underperforms an MLP applied to conventional S1+2 seasonal composites – but that returns diminish rapidly once a moderately nonlinear model is used.

The step from linear to nonlinear classifiers produced the largest single-factor improvement observed in the study: macro F1 increased by +0.17 on Tessera embeddings, exceeding the gain from switching input representations with a fixed MLP (+0.05 from S1+2 seasonal to Tessera). This indicates that the discriminative information in foundation-model embeddings is not linearly separable, and that a nonlinear classifier is required to exploit it. However, once this threshold is crossed, further increases in complexity yield bounded gains. Shallow MLPs provided consistent improvements over Random Forest baselines – the dominant classifier in operational remote-sensing classification (Belgiu and Drăguţ, 2016) – primarily reflected in macro-averaged performance (+0.08 on Tessera), while deeper or more flexible architectures failed to yield further improvements. The concentration of performance gains in macro F1 indicates that nonlinear classifiers mainly benefit minority species, likely by smoothing decision boundaries in regions of the embedding space where rare classes form small but coherent clusters. The relative stability of weighted F1 across nonlinear classifiers implies that, for dominant species, the representation is already sufficiently structured for tree-based or instance-based methods to achieve strong performance.

The nature of the discriminative signal offers a mechanistic explanation for this pattern. Species identity is encoded in a pixel’s spectral–temporal trajectory – phenology, leaf biochemistry – which is captured by per-pixel embeddings. Tessera’s training explicitly decorrelates spatial neighbourhoods through global shuffling, so the resulting embeddings encode spectral–temporal properties of individual pixels rather than spatial structure. For species mapping this is precisely the relevant signal, and a per-pixel MLP is sufficient to exploit it. By contrast, structural attributes such as canopy height or biomass may benefit from spatial context – shadow geometry, crown texture, gap structure – explaining why the same embeddings benefit from spatial decoders such as UNets when applied to regression tasks (Feng et al., 2025).

From a practical perspective, these results indicate that at least a shallow nonlinear classifier is needed to exploit foundation-model embeddings for species mapping, but that pursuing increasingly complex architectures beyond this point offers little additional benefit.

### 4.3. The limited role of ancillary environmental covariates

Environmental variables such as elevation, slope, and topographic position are well known to structure forest composition in mountain landscapes (Weiss and Walsh, 2009). These covariates are therefore commonly incorporated into species distribution and classification models to compensate for topography-induced spectral distortions (Dong et al., 2020) and to exploit the strong species–site relationships in montane forests (Pu, 2021). It is therefore reasonable to ask whether explicitly providing such covariates can improve species discrimination beyond what is achievable with foundation-model embeddings alone. In this study, however, we find no evidence that ancillary environmental data provide meaningful additional predictive power.

Augmenting foundation-model embeddings with terrain-derived variables yielded no improvement in classification performance for either representation. For AlphaEarth, this outcome is perhaps unsurprising: its training explicitly incorporates DEM-derived topographic targets among its auxiliary objectives, so elevation and slope information is likely already directly encoded in the embedding. For Tessera, which is trained purely from optical and SAR time series without any explicit environmental inputs, the null result is more noteworthy. One interpretation is that key abiotic gradients are already indirectly captured through their ecological effects on canopy phenology, structure, and biochemical composition – a hypothesis supported by the UMAP visualisation showing clear elevation-related structure in Tessera embedding space – but alternative explanations should be considered. Simple feature concatenation may be a suboptimal integration strategy, the 30 m DEM resampled to 10 m may be too coarse to add meaningful local information, and the five terrain-derived variables tested (elevation, slope, aspect, TPI, curvature) may not capture the most relevant environmental gradients for species discrimination. The null result therefore demonstrates that terrain covariates are redundant under the specific experimental design employed here, but does not establish that all environmental information is already encoded in the embeddings.

For species mapping, the practical implication is that incorporating terrain-derived covariates via simple concatenation adds complexity, data dependencies, and preprocessing overhead without delivering commensurate gains in accuracy under the conditions tested. Whether alternative integration strategies (e.g. attention-based fusion, multi-scale environmental features, or higher-resolution terrain data) could yield improvements remains an open question. While explicit environmental data may still play an important role in other ecological modelling tasks – particularly those involving extrapolation beyond observed conditions – the present results suggest that in this study region they are not a prerequisite for effective species discrimination when using foundation-model representations.

### 4.4. Label quality, soft supervision, and the limits of hard classification

A common assumption in species-mapping workflows is that label noise represents a primary constraint on model performance, particularly when training data are derived from coarse or mixed inventory units (Bolyn et al., 2022). Our results paint a more nuanced picture: classification from foundation-model embeddings is surprisingly robust to moderate label impurity, hard-label refinement strategies did not help, but reframing supervision as soft labels unlocks gains that neither approach achieves alone.

The robustness to label impurity is notable, particularly given the wider literature showing that training-set size and composition can substantially affect supervised classification accuracy (Ramezan et al., 2021). Under hard-label training, performance remains nearly flat from 30% to 80% dominant-species purity, with aggressive filtering to near-monocultures (≥90%) degrading performance as data volume collapses. We hypothesise that foundation-model embeddings provide sufficient spectral-temporal structure to tolerate moderate label uncertainty: at lower purity thresholds, increased noise is offset by greater sample availability, while at very high thresholds, data scarcity degrades performance, particularly for minority species. The failure of label distillation (Section S.4) – iteratively reassigning hard labels within parcels using embedding-space prototypes – to improve pixel-level accuracy reinforces this interpretation: at 10 m resolution, where individual pixels integrate reflectance from multiple tree crowns, re-sorting discrete labels within a parcel cannot recover information that hard labels are structurally unable to encode.

The success of soft-label training, by contrast, reveals a deeper issue. The distinction between hard-label distillation and soft-label training is not merely technical but epistemic. The species composition of a parcel is known from forest inventory data, but the spatial arrangement of species within that parcel is not; assigning each pixel a single species therefore fabricates pixel-level certainty from parcel-level knowledge. Simultaneously, a 10 m Sentinel-2 pixel integrates reflectance from multiple tree crowns and is itself a spectral mixture. Hard labels thus impose a single-species target on a mixed-species input, using information that was never spatially resolved. Soft labels resolve both mismatches, encoding exactly the compositional knowledge available from the inventory while presenting the model with targets that are consistent with the mixed nature of the spectral input.

The consequences are visible across the purity curve (Section 3.4; Figure 10). Where hard-label training produces an optimum at ∼80% purity, with performance falling on either side, the soft-label curve continues to improve from 30% to 50% purity. Mixed-species stands, which are noise sources under hard labelling, become correctly labelled training data under soft supervision, eliminating the trade-off between label quality and data volume. This pattern is consistent across both Tessera and AlphaEarth embeddings, with the latter showing a larger macro F1 advantage (+0.024 at peak), suggesting that soft labels can partially compensate for weaker embedding-level species discrimination. The improvement in Proportion L1 error (0.006–0.008 lower across the stable purity range for both embeddings) confirms that soft labels also improve recovery of landscape-scale species proportions. The macro F1 gains indicate that the primary beneficiaries are less common species that often occur as secondary components in mixed stands rather than forming monospecific parcels; under hard labelling their training signal is systematically suppressed whenever they are not the parcel majority, and soft labels recover it. Given that parcel-level species composition is already routinely recorded in national forest inventories (Brus et al., 2012), and that the majority of real forest parcels contain some degree of species mixing, these results suggest that the widespread practice of converting fractional composition data into hard per-pixel labels discards substantial usable information – and that the limiting factor is not label noise per se, but the assumption that supervision must be discrete. Our approach is complementary to recent work on sub-pixel species-fraction mapping using synthetically mixed training spectra (Klehr et al., 2025), which tackles the same fundamental problem – mixed pixels and discrete labels – from the feature-engineering side. Together, these lines of evidence suggest that moving from hard classification to fractional or soft-label formulations represents a broad methodological shift rather than an incremental refinement.

### 4.5. Temporal transferability as the remaining frontier

Despite the strong within-year performance achieved by geospatial foundation-model embeddings, our results show that temporal generalisation under single-year training merits further attention for species-level mapping, though straightforward strategies such as multi-year training offer a clear path forward. Domain shift between training and deployment periods is a well-known limitation in remote-sensing classification (Tuia et al., 2016), and a key question for foundation-model representations is whether their large-scale pre-training confers temporal robustness that task-specific models lack. When models trained on 2018 embeddings were applied to 2019 data, classification accuracy declined substantially, with the most pronounced losses observed in macro-averaged metrics. This pattern indicates that rare and less dominant species are particularly sensitive to year-to-year domain shifts, likely reflecting incomplete coverage of species-specific spectral-temporal variability within single-year training data.

Several factors are likely to contribute to this temporal sensitivity. Interannual variation in phenology, driven by climatic anomalies or shifts in seasonal timing, can alter the spectral and phenological signals encoded in the embeddings (Andreatta et al., 2025). Differences in snow persistence, cloud availability, and acquisition geometry may further exacerbate domain shift between years. Critically, the specific year-pair evaluated here (2018–2019) coincides with a major landscape disturbance: Storm Vaia struck the eastern Alps in late October 2018, part of a broader pattern of intensifying forest disturbance across Europe (Seidl et al., 2017; Senf and Seidl, 2021), causing widespread windthrow across Trentino and triggering extensive salvage logging in subsequent months. The resulting changes in canopy structure, gap formation, and management activity represent genuine compositional and structural change between the training and transfer years, making it impossible to fully separate representational drift from real landscape dynamics. More broadly, forest inventories are often temporally asynchronous, and genuine changes in species composition due to disturbance, harvesting, or management can introduce discrepancies between training labels and validation data that are difficult to disentangle from representational drift. Because these results are based on a single year-pair in one study region, the magnitude of temporal degradation should not be generalised without evaluation across additional years and landscapes.

From an operational perspective, this sensitivity has important implications for applications such as change detection and the production of consistent, multi-year habitat maps. If temporal variability in embeddings reflects both ecological change and observational effects, naïvely applying static classifiers across years risks conflating phenological variation with true compositional change. Robust long-term monitoring will therefore require either temporally invariant representations or multi-year training strategies to ensure consistency across years.

At the same time, the observed interannual variation in embedding space may represent an opportunity rather than solely a limitation. Because foundation-model embeddings respond to phenological, bio-chemical and structural changes, their temporal trajectories could potentially be exploited to study shifts in phenology, stress responses, or disturbance dynamics, even when species identity remains constant. Distinguishing stable species-specific traits from temporally varying ecological signals offers a promising direction for future work, bridging species mapping with functional and phenological monitoring. Leveraging the spatial variability of the derived embeddings provides an easy way to quantify trait and species variability, e.g. a simple biodiversity indicator.

Under cross-year transfer, Tessera retains a greater proportion of its within-year performance than AlphaEarth, suggesting that its purely remote-sensing–driven self-supervised training captures spectral–temporal features that are comparatively more invariant across years. Nonetheless, the magnitude of performance degradation observed for both models indicates that current foundation-model embeddings produce representations that respond to interannual variation in phenology and acquisition conditions, such that direct year-to-year transfer without adaptation incurs some performance loss – consistent with early evidence from embedding-based crop classification, where temporal alignment of training and inference data remains important (Lisaius et al., 2026). Independent evaluation of AlphaEarth for agricultural tasks has similarly identified limited time sensitivity as a key constraint of the current embedding design (Ma et al., 2025), suggesting that interannual variability in frozen foundation-model embeddings is a systematic consideration for classification tasks rather than an artefact of our specific study region.

Overall, these findings identify temporal generalisation as a critical frontier for geospatial foundation models. Addressing it – through multi-year stratified training, explicit temporal domain adaptation, or phenology-aware representation learning – will be essential both for reliable change detection and for leveraging temporal variation as an ecological signal rather than a source of noise.

### 4.6. Implications for operational forest monitoring

These findings have direct implications for how national forest inventory programmes and conservation agencies approach species-level mapping. National forest inventories increasingly seek to integrate field plots with remotely sensed wall-to-wall products (McRoberts and Tomppo, 2007; White et al., 2016; Lister et al., 2020), but this integration has traditionally required extensive task-specific feature engineering and large labelled training sets (McRoberts et al., 2010). The demonstration that foundation-model embeddings achieve strong classification performance with minimal downstream complexity and as few as 5% of available training parcels suggests that regional species maps could be produced with substantially reduced labelling effort compared to conventional approaches – lowering the barrier for inventory programmes with limited resources or spatial coverage. For agencies that already collect parcel-level species proportions as part of standard forest assessment, the soft-label finding is particularly actionable: existing compositional data that are routinely discarded when converted to hard labels can instead be used directly as training targets, yielding improved discrimination of minority species without any additional field effort.

The practical value of these advances should be viewed in the context of emerging policy needs. The EU Biodiversity Strategy for 2030 and associated Forest Strategy call for improved monitoring of forest condition, composition, and change across European landscapes (European Commission, 2021). Wall-to-wall species maps at 10 m resolution, such as the one produced here for Trentino (Figure 8), represent a step beyond the genus-level products currently available at European scale (De Keersmaecker et al., 2024) – provided that the temporal transfer challenge can be addressed. The combination of foundation-model representations with existing inventory infrastructure offers a scalable pathway for producing and updating species composition maps, but realising this potential at continental scales will require attention to cross-regional transferability, harmonisation of inventory protocols, and the development of temporally robust training strategies.

### 4.7. Limitations

Several limitations should be noted when interpreting these results. First, all experiments are conducted in a single study region (Trentino), which, while ecologically complex and demanding, represents only one biogeographic setting; the degree to which findings generalise to other forest types, climatic zones, or inventory systems remains to be tested. Second, temporal transfer is evaluated over a single year-pair (2018–2019) that coincides with Storm Vaia (Section 4.5), making it difficult to fully disentangle representational drift from genuine landscape change. Third, we adopt a pixel-level hard classification framework rather than sub-pixel fractional prediction. At Sentinel-2’s 10 m resolution, individual pixels typically capture one or a few tree crowns, limiting the degree of sub-pixel species mixing; moreover, pixel-level classification is broadly compatible with existing operational mapping workflows and inventory systems. Our soft-label experiments further demonstrate that compositional reference data can be exploited effectively within this framework, though a fully fractional formulation could offer additional benefits in genuinely mixed stands. Fourth, foundation-model embeddings are evaluated in a frozen, off-the-shelf configuration without fine-tuning; task-specific adaptation has shown promise for vegetation mapping (Ishikawa et al., 2025) and could potentially improve performance, particularly for rare species, but would complicate the assessment of representation quality per se. Similarly, hierarchical classification schemes that exploit the taxonomic structure of species labels (Jin and Wu, 2025) or hyperspectral foundation models with richer spectral resolution (Braham et al., 2025) remain unexplored directions that could further improve discrimination of spectrally similar taxa. Finally, the reference data are parcel-level inventory proportions rather than individual tree-level labels; while this is standard for forest inventory studies, it means that pixel-level predictions are validated against spatially aggregated proportions rather than at the scale of individual stems.

### 4.8. Future directions

The strong label efficiency demonstrated here – near-asymptotic performance from ≤5% of parcels – raises the question of whether even sparser supervision could suffice if training samples are selected more strategically. In preliminary experiments comparing alternative sampling strategies (Section S.7.4; Table S.13), coreset selection guided by K-means clustering in the embedding space consistently outperformed random stratified sampling across both foundation models, improving weighted F1 by up to 1.6 percentage points at 5% training fraction and by 3.3 points at 0.1% (∼45 parcels). These results point toward few-shot species mapping in data-sparse regions, where a small number of strategically selected reference parcels – guided by unsupervised structure in the embedding space – may be sufficient for operational classification without region-specific campaigns. More broadly, extending the framework to multi-year training with explicit temporal domain adaptation, sub-pixel fractional prediction building on the soft-label formulation explored here, and task-specific fine-tuning of embeddings would address several of the limitations noted above and move the approach closer to wall-to-wall operational deployment.

### 4.9. Conclusions

Geospatial foundation-model embeddings enable accurate and label-efficient tree species mapping in complex alpine forests, consistently outperforming conventional Sentinel-1+2 composites when used with a nonlinear classifier. Foundation-model embeddings encode species-discriminative information more effectively than conventional composites, with the advantage most pronounced for minority species; all representations reach near-asymptotic performance with limited training data, but embeddings do so at a consistently higher ceiling. Realising this advantage requires a nonlinear classifier – the step from linear to nonlinear models produces the single largest improvement – but once moderate nonlinearity is provided, further increases in model capacity yield diminishing returns. Classification is robust to moderate label impurity, and training with soft species-composition labels rather than hard per-pixel assignments substantially improves minority-species discrimination, suggesting that converting fractional inventory data into discrete labels discards usable information. Supplementary environmental covariates provide no additional predictive power when concatenated with foundation-model embeddings, though alternative integration strategies remain untested. Temporal transfer across years reveals marked performance degradation, particularly for rare species, with embedding choice influencing temporal stability.

Overall, these results suggest that geospatial foundation models shift a primary bottleneck in species mapping from feature engineering toward the availability, quality, and temporal alignment of ecological reference data. Realising their full potential for operational biodiversity monitoring will require temporally consistent training strategies and reference datasets that embrace the compositional complexity of real forest stands.

## Data and code availability statement

The forest inventory data used in this study were provided by the Autonomous Province of Trento and are available upon reasonable request to the corresponding author. Processed inventory parcels, foundation-model embeddings, other intermediate data and landscape predictions are available on Zenodo: https://zenodo.org/records/18924120. Code is available at https://github.com/PatBall1/trentino-trees.

## Declaration of generative AI and AI-assisted technologies in the writing process

During the preparation of this work, the authors used AI-assisted tools to assist with editing and proofreading text drafted by the authors. The authors reviewed and edited all AI-assisted output and take full responsibility for the content of this publication.

## Acknowledgements

JGCB was supported by the Medical Research Council (MR/Z505456/1). SJ was funded by a charitable donation from John Bernstein. ZF was supported by the Robert Sansom Studentship.

## CRediT authorship contribution statement

JGCB: Conceptualization, Methodology, Software, Formal analysis, Visualization, Writing – original draft, Writing – review & editing, Project administration. JAW: Data curation, Investigation, Writing – review & editing. ZF: Software, Writing – review & editing. JK: Writing – review & editing. SJ: Software, Writing – review & editing. AM: Supervision, Funding acquisition, Resources, Writing – review & editing. CA: Conceptualization, Methodology, Writing – review & editing, Supervision. MD: Data curation, Resources, Writing – review & editing, Supervision. DAC: Conceptualization, Supervision, Funding acquisition, Writing – review & editing.

## List of Abbreviations

AEF: AlphaEarth Foundations
BRDF: Bidirectional Reflectance Distribution Function
CLC: CORINE Land Cover
CLMS: Copernicus Land Monitoring Service
DEM: Digital Elevation Model
F1 F1: score (harmonic mean of precision and recall)
GFM: Geospatial Foundation Model
GRD: Ground Range Detected (Sentinel-1 product)
IW: Interferometric Wide swath (Sentinel-1 mode)
KL: Kullback–Leibler (divergence)
KNN: K-Nearest Neighbours
LR: Logistic Regression
MF1: Macro-averaged F1 score
MLP: Multi-Layer Perceptron
NBR: Normalised Burn Ratio
NDVI: Normalised Difference Vegetation Index
NDWI: Normalised Difference Water Index
PropL1: Proportion L1 error (parcel-level metric)
RF: Random Forest
SAR: Synthetic Aperture Radar
SCL: Scene Classification Layer (Sen2Cor)
SRTM: Shuttle Radar Topography Mission
TPI: Topographic Position Index
UMAP: Uniform Manifold Approximation and Projection
UTM: Universal Transverse Mercator
VV/VH: Vertical-Vertical / Vertical-Horizontal (SAR polarisations)
WF1: Weighted F1 score

## Footnotes

1 https://github.com/ucam-eo/geotessera

## References

Agnoletti, M., Biasio, R., 2013. Trentino Alto Adige, in: Agnoletti, M. (Ed.), Italian Historical Rural Landscapes: Cultural Values for the Environment and Rural Development. Springer Netherlands, Dordrecht, pp. 247–262. 10.1007/978-94-007-5354-9_10

Akiba, T., Sano, S., Yanase, T., Ohta, T., Koyama, M., 2019. Optuna: A Next-generation Hyperparameter Optimization Framework. 10.48550/arXiv.1907.10902

Andreatta, D., Buchmann, N., Jucker, T., Belelli Marchesini, L., Dalponte, M., Scotton, M., Vescovo, L., Gianelle, D., 2025. Diverging trends in plant phenology and productivity across European mountains in a warming world. Agric. For. Meteorol. 375, 110874. 10.1016/j.agrformet.2025.110874

Bai, J., Ren, C., Shi, X., Xiang, H., Zhang, W., Jiang, H., Ren, Y., Xi, Y., Wang, Z., Mao, D., 2024. Tree species diversity impacts on ecosystem services of temperate forests. Ecol. Indic. 167, 112639. 10.1016/j.ecolind.2024.112639

Ball, J.G.C., Jaffer, S., Laybros, A., Prieur, C., Jackson, T., Madhavapeddy, A., Barbier, N., Vincent, G., Coomes, D.A., 2024. Harnessing temporal and spectral dimensionality to map and identify species of individual trees in diverse tropical forests. bioRxiv 2024.06.24.600405.

Belgiu, M., Drăguţ, L., 2016. Random forest in remote sensing: A review of applications and future directions. ISPRS J. Photogramm. Remote Sens. 114, 24–31. 10.1016/j.isprsjprs.2016.01.011

Bergseng, E., Ørka, H.O., Næsset, E., Gobakken, T., 2015. Assessing forest inventory information obtained from different inventory approaches and remote sensing data sources. Ann. For. Sci. 72, 33–45. 10.1007/s13595-014-0389-x

Blickensdörfer, L., Oehmichen, K., Pflugmacher, D., Kleinschmit, B., Hostert, P., 2024. National tree species mapping using Sentinel-1/2 time series and German National Forest Inventory data. Remote Sens. Environ. 304, 114069. 10.1016/j.rse.2024.114069

Bolyn, C., Lejeune, P., Michez, A., Latte, N., 2022. Mapping tree species proportions from satellite imagery using spectral–spatial deep learning. Remote Sens. Environ. 280, 113205. 10.1016/j.rse.2022.113205

Braham, N.A.A., Albrecht, C.M., Mairal, J., Chanussot, J., Wang, Y., Zhu, X.X., 2025. SpectralEarth: Training Hyperspectral Foundation Models at Scale. IEEE J. Sel. Top. Appl. Earth Obs. Remote Sens. 18, 16780–16797. 10.1109/JSTARS.2025.3581451

Brandt, M., Chave, J., Li, S., Fensholt, R., Ciais, P., Wigneron, J.-P., Gieseke, F., Saatchi, S., Tucker, C.J., Igel, C., 2025. High-resolution sensors and deep learning models for tree resource monitoring. Nat. Rev. Electr. Eng. 2, 13–26. 10.1038/s44287-024-00116-8

Bray, J.R., Curtis, J.T., 1957. An Ordination of the Upland Forest Communities of Southern Wisconsin. Ecol. Monogr. 27, 326–349. 10.2307/1942268

Breiman, L., 2001. Random Forests. Mach. Learn. 45, 5–32. 10.1023/A:1010933404324

Brockerhoff, E.G., Barbaro, L., Castagneyrol, B., Forrester, D.I., Gardiner, B., González-Olabarria, J.R., Lyver, P.O., Meurisse, N., Oxbrough, A., Taki, H., Thompson, I.D., van der Plas, F., Jactel, H., 2017. Forest biodiversity, ecosystem functioning and the provision of ecosystem services. Biodivers Conserv 26, 3005–3035.

Brown, C.F., Kazmierski, M.R., Pasquarella, V.J., Rucklidge, W.J., Samsikova, M., Zhang, C., Shelhamer, E., Lahera, E., Wiles, O., Ilyushchenko, S., Gorelick, N., Zhang, L.L., Alj, S., Schechter, E., Askay, S., Guinan, O., Moore, R., Boukouvalas, A., Kohli, P., 2025. AlphaEarth Foundations: An embedding field model for accurate and efficient global mapping from sparse label data. 10.48550/arXiv.2507.22291

Brus, D.J., Hengeveld, G.M., Walvoort, D.J.J., Goedhart, P.W., Heidema, A.H., Nabuurs, G.J., Gunia, K., 2012. Statistical mapping of tree species over Europe. Eur. J. For. Res. 131, 145–157. 10.1007/s10342-011-0513-5

Cover, T., Hart, P., 1967. Nearest neighbor pattern classification. IEEE Trans. Inf. Theory 13, 21–27. 10.1109/TIT.1967.1053964

Dalponte, M., Andreatta, D., Coomes, D.A., Belelli Marchesini, L., Marinelli, D., Vescovo, L., Gianelle, D., 2025. Canopy spectral responses of temperate forests to late spring frost and hot drought events assessed with Sentinel-2 NDVI time series. Remote Sens. Appl. Soc. Environ. 40, 101737. 10.1016/j.rsase.2025.101737

Dalponte, M., Bruzzone, L., Gianelle, D., 2012. Tree species classification in the Southern Alps based on the fusion of very high geometrical resolution multispectral/hyperspectral images and LiDAR data. Remote Sens. Environ. 123, 258–270. 10.1016/j.rse.2012.03.013

De Keersmaecker, W., Zanaga, D., Senf, C., Viana-Soto, A., Klapper, J., Blickensdörfer, L., Govaere, L., Lerink, B., Leyman, A., Schelhaas, M.-J., Teeuwen, S., Verkerk, P.J., Van De Kerchove, R., 2024. ForestPaths: European tree genus map. 10.5281/zenodo.13341104

Dong, C., Zhao, G., Meng, Y., Li, B., Peng, B., 2020. The Effect of Topographic Correction on Forest Tree Species Classification Accuracy. Remote Sens. 12, 787. 10.3390/rs12050787

European Commission, 2021. New EU Forest Strategy for 2030.

Farr, T.G., Rosen, P.A., Caro, E., Crippen, R., Duren, R., Hensley, S., Kobrick, M., Paller, M., Rodriguez, E., Roth, L., Seal, D., Shaffer, S., Shimada, J., Umland, J., Werner, M., Oskin, M., Burbank, D., Alsdorf, D., 2007. The Shuttle Radar Topography Mission. Rev. Geophys. 45. 10.1029/2005RG000183

Fassnacht, F.E., Latifi, H., Stereńczak, K., Modzelewska, A., Lefsky, M., Waser, L.T., Straub, C., Ghosh, A., 2016. Review of studies on tree species classification from remotely sensed data. Remote Sens Env. 186, 64–87.

Feng, Z., Atzberger, C., Jaffer, S., Knezevic, J., Sormunen, S., Young, R., Lisaius, M.C., Immitzer, M., Ball, J., Coomes, D.A., Madhavapeddy, A., Blake, A., Keshav, S., 2025. TESSERA: Temporal Embeddings of Surface Spectra for Earth Representation and Analysis. 10.48550/arXiv.2506.20380

Frei, E.R., Barbeito, I., Erdle, L.M., Leibold, E., Bebi, P., 2023. Evidence for 40 Years of Treeline Shift in a Central Alpine Valley. Forests 14, 412. 10.3390/f14020412

Gao, X., Chi, H., Huang, J., Han, Y., Li, Y., Ling, F., 2024. Comparison of Cloud-Mask Algorithms and Machine-Learning Methods Using Sentinel-2 Imagery for Mapping Paddy Rice in Jianghan Plain. Remote Sens. 16. 10.3390/rs16071305

Gasparini, P., Di Cosmo, L., Floris, A., De Laurentis, D. (Eds.), 2022. Italian National Forest Inventory—Methods and Results of the Third Survey: Inventario Nazionale delle Foreste e dei Serbatoi Forestali di Carbonio—Metodi e Risultati della Terza Indagine, Springer Tracts in Civil Engineering. Springer International Publishing, Cham. 10.1007/978-3-030-98678-0

Germain, S.J., Lutz, J.A., 2024. Stand diversity increases pine resistance and resilience to compound disturbance. Fire Ecol. 20, 53. 10.1186/s42408-024-00283-x

Gibbs, A.L., Su, F.E., 2002. On Choosing and Bounding Probability Metrics. Int. Stat. Rev. 70, 419–435. 10.1111/j.1751-5823.2002.tb00178.x

Hastie, T., Tibshirani, R., Friedman, J., 2009. Linear Methods for Classification, in: Hastie, T., Tibshirani, R., Friedman, J. (Eds.), The Elements of Statistical Learning: Data Mining, Inference, and Prediction. Springer, New York, NY, pp. 101–137. 10.1007/978-0-387-84858-7_4

Hu, Z., Xiao, F., Du, Y., Wang, Z., Luo, J., Feng, Q., Chen, M., 2025. Application of Landsat High Spatial Resolution Phenological Synthesized Data in Mountainous Land Cover Classification. Remote Sens. 17, 2603. 10.3390/rs17152603

Immitzer, M., Atzberger, C., 2023. Tree Species Diversity Mapping—Success Stories and Possible Ways Forward. Remote Sens. 15, 3074. 10.3390/rs15123074

Immitzer, M., Neuwirth, M., Böck, S., Brenner, H., Vuolo, F., Atzberger, C., 2019. Optimal Input Features for Tree Species Classification in Central Europe Based on Multi-Temporal Sentinel-2 Data. Remote Sens. 11, 2599. 10.3390/rs11222599

Ishikawa, T., Bonannella, C., Lerink, B.J.W., Rußwurm, M., 2025. Deep Pre-trained Time Series Features for Tree Species Classification in the Dutch Forest Inventory. 10.48550/arXiv.2508.18829

Jin, Y., Wu, C., 2025. Plant Taxonomy Meets Deep Learning: Hierarchical Classification for Fine-Grained Tree Species Identification with Hyperspectral Foundation Models. 10.2139/ssrn.5573998

Kellndorfer, J.M., Pierce, L.E., Dobson, M.C., Ulaby, F.T., 1998. Toward consistent regional-to-global-scale vegetation characterization using orbital SAR systems. IEEE Trans. Geosci. Remote Sens. 36, 1396–1411. 10.1109/36.718844

Klehr, D., Stoffels, J., Hill, A., Pham, V.-D., van der Linden, S., Frantz, D., 2025. Mapping tree species fractions in temperate mixed forests using Sentinel-2 time series and synthetically mixed training data. Remote Sens. Environ. 323, 114740. 10.1016/j.rse.2025.114740

Lisaius, M.C., Blake, A., Keshav, S., Atzberger, C., 2024. Using Barlow Twins to Create Representations From Cloud-Corrupted Remote Sensing Time Series. IEEE J. Sel. Top. Appl. Earth Obs. Remote Sens. 17, 13162–13168. 10.1109/JSTARS.2024.3426044

Lisaius, M.C., Keshav, S., Blake, A., Atzberger, C., 2026. Embedding -based Crop Type Classification in the Groundnut Basin of Senegal. 10.48550/arXiv.2601.16900

Lister, A.J., Andersen, H., Frescino, T., Gatziolis, D., Healey, S., Heath, L.S., Liknes, G.C., McRoberts, R., Moisen, G.G., Nelson, M., Riemann, R., Schleeweis, K., Schroeder, T.A., Westfall, J., Wilson, B.T., 2020. Use of Remote Sensing Data to Improve the Efficiency of National Forest Inventories: A Case Study from the United States National Forest Inventory. Forests 11, 1364. 10.3390/f11121364

Ma, Y., Shen, Y., Swatantran, A., Lobell, D.B., 2025. Harvesting AlphaEarth: Benchmarking the Geospatial Foundation Model for Agricultural Downstream Tasks. 10.48550/arXiv.2601.00857

Marsocci, V., Jia, Y., Bellier, G.L., Kerekes, D., Zeng, L., Hafner, S., Gerard, S., Brune, E., Yadav, R., Shibli, A., Fang, H., Ban, Y., Vergauwen, M., Audebert, N., Nascetti, A., 2025. PANGAEA: A Global and Inclusive Benchmark for Geospatial Foundation Models. 10.48550/arXiv.2412.04204

McRoberts, R.E., Tomppo, E.O., 2007. Remote sensing support for national forest inventories. Remote Sens. Environ., ForestSAT Special Issue 110, 412–419. 10.1016/j.rse.2006.09.034

McRoberts, R.E., Tomppo, E.O., Næsset, E., 2010. Advances and emerging issues in national forest inventories. Scand. J. For. Res. 25, 368–381. 10.1080/02827581.2010.496739

Pedrotti, F., 2018. Vegetation Series Along Climatic Gradients in the Central Southern Alps (Trentino-Alto Adige Region), in: Pedrotti, F. (Ed.), Climate Gradients and Biodiversity in Mountains of Italy. Springer International Publishing, Cham, pp. 51–81. 10.1007/978-3-319-67967-9_3

Pu, R., 2021. Mapping Tree Species Using Advanced Remote Sensing Technologies: A State-of-the-Art Review and Perspective. J. Remote Sens. 2021. 10.34133/2021/9812624

Ramezan, C.A., Warner, T.A., Maxwell, A.E., Price, B.S., 2021. Effects of Training Set Size on Supervised Machine-Learning Land-Cover Classification of Large-Area High-Resolution Remotely Sensed Data. Remote Sens. 13. 10.3390/rs13030368

Rautiainen, M., Mõttus, M., Heiskanen, J., Akujärvi, A., Majasalmi, T., Stenberg, P., 2011. Seasonal reflectance dynamics of common understory types in a northern European boreal forest. Remote Sens. Environ. 115, 3020–3028. 10.1016/j.rse.2011.06.005

Rumelhart, D.E., Hinton, G.E., Williams, R.J., 1986. Learning representations by back-propagating errors. Nature 323, 533–536. 10.1038/323533a0

Scherrer, D., Vitasse, Y., Guisan, A., Wohlgemuth, T., Lischke, H., 2020. Competition and demography rather than dispersal limitation slow down upward shifts of trees’ upper elevation limits in the Alps. J. Ecol. 108, 2416–2430. 10.1111/1365-2745.13451

Schwaiger, F., Poschenrieder, W., Biber, P., Pretzsch, H., 2018. Species Mixing Regulation with Respect to Forest Ecosystem Service Provision. Forests 9, 632. 10.3390/f9100632

Seidl, R., Thom, D., Kautz, M., Martin-Benito, D., Peltoniemi, M., Vacchiano, G., Wild, J., Ascoli, D., Petr, M., Honkaniemi, J., Lexer, M.J., Trotsiuk, V., Mairota, P., Svoboda, M., Fabrika, M., Nagel, T.A., Reyer, C.P.O., 2017. Forest disturbances under climate change. Nat. Clim. Change 7, 395–402. 10.1038/nclimate3303

Senf, C., Seidl, R., 2021. Persistent impacts of the 2018 drought on forest disturbance regimes in Europe. Biogeosciences 18, 5223–5230. 10.5194/bg-18-5223-2021

Silva Pedro, M., Rammer, W., Seidl, R., 2015. Tree species diversity mitigates disturbance impacts on the forest carbon cycle. Oecologia 177, 619–630. 10.1007/s00442-014-3150-0

Thompson, I., Mackey, B., McNulty, S., Mosseler, A., 2009. Forest Resilience, Biodiversity, and Climate Change. Secr. Conv. Biol. Divers. Montr. Tech. Ser. No 43 1–67 43, 1–67.

Thrippleton, T., Temperli, C., Krumm, F., Mey, R., Zell, J., Stroheker, S., Gossner, M.M., Bebi, P., Thürig, E., Schweier, J., 2023. Balancing disturbance risk and ecosystem service provisioning in Swiss mountain forests: an increasing challenge under climate change. Reg. Environ. Change 23, 29.

Tilman, D., Kinzig, A.P., Pacala, S., 2013. The Functional Consequences of Biodiversity: Empirical Progress and Theoretical Extensions. Princeton University Press. 10.1515/9781400847303

Tuia, D., Persello, C., Bruzzone, L., 2016. Domain Adaptation for the Classification of Remote Sensing Data: An Overview of Recent Advances. IEEE Geosci. Remote Sens. Mag. 4, 41–57. 10.1109/MGRS.2016.2548504

Wang, M., Zheng, Y., Huang, C., Meng, R., Pang, Y., Jia, W., Zhou, J., Huang, Z., Fang, L., Zhao, F., 2022. Assessing Landsat-8 and Sentinel-2 spectral-temporal features for mapping tree species of northern plantation forests in Heilongjiang Province, China. For. Ecosyst. 9, 100032. 10.1016/j.fecs.2022.100032

Wang, R., Springer, K.R., Gamon, J.A., 2023. Confounding effects of snow cover on remotely sensed vegetation indices of evergreen and deciduous trees: An experimental study. Glob. Change Biol. 29, 6120–6138. 10.1111/gcb.16916

Weiss, D.J., Walsh, S.J., 2009. Remote Sensing of Mountain Environments. Geogr. Compass 3, 1–21. 10.1111/j.1749-8198.2008.00200.x

White, J.C., Coops, N.C., Wulder, M.A., Vastaranta, M., Hilker, T., Tompalski, P., 2016. Remote Sensing Technologies for Enhancing Forest Inventories: A Review. Can. J. Remote Sens. 42, 619–641. 10.1080/07038992.2016.1207484

Wilson, J.P., Gallant, J.C., 2000. Terrain Analysis: Principles and Applications. John Wiley & Sons.

Xiao, A., Xuan, W., Wang, J., Huang, J., Tao, D., Lu, S., Yokoya, N., 2025. Foundation Models for Remote Sensing and Earth Observation: A survey. IEEE Geosci. Remote Sens. Mag. 2–29. 10.1109/MGRS.2025.3576766

Xu, K., Zhang, Z., Yu, W., Zhao, P., Yue, J., Deng, Y., Geng, J., 2021. How Spatial Resolution Affects Forest Phenology and Tree-Species Classification Based on Satellite and Up-Scaled Time-Series Images. Remote Sens. 13, 2716. 10.3390/rs13142716

Zbontar, J., Jing, L., Misra, I., LeCun, Y., Deny, S., 2021. Barlow Twins: Self-Supervised Learning via Redundancy Reduction. 10.48550/arXiv.2103.03230

Zhong, L., Dai, Z., Fang, P., Cao, Y., Wang, L., 2024. A Review: Tree Species Classification Based on Remote Sensing Data and Classic Deep Learning-Based Methods. Forests 15, 852. 10.3390/f15050852

