## Supplementary Materials for "Geospatial foundation models enable data-efficient tree species mapping in temperate mountain forests"

### S.1 Study region, inventory, and preprocessing

#### S.1.1 Study region ecological details

The study was conducted in the Autonomous Province of Trento (Trentino), located in the central–eastern Italian Alps. The region encompasses highly diverse mountain landscapes, extending from valley bottoms at 200 m to high alpine peaks exceeding 3,800 m.

Trentino is dominated by Triassic carbonate platforms in the east and centre (limestones and dolomites), contrasting with Paleozoic crystalline and metamorphic rocks and large Oligocene intrusive bodies in the west. This geological mosaic exerts control on soil development, nutrient availability, drainage, and water-holding capacity, generating pronounced edaphic gradients over short distances. Carbonate substrates are typically associated with shallow, alkaline soils and strong local-scale species filtering, whereas siliceous and crystalline substrates support deeper, more acidic soils with higher moisture retention. Together with Quaternary glacial deposits in valley bottoms and lower slopes, this substrate-driven variation interacts with topography and climate to structure forest composition, productivity, and phenology across the region.

Biogeographically, Trentino lies at a transition zone between Mediterranean and Central European floras, contributing to high plant diversity and marked turnover along elevational gradients. Land use has played a major role in shaping these ecosystems: centuries of forestry, livestock grazing, and meadow management have influenced forest distribution, age structure, and species composition, leaving a fine-grained cultural landscape typical of the southern Alps.

#### S.1.2 Forest inventory characteristics

The forest inventory is described in the main text (Section 2.2.2). This section provides additional distributional and temporal detail.

#### S.1.3 Parcel statistics and species distributions

The forest inventory parcels were collected over an extended period from 2010 to 2021. Figure S.1 shows the distribution of survey effort across years.

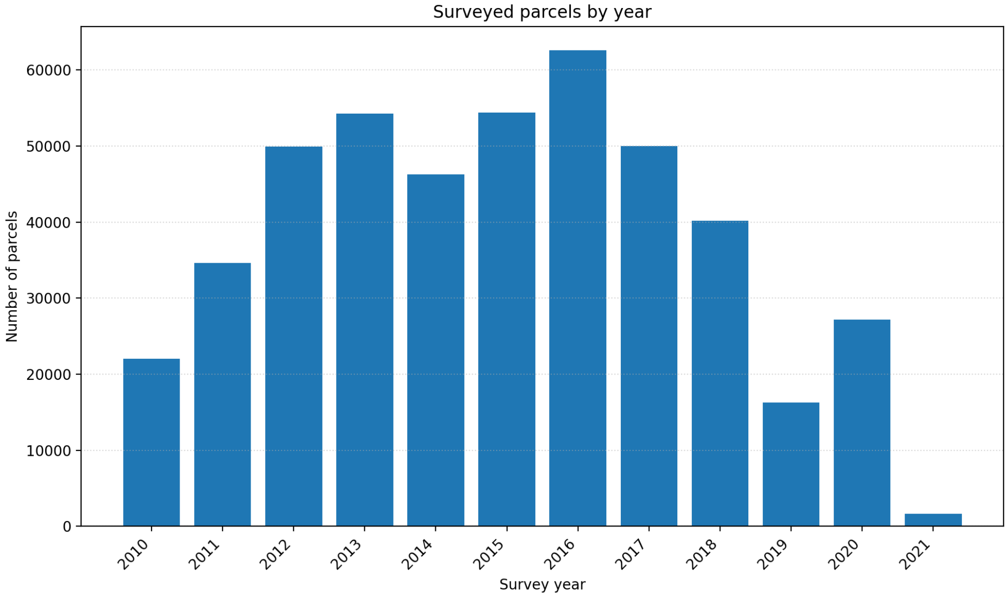

Figure S.1: Number of forest inventory parcels surveyed by year. The inventory was collected from 2010 to 2021, with survey effort concentrated in the mid-2010s.

Figure S.2 compares the area occupied by each species when computed using (i) fraction-weighted area (summing each species' proportional cover across all parcels) and (ii) dominant-only area (counting only parcels where the species has the highest cover). The disparity between these two estimates reflects the prevalence of mixed stands: species that frequently occur as secondary components (e.g. *Larix decidua*, *Abies alba*) occupy considerably more fraction-weighted area than their dominant-only counts suggest, illustrating why retaining mixed parcels in the analysis is ecologically important.

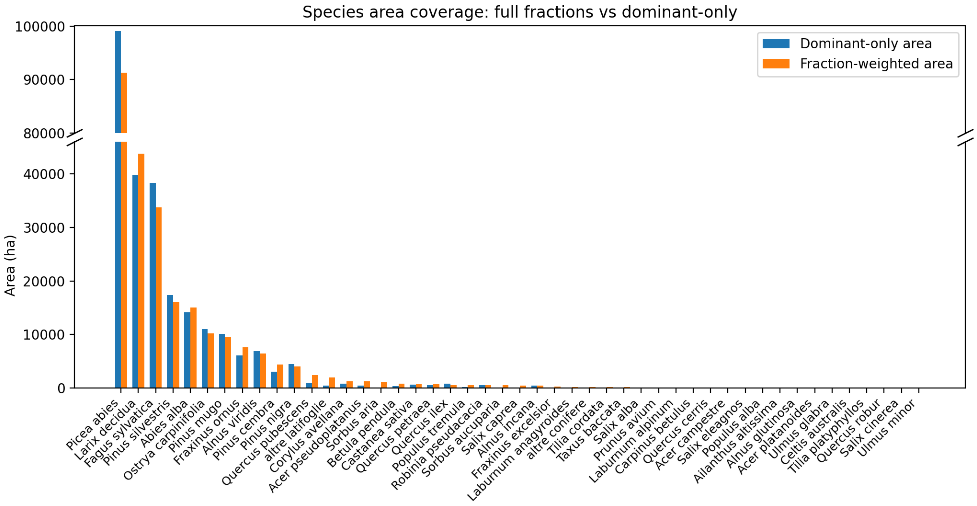

Figure S.2: Species area coverage computed from the forest inventory: dominant-only area (parcels where the species has the highest cover) versus fraction-weighted area (summing proportional cover across all parcels). The difference highlights the prevalence of mixed stands and the importance of secondary species components.

##### S.1.4 Land-cover filtering and masking

As described in the main text (Section 2.4.2), non-forest pixels were filtered using the Copernicus Global Land Cover 2020 product. Land-cover information was incorporated at two stages:

(i) **Pre-distilled label generation.** During rasterisation of forest inventory parcels, only pixels classified as forest were eligible for species labels. Non-forest pixels were assigned fixed override classes (e.g. water, sealed surfaces, snow and ice).

(ii) **Label distillation.** The same land-cover mask constrained prototype extraction, pixel-level assignment, and parcel-level proportion fitting. Non-forest pixels bypassed distillation entirely and retained their override labels.

#### S.2 Species classes and grouping decisions

The original Trentino forest inventory distinguishes approximately 45 tree species. While this taxonomic resolution is appropriate for field-based management, many of the less abundant species occur infrequently, primarily in mixed stands, and are poorly represented as dominant classes at the parcel scale. Retaining all species as independent targets would therefore lead to extreme class imbalance, unstable performance estimates, and confusion matrices dominated by sampling artefacts rather than ecologically meaningful signal.

Grouping decisions were guided by three complementary criteria:

(i) **Co-occurrence patterns at the parcel level.** Species that frequently co-occurred within the same parcels and rarely formed monospecific stands were considered candidates for grouping. Pairwise co-occurrence was quantified using Jaccard similarity and spatial correlation metrics (Table S.2).

(ii) **Shared ecological niches and habitat affinities.** Species were grouped only when they exhibit similar ecological preferences with respect to elevation, moisture regime, soil type, and disturbance context.

(iii) **Spectral–phenological similarity at satellite resolution.** At 10 m spatial resolution, species with similar leaf traits, canopy structure, and phenological dynamics are unlikely to be reliably separable using satellite-derived representations.

##### S.2.1 Original inventory species

Table S.1: Species present in the Trentino forest inventory with parcel-level summary statistics. Species are sorted by decreasing prevalence (number of parcels in which the species is recorded).

| Species | Parcels | Prevalence (%) | Mean frac. | Max frac. |
|---|---|---|---|---|
| *Picea abies* | 68,427 | 77.0 | 0.473 | 1.00 |
| *Larix decidua* | 54,655 | 61.5 | 0.250 | 1.00 |
| *Fagus sylvatica* | 38,092 | 42.8 | 0.309 | 1.00 |
| *Pinus silvestris* | 23,333 | 26.2 | 0.283 | 1.00 |
| *Abies alba* | 22,295 | 25.1 | 0.225 | 1.00 |
| *Ostrya carpinifolia* | 14,384 | 16.2 | 0.221 | 1.00 |
| *Fraxinus ornus* | 14,081 | 15.8 | 0.187 | 0.95 |
| *Corylus avellana* | 6,309 | 7.1 | 0.155 | 1.00 |
| *Pinus cembra* | 6,123 | 6.9 | 0.197 | 1.00 |
| *Acer pseudoplatanus* | 5,677 | 6.4 | 0.089 | 1.00 |
| *Quercus pubescens* | 5,640 | 6.3 | 0.130 | 0.84 |
| *Pinus nigra* | 5,398 | 6.1 | 0.339 | 1.00 |
| *Alnus alnobetula* | 5,123 | 5.8 | 0.326 | 1.00 |
| *Sorbus aria* | 5,116 | 5.8 | 0.072 | 0.60 |
| altre latifoglie | 4,485 | 5.0 | 0.113 | 1.00 |
| *Pinus mugo* | 4,371 | 4.9 | 0.458 | 1.00 |
| *Betula pendula* | 4,007 | 4.5 | 0.092 | 1.00 |
| *Castanea sativa* | 3,314 | 3.7 | 0.137 | 1.00 |
| *Populus tremula* | 3,300 | 3.7 | 0.090 | 0.95 |
| *Sorbus aucuparia* | 3,258 | 3.7 | 0.064 | 1.00 |
| *Quercus petraea* | 2,863 | 3.2 | 0.161 | 1.00 |
| *Salix caprea* | 2,592 | 2.9 | 0.101 | 1.00 |
| *Fraxinus excelsior* | 1,995 | 2.2 | 0.125 | 1.00 |
| *Robinia pseudacacia* | 1,983 | 2.2 | 0.273 | 1.00 |
| *Alnus incana* | 1,320 | 1.5 | 0.257 | 1.00 |
| *Laburnum anagyroides* | 1,220 | 1.4 | 0.090 | 1.00 |
| *Prunus avium* | 1,097 | 1.2 | 0.056 | 1.00 |
| *Laburnum alpinum* | 868 | 1.0 | 0.064 | 0.70 |
| *Tilia cordata* | 844 | 0.9 | 0.125 | 0.70 |
| *Salix alba* | 778 | 0.9 | 0.080 | 0.85 |
| *Quercus ilex* | 542 | 0.6 | 0.251 | 0.95 |
| altre conifere | 441 | 0.5 | 0.126 | 1.00 |
| *Taxus baccata* | 389 | 0.4 | 0.050 | 0.50 |
| *Acer campestre* | 341 | 0.4 | 0.055 | 0.30 |
| *Carpinus betulus* | 202 | 0.2 | 0.171 | 1.00 |
| *Quercus cerris* | 142 | 0.2 | 0.124 | 0.50 |
| *Ailanthus altissima* | 142 | 0.2 | 0.092 | 1.00 |
| *Populus alba* | 141 | 0.2 | 0.096 | 0.90 |
| *Alnus glutinosa* | 95 | 0.1 | 0.232 | 0.85 |
| *Salix eleagnos* | 86 | 0.1 | 0.217 | 0.80 |
| *Acer platanoides* | 64 | 0.1 | 0.058 | 0.20 |
| *Ulmus glabra* | 55 | 0.1 | 0.048 | 0.20 |
| *Celtis australis* | 30 | 0.0 | 0.133 | 0.80 |
| *Tilia platyphyllos* | 22 | 0.0 | 0.064 | 0.30 |
| *Quercus robur* | 13 | 0.0 | 0.114 | 0.20 |
| *Ulmus minor* | 12 | 0.0 | 0.046 | 0.05 |
| *Salix cinerea* | 10 | 0.0 | 0.160 | 0.40 |

##### S.2.2 Ecological grouping rationale

Applying the criteria above resulted in five composite species groups: (1) *Ostrya–Fraxinus ornus* thermophilous deciduous communities, which exhibited the highest pairwise co-occurrence (Jaccard = 0.68); (2) *Quercus–Carpinus* oak–hornbeam woodlands; (3) *Acer–Fraxinus–Tilia* mixed mesophilous forests; (4) Hygrophilous riparian formations; and (5) Pioneer deciduous formations.

Grey alder (*Alnus incana*) and green alder (*Alnus alnobetula*) were retained as separate classes despite their taxonomic affinity, as coexistence analysis showed they cluster with pioneer and mesophilous species respectively rather than true hygrophilous communities. Holm oak (*Quercus ilex*) was similarly retained as a distinct class, as this evergreen Mediterranean species exhibits distinct phenology.

Table S.2: Top 20 species pairs by Jaccard similarity, computed from parcel-level co-occurrence across the Trentino forest inventory.

| Species 1 | Species 2 | Co-occurrence | Jaccard | Correlation |
|---|---|---|---|---|
| *Fraxinus ornus* | *Ostrya carpinifolia* | 11,533 | 0.681 | 0.710 |
| *Larix decidua* | *Picea abies* | 48,700 | 0.655 | −0.146 |
| *Fagus sylvatica* | *Picea abies* | 28,972 | 0.374 | −0.330 |
| *Fraxinus ornus* | *Quercus pubescens* | 4,912 | 0.332 | 0.415 |
| *Ostrya carpinifolia* | *Quercus pubescens* | 4,923 | 0.326 | 0.383 |
| *Abies alba* | *Picea abies* | 21,699 | 0.314 | 0.027 |
| *Abies alba* | *Fagus sylvatica* | 13,800 | 0.296 | −0.027 |
| *Fraxinus ornus* | *Pinus silvestris* | 8,452 | 0.292 | 0.068 |
| *Fagus sylvatica* | *Larix decidua* | 20,939 | 0.292 | −0.211 |
| *Ostrya carpinifolia* | *Pinus silvestris* | 8,012 | 0.270 | 0.023 |
| *Fagus sylvatica* | *Pinus silvestris* | 12,588 | 0.258 | −0.062 |
| *Abies alba* | *Larix decidua* | 14,190 | 0.226 | −0.155 |
| *Ostrya carpinifolia* | *Sorbus aria* | 3,486 | 0.218 | 0.265 |
| *Fraxinus ornus* | *Pinus nigra* | 3,423 | 0.213 | 0.085 |
| *Fraxinus ornus* | *Sorbus aria* | 3,360 | 0.212 | 0.261 |
| *Castanea sativa* | *Quercus petraea* | 1,012 | 0.196 | 0.138 |
| *Fagus sylvatica* | *Ostrya carpinifolia* | 8,412 | 0.191 | −0.017 |
| *Picea abies* | *Pinus silvestris* | 14,614 | 0.189 | −0.283 |
| *Larix decidua* | *Pinus silvestris* | 12,357 | 0.188 | −0.160 |
| *Ostrya carpinifolia* | *Pinus nigra* | 3,081 | 0.184 | 0.042 |

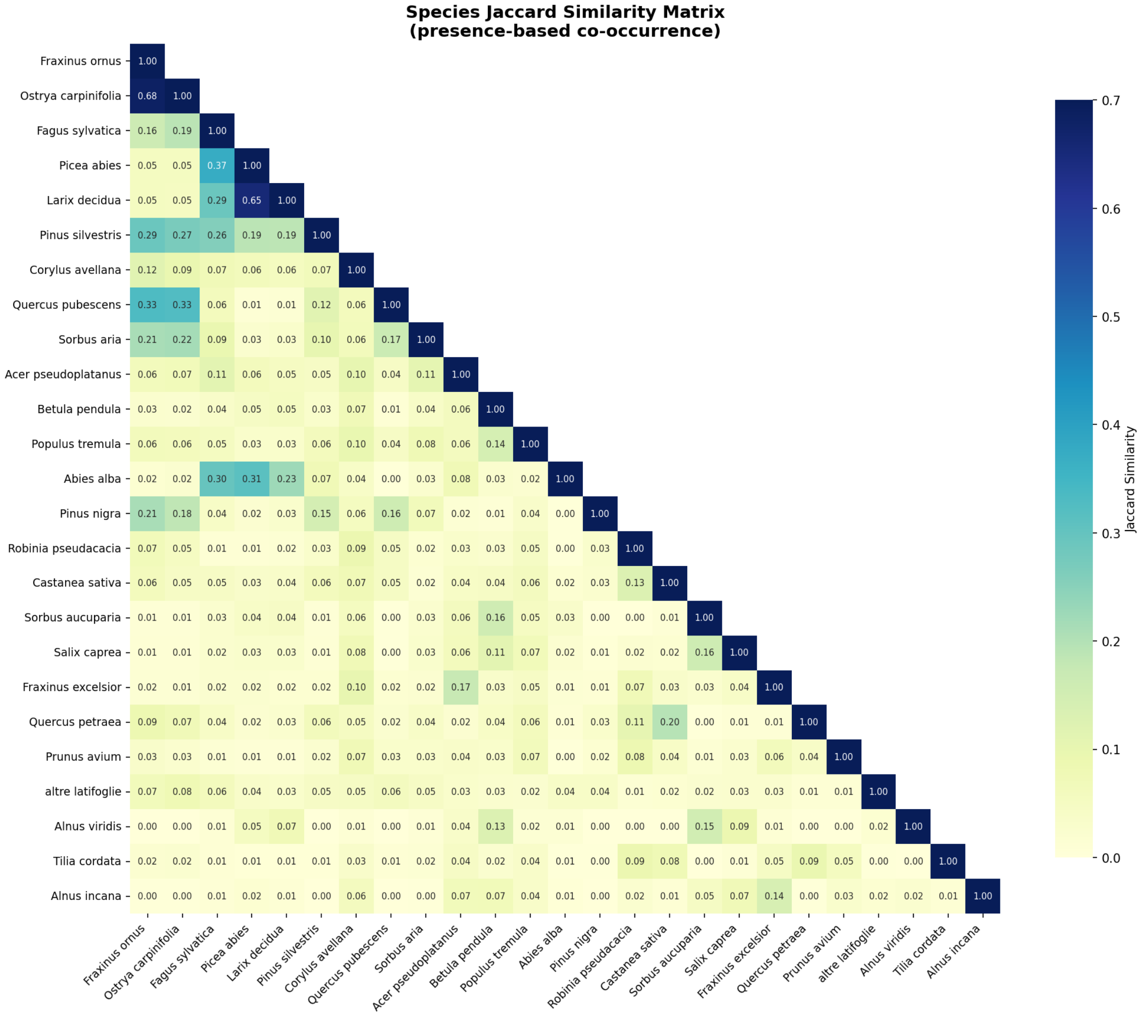

Figure S.3: Jaccard similarity heatmap for parcel-level species co-occurrence across the Trentino forest inventory. Higher values (warmer colours) indicate species pairs that frequently share the same parcels, motivating their grouping into composite classes.

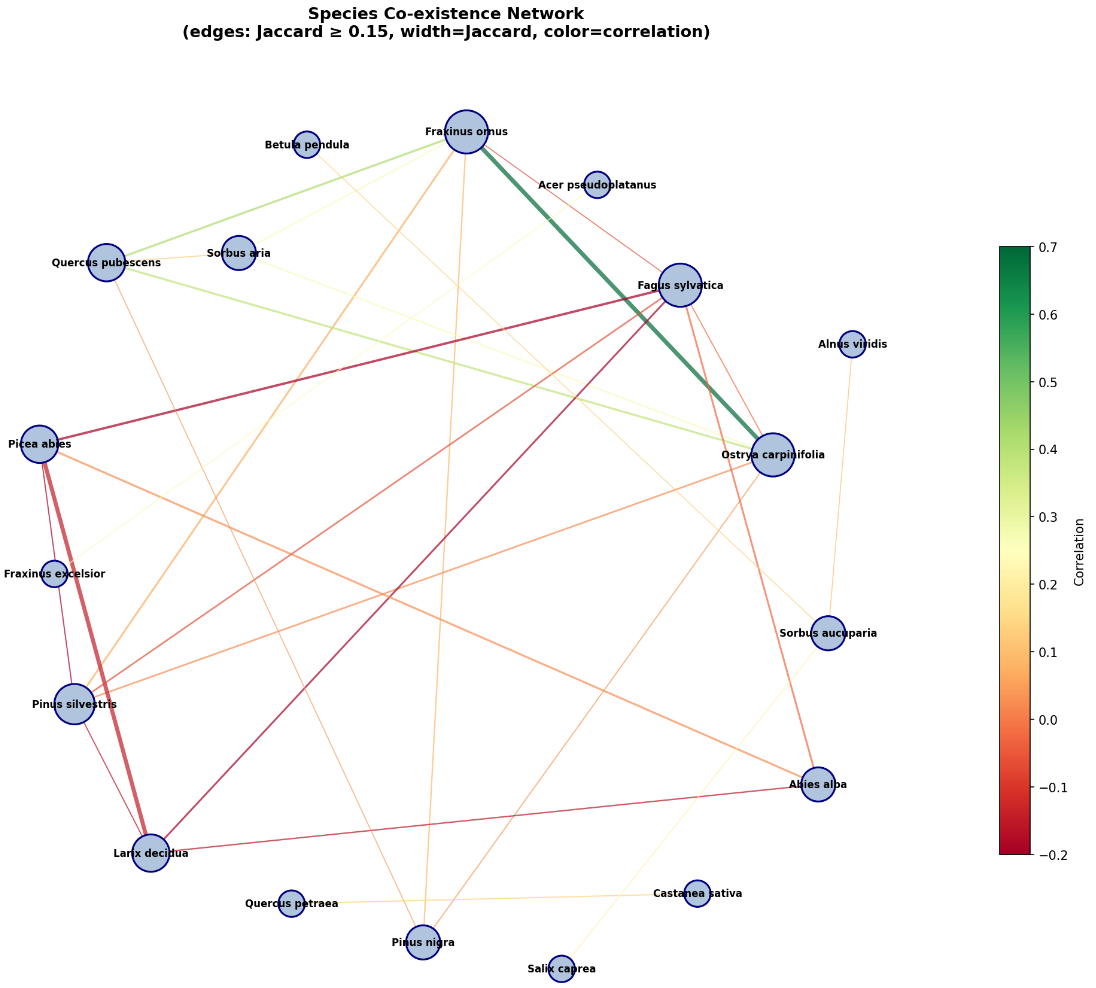

Figure S.4: Co-occurrence network of Trentino tree species. Nodes represent species and edges connect pairs with Jaccard similarity $\geq 0.15$, with edge width proportional to Jaccard similarity and edge colour indicating correlation. Tightly connected clusters informed the species grouping decisions.

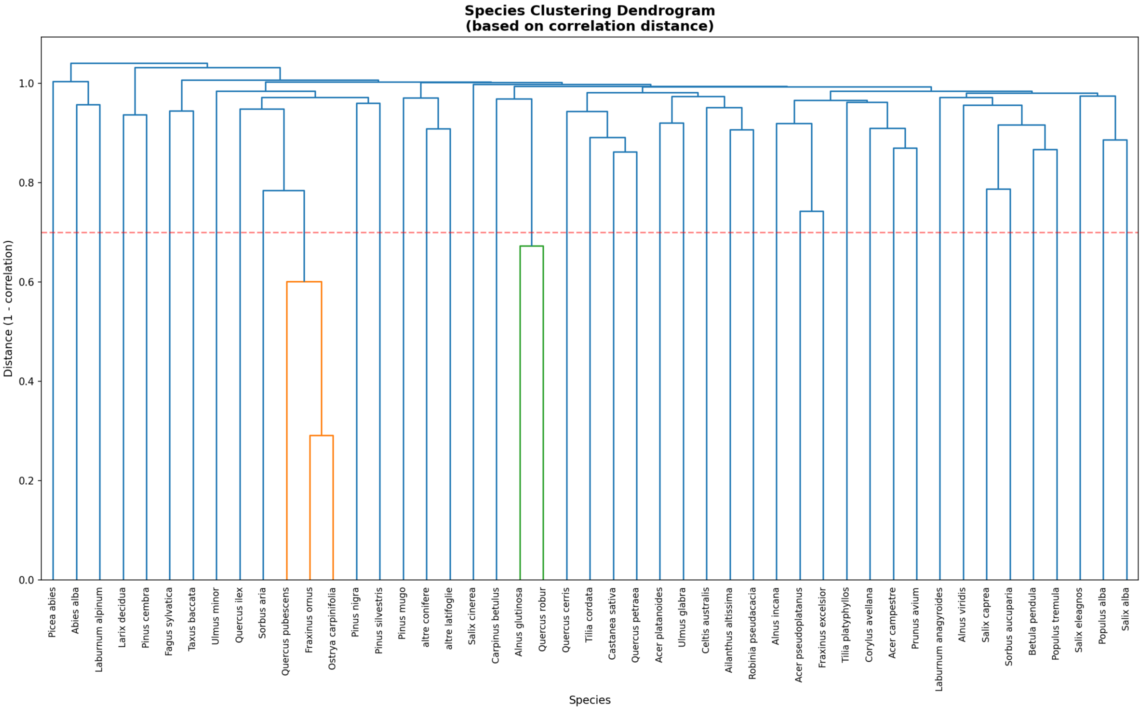

Figure S.5: Hierarchical clustering dendrogram of species co-occurrence patterns. Species that cluster together at low linkage distance share similar habitat associations and parcel-level distributions.

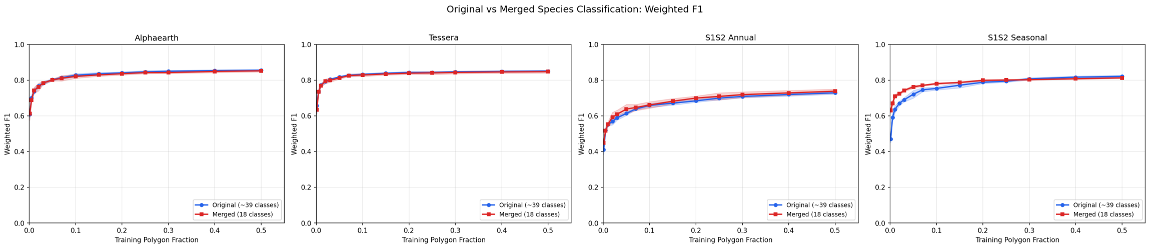

Figure S.6: Effect of species merging on parcel size distribution. Grouping minor co-occurring species into community classes increases the effective parcel count and mean parcel purity for the resulting classes, improving the quality and quantity of training labels.

##### S.2.3 Final class list and summary statistics

Table S.3 presents the 18 final target classes used in all experiments, with sample sizes and elevation distributions.

Table S.3: Final target classes with training sample sizes and elevation distributions. The classification uses 18 classes comprising 13 dominant species retained as individual classes and 5 community groups formed by merging ecologically co-occurring minor species.

| Class | Common name | Parcels | Pixels | Elev. range (m) | Elev. mean (m) |
|---|---|---|---|---|---|
| *Dominant species* | | | | | |
| *Picea abies* | Norway spruce | 33,885 | 9,555,996 | 151–2849 | 1482 |
| *Fagus sylvatica* | European beech | 11,991 | 3,540,217 | 130–2008 | 1160 |
| *Larix decidua* | European larch | 11,274 | 3,515,723 | 200–2611 | 1700 |
| *Pinus silvestris* | Scots pine | 6,255 | 1,559,357 | 76–2794 | 1011 |
| *Abies alba* | Silver fir | 4,464 | 1,406,263 | 245–2192 | 1297 |
| *Pinus mugo* | Mountain pine | 1,887 | 841,517 | 318–2366 | 1715 |
| *Pinus nigra* | Black pine | 1,832 | 404,008 | 73–1802 | 705 |
| *Alnus alnobetula* | Green alder | 1,680 | 381,005 | 815–2561 | 1910 |
| *Pinus cembra* | Swiss stone pine | 772 | 279,831 | 1439–2455 | 1980 |
| *Corylus avellana* | Common hazel | 662 | 65,432 | 211–2205 | 1087 |
| *Robinia pseudacacia* | Black locust | 447 | 52,813 | 201–1322 | 690 |
| *Alnus incana* | Grey alder | 299 | 37,140 | 468–2160 | 1327 |
| *Quercus ilex* | Holm oak | 120 | 57,819 | 63–1303 | 388 |
| *Community groups* | | | | | |
| Ostrya-Fraxinus ornus[a] | Thermophilous deciduous | 6,304 | 2,095,415 | 72–1914 | 790 |
| Quercus-Carpinus[b] | Oak-hornbeam woodland | 692 | 114,640 | 152–1459 | 795 |
| Pioneer deciduous[c] | Early successional | 610 | 81,989 | 223–2131 | 1374 |
| Acer-Fraxinus-Tilia[d] | Mixed mesophilous | 444 | 59,162 | 139–1864 | 1092 |
| Hygrophilous[e] | Riparian/wetland | 66 | 6,060 | 165–2068 | 1097 |
| **Total** | (18 classes) | **83684** | **24054387** | | |

[a] *Ostrya carpinifolia*, *Fraxinus ornus*, *Quercus pubescens*, *Sorbus aria*, *Acer campestre*, *Ulmus minor*, *Celtis australis*
[b] *Carpinus betulus*, *Castanea sativa*, *Prunus avium*, *Quercus cerris*, *Q. petraea*, *Q. robur*
[c] *Betula pendula*, *Populus tremula*, *Salix caprea*, *Sorbus aucuparia*, *Ailanthus altissima*, *Laburnum alpinum*, *L. anagyroides*
[d] *Acer platanoides*, *A. pseudoplatanus*, *Fraxinus excelsior*, *Taxus baccata*, *Tilia cordata*, *T. platyphyllos*, *Ulmus glabra*
[e] *Alnus glutinosa*, *Populus alba*, *Salix alba*, *S. cinerea*, *S. eleagnos*

#### S.3 Satellite data preprocessing details

The satellite data acquisition and compositing strategy is described in the main text (Section 2.3). This section provides additional technical detail on the preprocessing implementation.

##### S.3.1 Sentinel-2 pixel-level masking

Imagery was obtained from the `COPERNICUS/S2_SR_HARMONIZED` collection in Google Earth Engine. Pixel-level masking used the Sen2Cor Scene Classification Layer (SCL) and QA60 cloud bitmask jointly: pixels labelled as cloud shadow, medium or high probability cloud, thin cirrus, snow/ice, saturated or defective, or no-data were excluded, while vegetation, bare soil, water, and unclassified pixels were retained. QA60 bits corresponding to opaque cloud and cirrus contamination were additionally required to be unset.

##### S.3.2 Sentinel-1 compositing in linear space

Imagery was obtained from the `COPERNICUS/S1_GRD` collection (IW mode, dual VV/VH polarisation). The GRD collection is radiometrically calibrated and terrain corrected, with backscatter provided in decibels (dB). To avoid bias from aggregating in logarithmic space, compositing was performed in linear power units: VV and VH values were converted from dB to linear, per-pixel medians were computed within each temporal window, and the results were converted back to dB. Two polarisation-derived features were additionally computed: VV−VH (in dB) and VV/VH ratio (in linear space). The incidence angle band was retained.

##### S.3.3 Grid alignment

All outputs were exported on a fixed 10 m UTM grid aligned exactly to the study region. Sentinel-2 bands at 20 m and 60 m were resampled to 10 m using nearest-neighbour assignment; Sentinel-1 features were exported on the same grid.

##### S.3.4 Reproducibility

All preprocessing, compositing, and export steps were implemented in Google Earth Engine. The full script is included in the project repository: `https://github.com/PatBall1/trentino-trees`.

#### S.4 Label distillation algorithm

Label distillation converts parcel-level species fractions into pixel-level pseudo-labels by combining inventory constraints with foundation model (FM) embeddings. The algorithm proceeds as follows.

##### S.4.1 Inputs

- Parcel polygons with species fraction columns $f_{p,s}$ indicating the proportion of species $s$ in parcel $p$ (fractions sum to 1 within each parcel).
- FM pixel embeddings $\mathbf{x}_i$ for each valid pixel $i$.
- Land-cover mask for non-forest exclusion (optional).

##### S.4.2 Stage 1: Anchor extraction

Parcels with dominant fraction $\geq \tau_0$ (default 0.90) are treated as anchors. For each anchor parcel:

1. All enclosed pixels receive the dominant species label directly.
2. These pixels form an exemplar set for each species $s$.

From the exemplars, we construct:

- A $k$-NN index over all anchor pixels for local similarity queries.
- Species prototypes $\boldsymbol{\mu}_s$: mean embeddings per species (optionally, $K$ prototypes via $k$-means clustering to handle within-species variability).

##### S.4.3 Stage 2: Mixed parcel assignment

Mixed parcels are processed in descending purity brackets using an annealing schedule (Table S.5).

For each parcel $p$ with $n_p$ pixels:

1. Identify allowed species: $\mathcal{S}_p = \{s : f_{p,s} > \epsilon\}$, where $\epsilon$ is a minimum fraction threshold (default 0.001).

2. Compute hybrid probabilities. For each pixel $i$, blend $k$-NN and prototype similarities:
$$P(s \mid \mathbf{x}_i) = \alpha \cdot P_{\text{kNN}}(s \mid \mathbf{x}_i) + (1-\alpha) \cdot P_{\text{proto}}(s \mid \mathbf{x}_i) \tag{S.1}$$
where $\alpha = 0.7$ (empirically tuned), $P_{\text{kNN}}$ is the fraction of $k$ nearest exemplars belonging to species $s$, and $P_{\text{proto}}$ is the softmax over cosine similarities to species prototypes at temperature $T$.

3. Enforce parcel proportions via iterative proportional fitting (IPF): rescale per-pixel probabilities so that
$$\sum_{i \in p} P(s \mid \mathbf{x}_i) \approx n_p \cdot f_{p,s} \quad \forall\, s \in \mathcal{S}_p \tag{S.2}$$
IPF iteratively adjusts row and column scaling factors until convergence (typically <10 iterations).

4. Assign labels: each pixel receives the species with maximum adjusted probability. Confidence scores (max probability) are recorded.

5. Update prototypes (optional): incorporate newly labelled high-confidence pixels using exponential moving average with momentum 0.8.

##### S.4.4 Hyperparameters

Table S.4: Default hyperparameters for label distillation.

| Parameter | Default | Description |
|---|---|---|
| Pure threshold $\tau_0$ | 0.90 | Minimum dominant fraction for anchor parcels |
| $k$-NN neighbours | 15 | Neighbours for exemplar voting |
| $k$-NN blend $\alpha$ | 0.70 | Weight for $k$-NN vs. prototype |
| Prototypes per species $K$ | 3 | Number of prototype clusters |
| Initial temperature | 1.0 | Softmax sharpness |
| Temperature decay | 0.85 | Per-stage multiplicative decay |
| Minimum temperature | 0.30 | Floor for temperature annealing |
| Prototype momentum | 0.80 | EMA weight for prototype updates |
| IPF tolerance | 0.01 | Convergence threshold for proportion fitting |

Table S.5: Annealing schedule for mixed parcel distillation. Purity brackets are processed in descending order; the temperature is decayed multiplicatively at each stage.

| Stage | Purity range | Temperature $T$ |
|---|---|---|
| 1 | 0.90–0.85 | 1.00 |
| 2 | 0.85–0.75 | 0.85 |
| 3 | 0.75–0.65 | 0.72 |
| 4 | 0.65–0.55 | 0.61 |
| ... | ... | $T \leftarrow T \times 0.85$ (min 0.30) |

##### S.4.5 Outputs

Three co-registered GeoTIFFs:

- **Class raster**: distilled species ID per pixel.
- **Parcel raster**: source parcel ID for provenance tracking.
- **Fraction raster**: dominant fraction of the source parcel (proxy for label confidence).

##### S.4.6 Distillation tuning results

We performed a grid search over distillation hyperparameters separately for AlphaEarth (GSE) and Tessera representations. Table S.6 reports the best configuration for each, and Table S.7 shows the effect of the minimum purity parameter, which was the most impactful hyperparameter for the GSE representation.

Table S.6: Best distillation configurations for GSE and Tessera representations, selected by macro F1 on validation data.

| Metric | GSE 2018 | Tessera 2018 |
|---|---|---|
| Macro F1 | 0.586 | 0.425 |
| Weighted F1 | 0.879 | 0.819 |
| Accuracy | 0.876 | 0.807 |
| Min purity | 0.55 | 0.4 |
| Anneal thresholds | 0.85,0.75,0.65,0.55 | 0.85,0.75,0.65,0.55 |
| Temperature decay | 0.85 | 0.85 |
| Update confidence | 0.7 | 0.6 |

Table S.7: Effect of minimum purity threshold on distillation performance (GSE 2018 representation). Higher purity thresholds filter out more ambiguous labels, improving downstream classifier quality.

| Min purity | Mean macro F1 | Max macro F1 | $n$ configs |
|---|---|---|---|
| 0.40 | 0.492 | 0.512 | 25 |
| 0.45 | 0.506 | 0.533 | 24 |
| 0.50 | 0.528 | 0.562 | 24 |
| 0.55 | 0.566 | 0.586 | 16 |
| 0.60 | 0.576 | 0.586 | 12 |
| 0.65 | 0.576 | 0.586 | 12 |
| 0.70 | 0.576 | 0.586 | 12 |

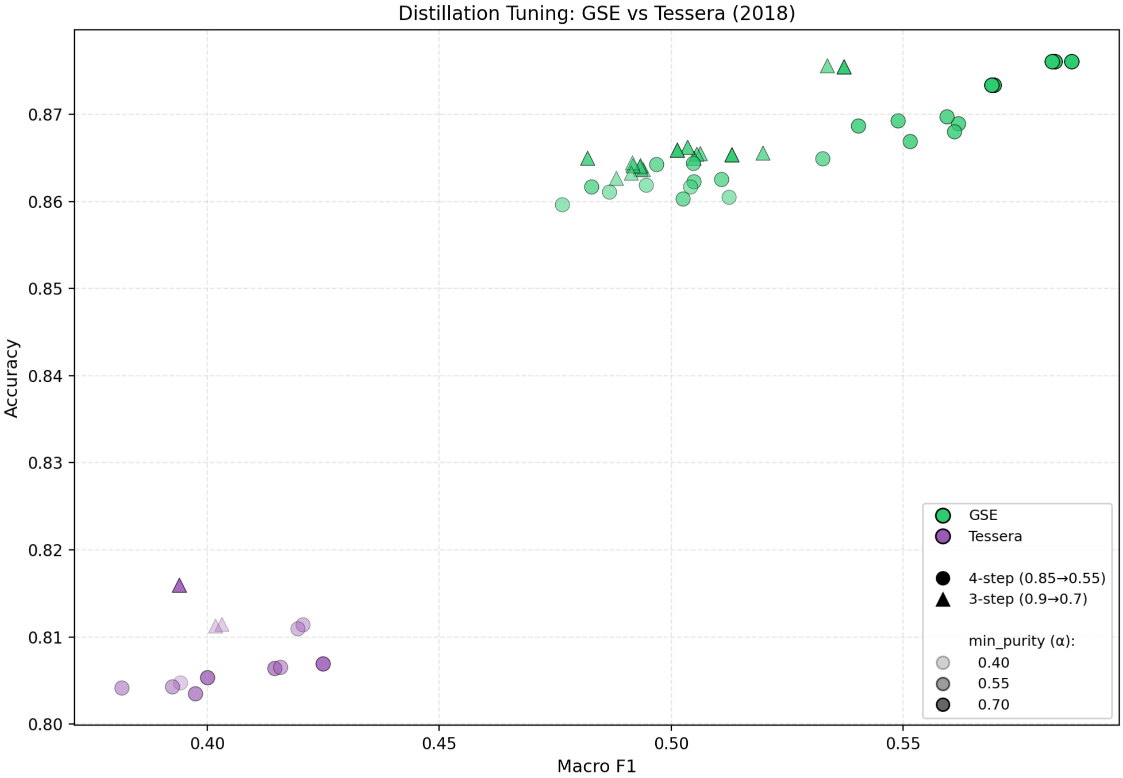

Figure S.7: Effect of distillation hyperparameters on downstream classification performance. Each point represents one configuration from the grid search; the x-axis shows macro F1 and the y-axis shows overall accuracy.

Figure S.8 compares per-species pixel area from pre-distilled labels (dominant species only) and distilled labels (refined using parcel-level proportions). Distillation redistributes area from the most dominant species toward secondary species, producing a species distribution closer to the fraction-weighted inventory estimates.

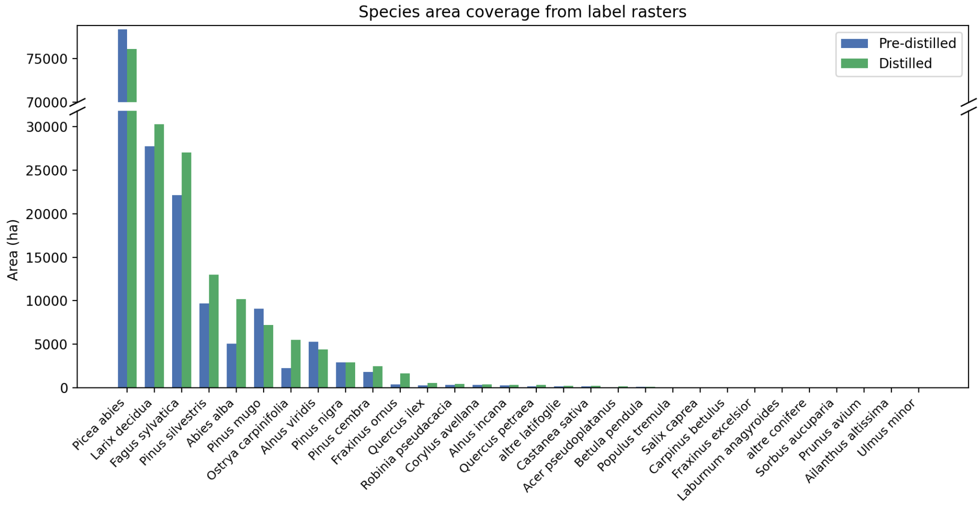

Figure S.8: Species area coverage from label rasters before (pre-distilled) and after (distilled) label distillation. Distillation redistributes pixel area from dominant species toward secondary species, producing a species distribution that better reflects the fraction-weighted inventory proportions.

#### S.5 Model training and hyperparameter tuning

##### S.5.1 Random Forest hyperparameter grid

Random Forest hyperparameter tuning was performed using an exhaustive grid search across 108 parameter combinations for each of four embedding representations. The parameter grid explored:

- `n_estimators`: 64, 128, 256
- `max_depth`: 20, 40, unlimited
- `max_features`: sqrt, all
- `min_samples_leaf`: 1, 2, 5
- `class_weight`: None, balanced

Training data was filtered using a minimum purity threshold of 0.6, while validation data required a stricter threshold of 0.8 to ensure high-quality labels.

##### S.5.2 Class balancing and purity distribution

The species classes in the Trentino inventory are highly imbalanced: the three most abundant species (*Picea abies*, *Larix decidua*, and *Fagus sylvatica*) collectively account for over 69% of all labelled pixels. To mitigate this during model training, parcel-aware downsampling was applied: the pixel counts of the three most abundant species were reduced within each training fold to match the fourth most abundant species, while distributing retained pixels across as many parcels as possible to preserve spatial and environmental variability. Figure S.9 shows the distribution of training pixels and parcels across purity bins, illustrating the trade-off between label quality and data volume that motivates the purity-threshold experiments in this study.

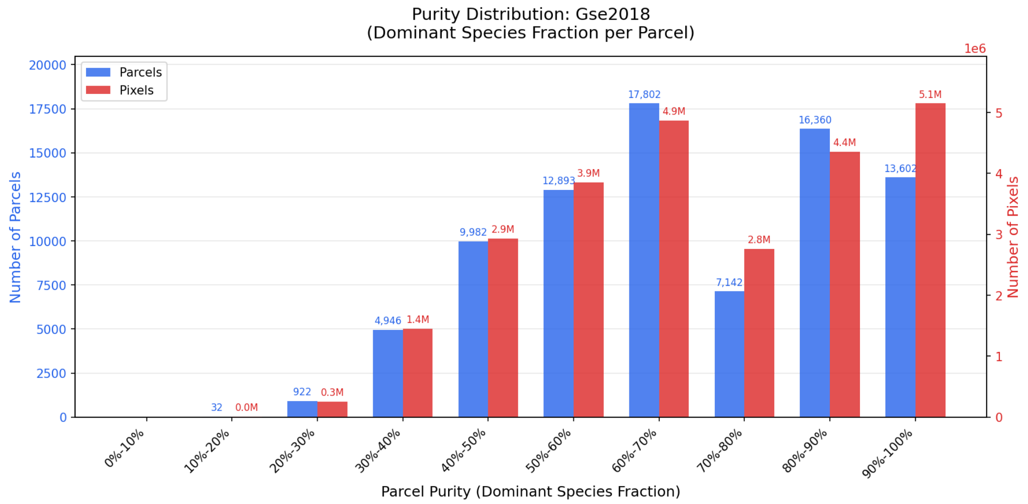

Figure S.9: Distribution of pixels and parcels across dominant-species purity bins. Most parcels have moderate purity (50–80%), with relatively few pure monospecific stands (≥90%). Restricting training to high-purity parcels substantially reduces the available training data.

##### S.5.3 Baseline classifier comparison

Table S.8 summarises the best performance achieved by each baseline classifier across the four input representations.

Table S.8: Baseline model comparison across input representations. Performance metrics (mean across 5 folds) for Logistic Regression (LR), *k*-Nearest Neighbours (KNN), and Random Forest (RF).

| Embedding | Model | Weighted F1 | Macro F1 | Prop. L1 |
|---|---|---|---|---|
| Tessera | LR | 0.717 | 0.376 | 0.465 |
| | KNN | 0.769 | 0.455 | 0.358 |
| | RF | 0.779 | 0.467 | 0.379 |
| AlphaEarth | LR | 0.714 | 0.363 | 0.520 |
| | KNN | 0.762 | 0.430 | 0.380 |
| | RF | 0.800 | 0.470 | 0.401 |
| S1S2 seasonal | LR | 0.651 | 0.296 | 0.522 |
| | KNN | 0.720 | 0.364 | 0.386 |
| | RF | 0.790 | 0.454 | 0.379 |
| S1S2 annual | LR | 0.532 | 0.219 | 0.604 |
| | KNN | 0.659 | 0.308 | 0.422 |
| | RF | 0.724 | 0.397 | 0.411 |

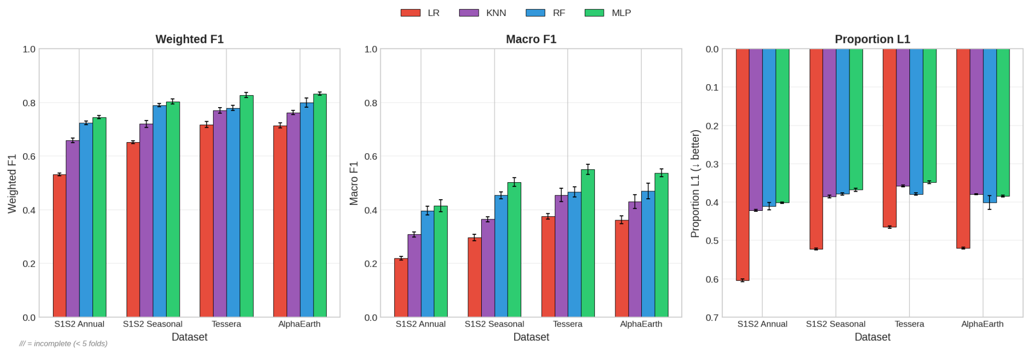

Figure S.10: Comparison of baseline classifier performance across input representations. Bar heights show mean weighted F1 and macro F1 across 5 folds for Logistic Regression, $k$-NN, and Random Forest.

##### S.5.4 MLP nested cross-validation results

Table S.9 reports the full MLP nested cross-validation results for all representation–configuration combinations, including DEM augmentation and distillation variants.

Table S.9: MLP classification performance across input representations and configurations using 5-fold nested cross-validation. Values shown are mean ± std across outer folds.

| Configuration | Weighted F1 | Macro F1 | Prop. L1 |
|---|---|---|---|
| Tessera | 0.827 ± 0.009 | 0.551 ± 0.019 | 0.349 ± 0.004 |
| AlphaEarth | 0.833 ± 0.006 | 0.537 ± 0.015 | 0.384 ± 0.002 |
| Tessera + DEM | 0.827 ± 0.008 | 0.548 ± 0.021 | 0.350 ± 0.003 |
| AlphaEarth + DEM | 0.831 ± 0.005 | 0.540 ± 0.018 | 0.384 ± 0.004 |
| Tessera (distilled) | 0.816 ± 0.007 | 0.548 ± 0.015 | 0.358 ± 0.005 |
| AlphaEarth (distilled) | 0.817 ± 0.007 | 0.526 ± 0.016 | 0.390 ± 0.003 |
| S1S2 seasonal | 0.803 ± 0.010 | 0.503 ± 0.017 | 0.368 ± 0.004 |
| S1S2 annual | 0.745 ± 0.005 | 0.414 ± 0.022 | 0.401 ± 0.002 |

#### S.6 Soft label training

A central finding of this study is that replacing hard dominant-species labels with soft parcel-proportion labels substantially improves classification performance, particularly for minority species. This section provides additional details on the soft-label formulation, controlled comparisons across training regimes, and per-class analysis of where gains are concentrated.

##### S.6.1 Soft-label formulation

For each pixel, the training target is a probability distribution where each class receives weight proportional to its fractional coverage within the parent parcel. Formally, a pixel within a parcel comprising 70% Norway spruce and 30% European larch is assigned target probabilities of 0.70 and 0.30 for these classes respectively.

The model is trained using cross-entropy loss generalised to soft targets:

$$\mathcal{L} = -\sum_{c=1}^{C} y_c \log \hat{y}_c \tag{S.3}$$

where $y_c$ is the target probability (parcel fraction) for class $c$ and $\hat{y}_c$ is the predicted probability from the softmax output. This loss is equivalent to the Kullback–Leibler (KL) divergence between target and predicted distributions (up to an additive constant) and encourages the model to output calibrated probabilities that, when aggregated across parcels, better approximate true species proportions. The motivation for this formulation is discussed in the main text (Section 4.4).

##### S.6.2 Comparison of label training strategies

Table S.10: Classification performance comparison across three label training strategies using Tessera embeddings with 5-fold cross-validation. Hard labels assign the dominant species to all pixels; soft labels use parcel-level species proportions as targets; purity-adaptive filtering restricts training to higher-purity parcels.

| Method | Accuracy | Weighted F1 | Macro F1 | Prop. L1 |
|---|---|---|---|---|
| Hard labels | 0.822 ± 0.012 | 0.831 ± 0.011 | 0.543 ± 0.023 | 0.894 ± 0.002 |
| Soft labels | 0.834 ± 0.007 | 0.838 ± 0.008 | 0.534 ± 0.018 | 0.889 ± 0.002 |
| Purity-adaptive | 0.830 ± 0.010 | 0.836 ± 0.009 | 0.541 ± 0.024 | 0.891 ± 0.001 |

##### S.6.3 Performance across purity thresholds

To quantify how soft-label training interacts with training-data quality, we compare performance at each purity threshold under both hard-label and soft-label regimes. Both training strategies show stable performance across the 0.3–0.8 purity range, with a sharp degradation at thresholds above 0.9 where training data volume collapses. Soft labels consistently achieve higher macro F1 ($\sim$0.58 vs. $\sim$0.57 for hard labels) across this plateau, while the gap widens at low purity thresholds where hard labels assign more incorrect dominant-species labels to mixed parcels.

Table S.11: Performance comparison of hard-label and soft-label training across purity thresholds (Tessera embeddings, MLP classifier). Both regimes are stable from 0.3 to 0.8 purity before degrading sharply at $\geq$0.9 as training data becomes scarce. Soft labels achieve consistently higher macro F1 across all thresholds. Values are mean across 15 runs (3 seeds $\times$ 5 folds).

| | Hard labels | | | Soft labels | | |
|---|---|---|---|---|---|---|
| Purity | W-F1 | M-F1 | Prop. L1 | W-F1 | M-F1 | Prop. L1 |
| 0.30 | 0.836 | 0.566 | 0.343 | 0.851 | 0.586 | 0.335 |
| 0.40 | 0.837 | 0.569 | 0.343 | 0.852 | 0.586 | 0.336 |
| 0.50 | 0.839 | 0.574 | 0.344 | 0.851 | 0.584 | 0.337 |
| 0.60 | 0.843 | 0.577 | 0.344 | 0.851 | 0.582 | 0.341 |
| 0.70 | 0.841 | 0.577 | 0.354 | 0.847 | 0.577 | 0.350 |
| 0.80 | 0.842 | 0.581 | 0.355 | 0.846 | 0.571 | 0.351 |
| 0.90 | 0.813 | 0.459 | 0.374 | 0.821 | 0.455 | 0.371 |
| 0.95 | 0.795 | 0.423 | 0.389 | 0.806 | 0.448 | 0.386 |
| 1.00 | 0.796 | 0.423 | 0.388 | 0.805 | 0.449 | 0.386 |

Table S.12: Performance comparison of hard-label and soft-label training across purity thresholds (AlphaEarth embeddings, MLP classifier). Soft labels show a larger macro F1 advantage than with Tessera embeddings, particularly at low-to-moderate purity thresholds. Values are mean across 15 runs (3 seeds $\times$ 5 folds).

| | Hard labels | | | Soft labels | | |
|---|---|---|---|---|---|---|
| Purity | W-F1 | M-F1 | Prop. L1 | W-F1 | M-F1 | Prop. L1 |
| 0.30 | 0.836 | 0.550 | 0.377 | 0.857 | 0.584 | 0.368 |
| 0.40 | 0.838 | 0.551 | 0.376 | 0.857 | 0.580 | 0.369 |
| 0.50 | 0.841 | 0.560 | 0.376 | 0.857 | 0.589 | 0.370 |
| 0.60 | 0.846 | 0.565 | 0.376 | 0.856 | 0.581 | 0.374 |
| 0.70 | 0.848 | 0.564 | 0.387 | 0.854 | 0.572 | 0.382 |
| 0.80 | 0.846 | 0.561 | 0.388 | 0.852 | 0.565 | 0.385 |
| 0.90 | 0.818 | 0.448 | 0.409 | 0.827 | 0.451 | 0.404 |
| 0.95 | 0.794 | 0.409 | 0.428 | 0.811 | 0.446 | 0.421 |
| 1.00 | 0.792 | 0.409 | 0.431 | 0.807 | 0.441 | 0.423 |

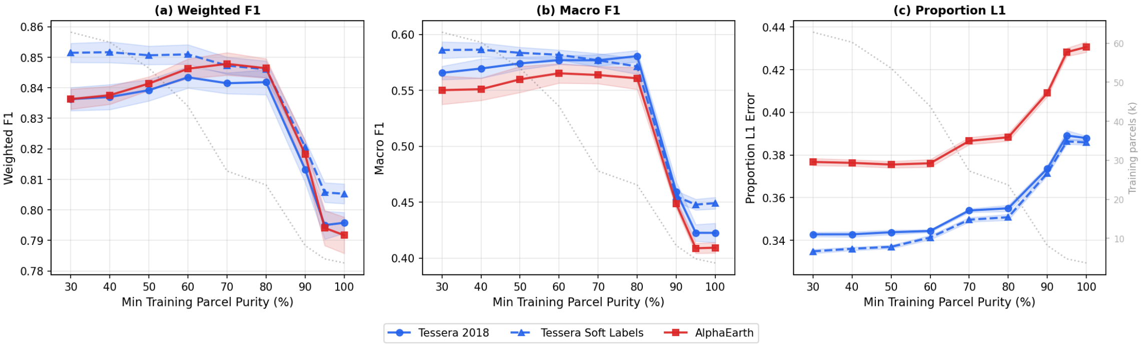

Figure S.11: Purity curve comparison of key classification metrics under hard-label and soft-label training regimes (Tessera and AlphaEarth embeddings, MLP classifier). Both regimes show a stable plateau from 0.3 to 0.8 purity, with a sharp drop above 0.9 where training data volume collapses. Soft labels achieve consistently higher macro F1 across all thresholds for both embeddings.

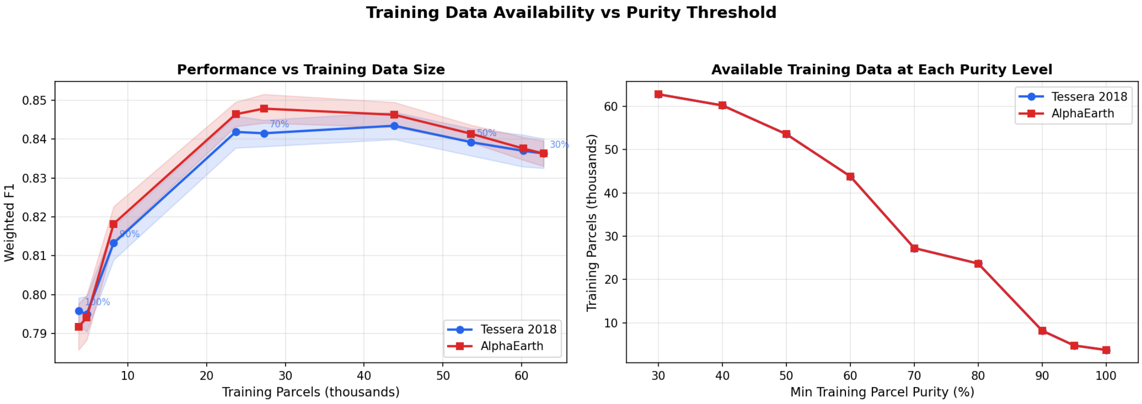

Figure S.12: Training data volume as a function of purity threshold. As the purity filter is tightened (moving right), the number of training parcels and pixels drops sharply. Under hard labels this trade-off limits the accessible performance envelope; soft labels remove this constraint.

#### S.7 Additional results and diagnostics

##### S.7.1 Fraction curve summary

Table S.13 reports macro F1 as a function of training data fraction for each input representation (corresponding to the label efficiency curves presented in the main text). Tessera embeddings achieve the highest macro F1 at every data fraction. The ranking is less clear-cut for proportion L1, where S1S2 seasonal composites match or outperform AlphaEarth.

Table S.13: Label efficiency: macro F1 as a function of training data fraction across input representations. Values are mean across 15 runs (3 seeds $\times$ 5 folds). 100% training data corresponds to approximately 45,200 parcels.

| Fraction | Tessera | AlphaEarth | S1S2 seasonal | S1S2 annual |
|---|---|---|---|---|
| 0.001 | 0.263 | 0.219 | 0.184 | 0.150 |
| 0.005 | 0.332 | 0.315 | 0.254 | 0.204 |
| 0.010 | 0.375 | 0.332 | 0.282 | 0.237 |
| 0.050 | 0.447 | 0.434 | 0.379 | 0.316 |
| 0.100 | 0.477 | 0.457 | 0.398 | 0.343 |
| 0.500 | 0.550 | 0.534 | 0.490 | 0.416 |
| 1.000 | 0.577 | 0.565 | 0.520 | 0.432 |

##### S.7.2 Embedding space visualisation

The main text (Figure 6) visualises Tessera embeddings coloured by the 18 final classes (merged species groups). To complement that view, the figures below show UMAP projections of both Tessera and AlphaEarth embeddings coloured by individual species (pre-merge), genus, conifer/broadleaf functional type, and elevation, revealing finer-grained structure in the embedding space. Point ordering uses the third UMAP dimension (plotted back-to-front) to convey depth.

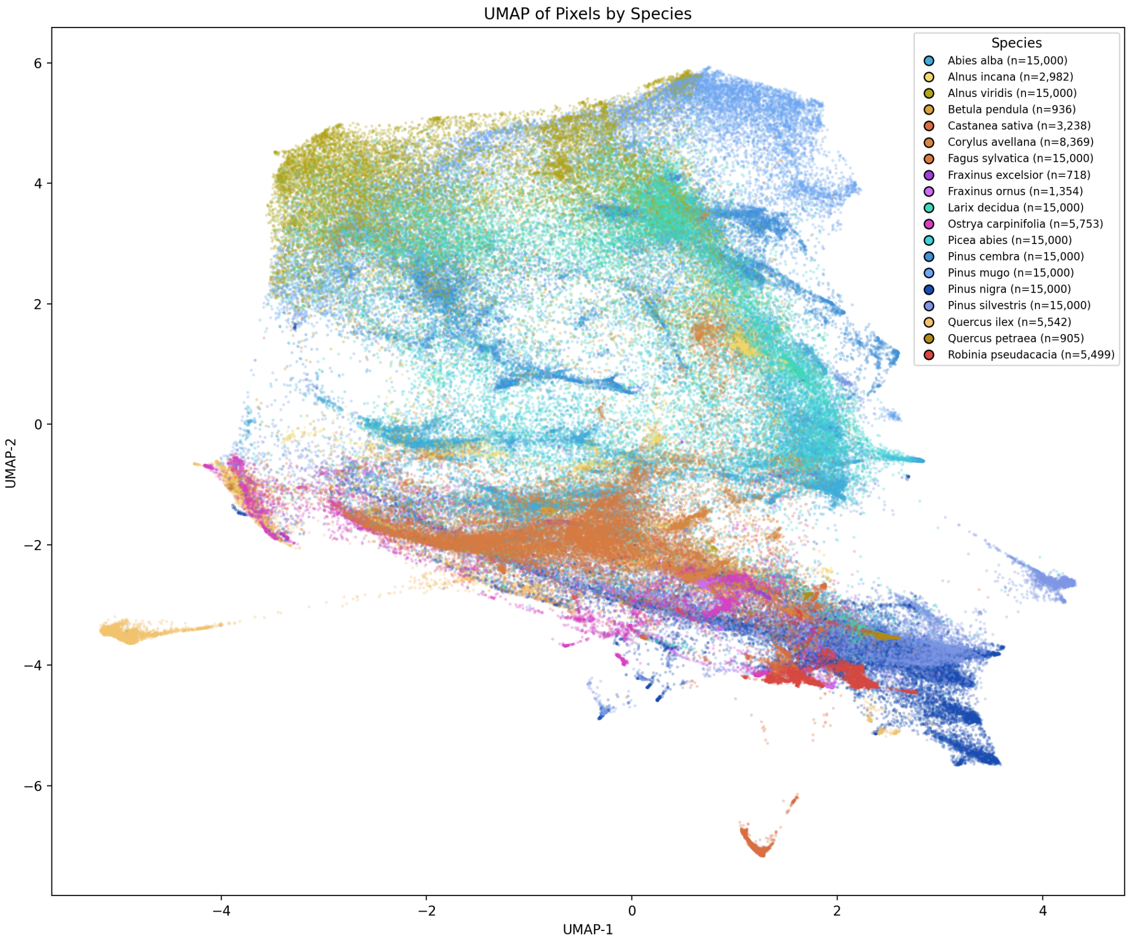

Figure S.13: UMAP projection of Tessera 2018 pixel embeddings, coloured by individual species (pre-merge). Species that are merged into community groups in the main-text classification (e.g. *Ostrya carpinifolia* and *Fraxinus ornus*) are shown separately here, demonstrating that the foundation model resolves species-level structure even among co-occurring taxa.

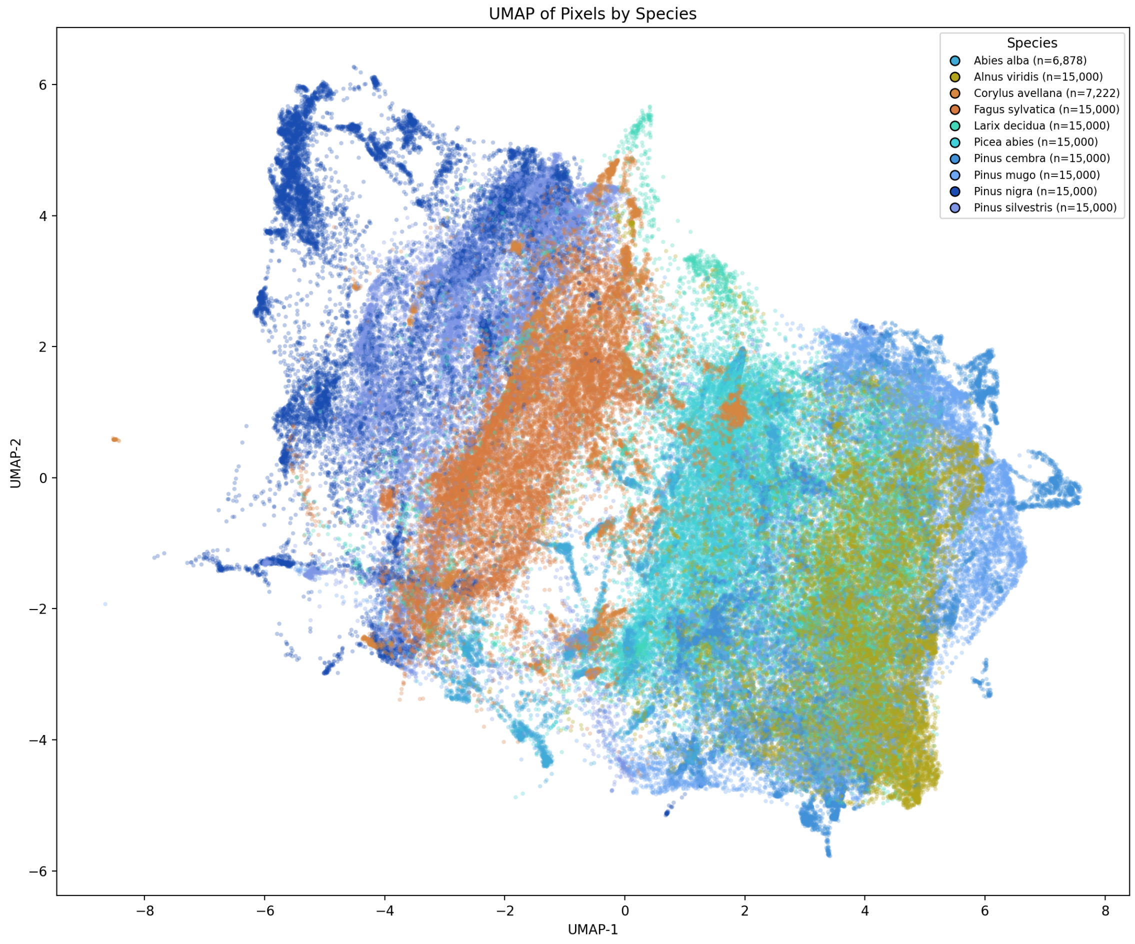

Figure S.14: UMAP projection of AlphaEarth (GSE) 2018 pixel embeddings, coloured by individual species (pre-merge). AlphaEarth shows broadly similar clustering structure to Tessera (Figure S.13), though with fewer minority species reaching the sample-size threshold for display.

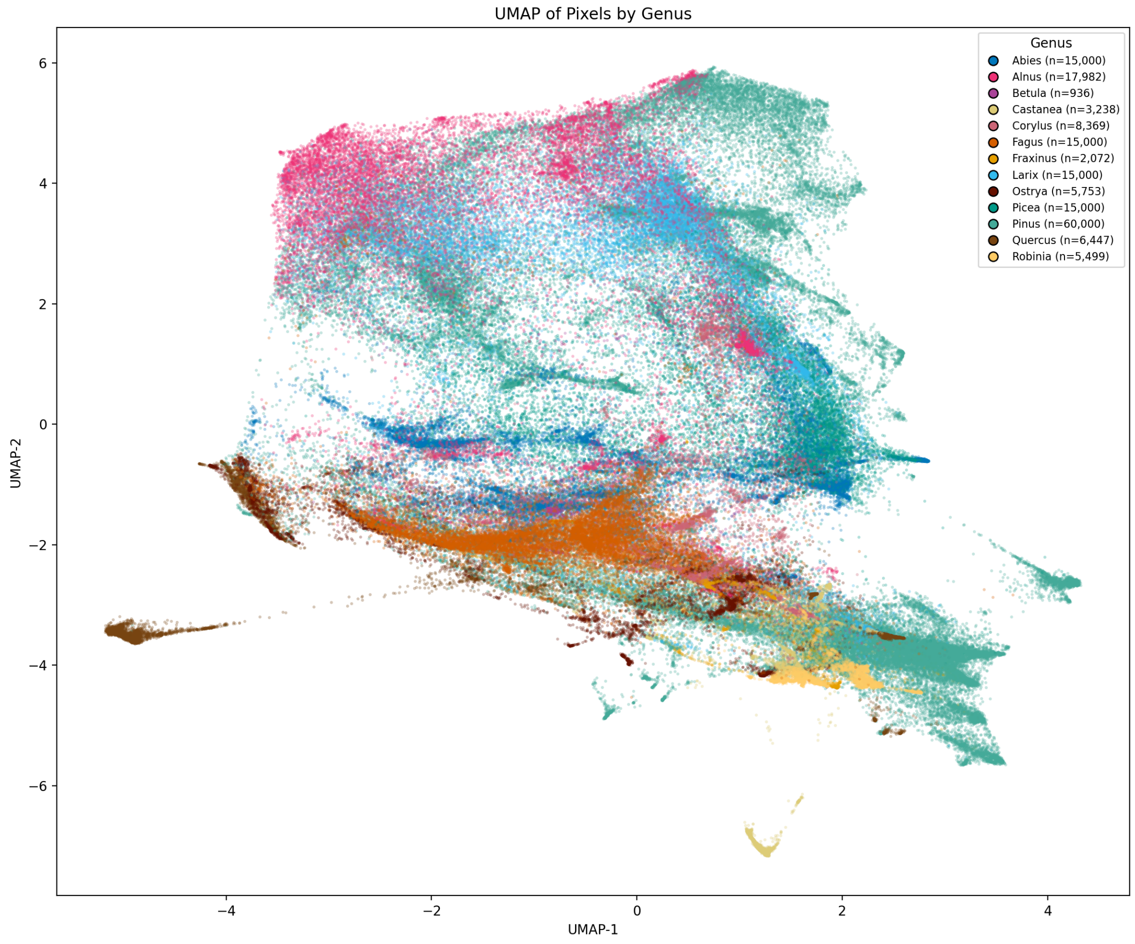

Figure S.15: UMAP projection of Tessera 2018 pixel embeddings coloured by genus (pre-merge). Genera occupy coherent regions of the embedding space, with within-genus separation aligned with known ecological and elevational niches.

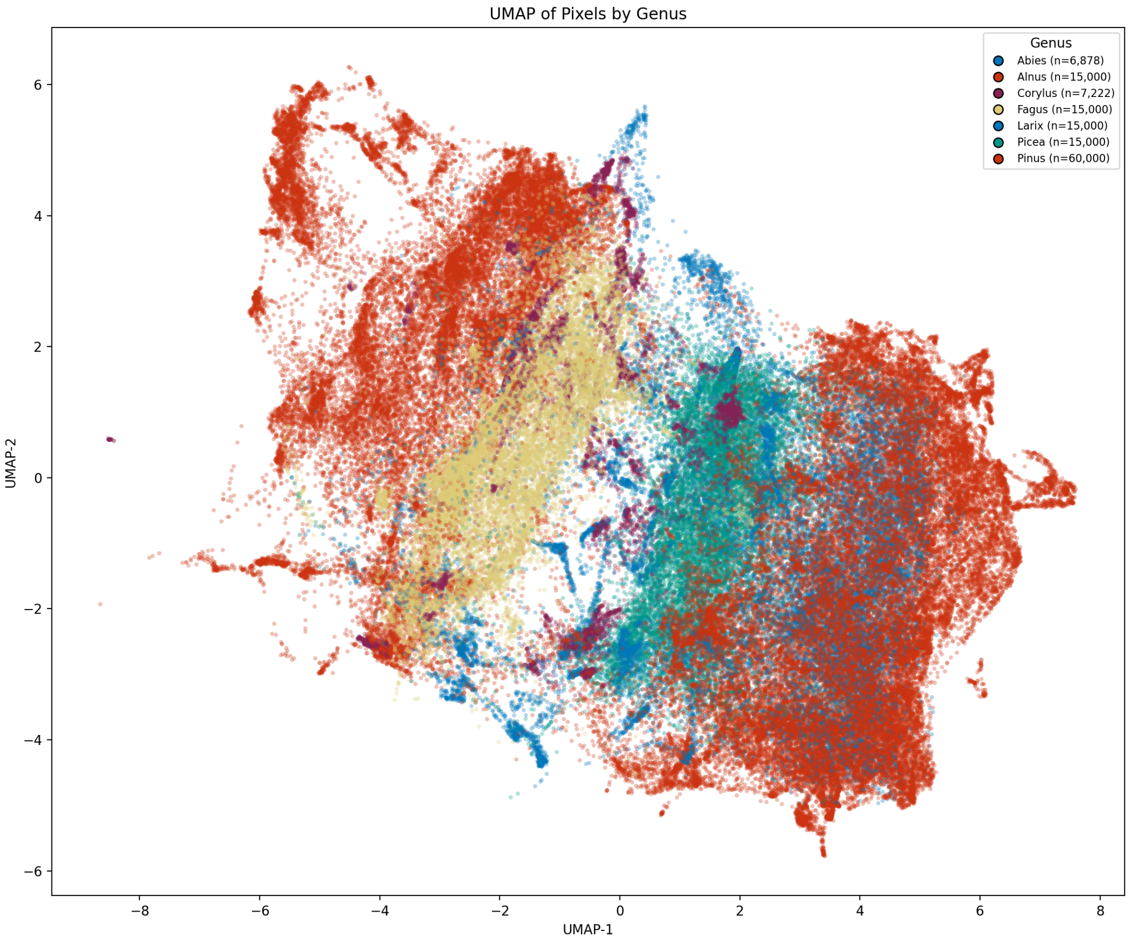

Figure S.16: UMAP projection of AlphaEarth 2018 pixel embeddings coloured by genus (pre-merge). AlphaEarth shows comparable genus-level clustering to Tessera (Figure S.15).

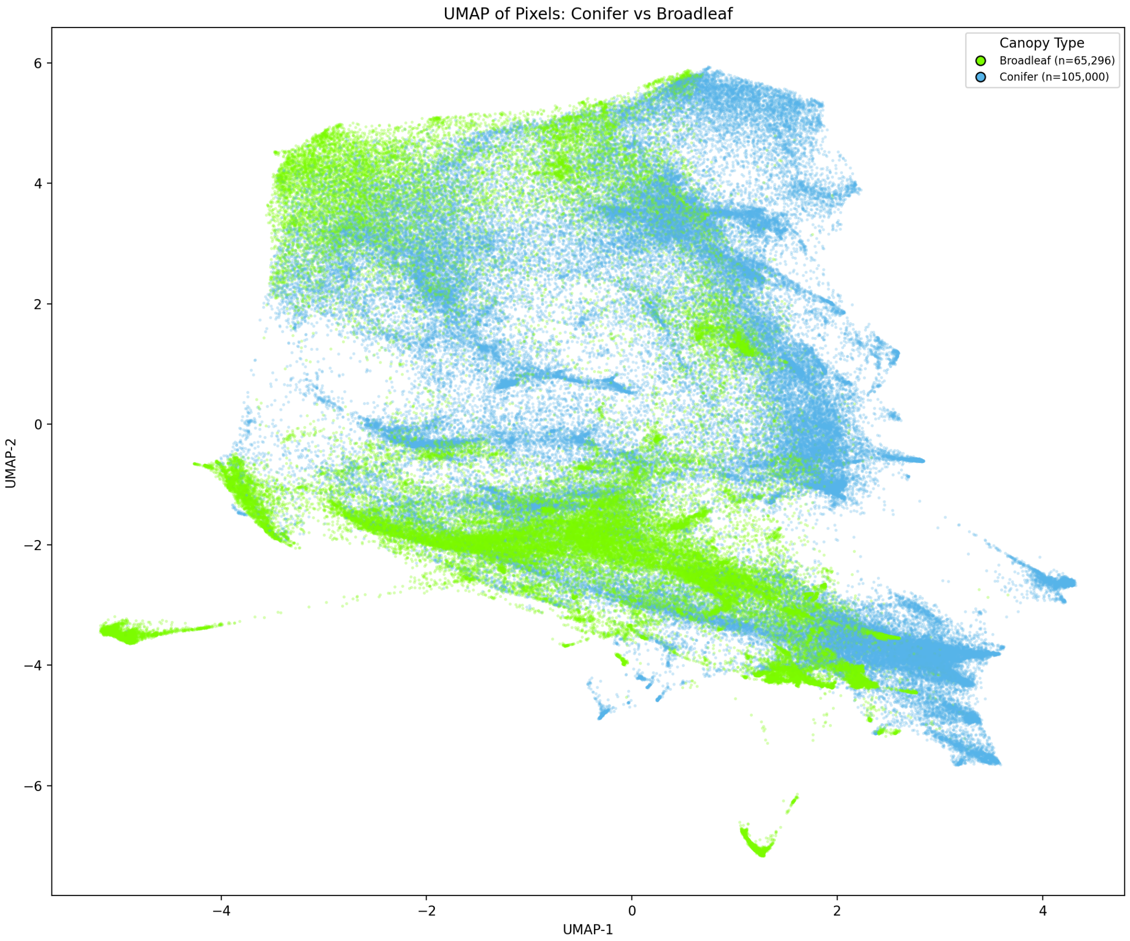

Figure S.17: UMAP projection of Tessera 2018 embeddings coloured by conifer/broadleaf functional type, showing clear separation between the two groups in the learned representation.

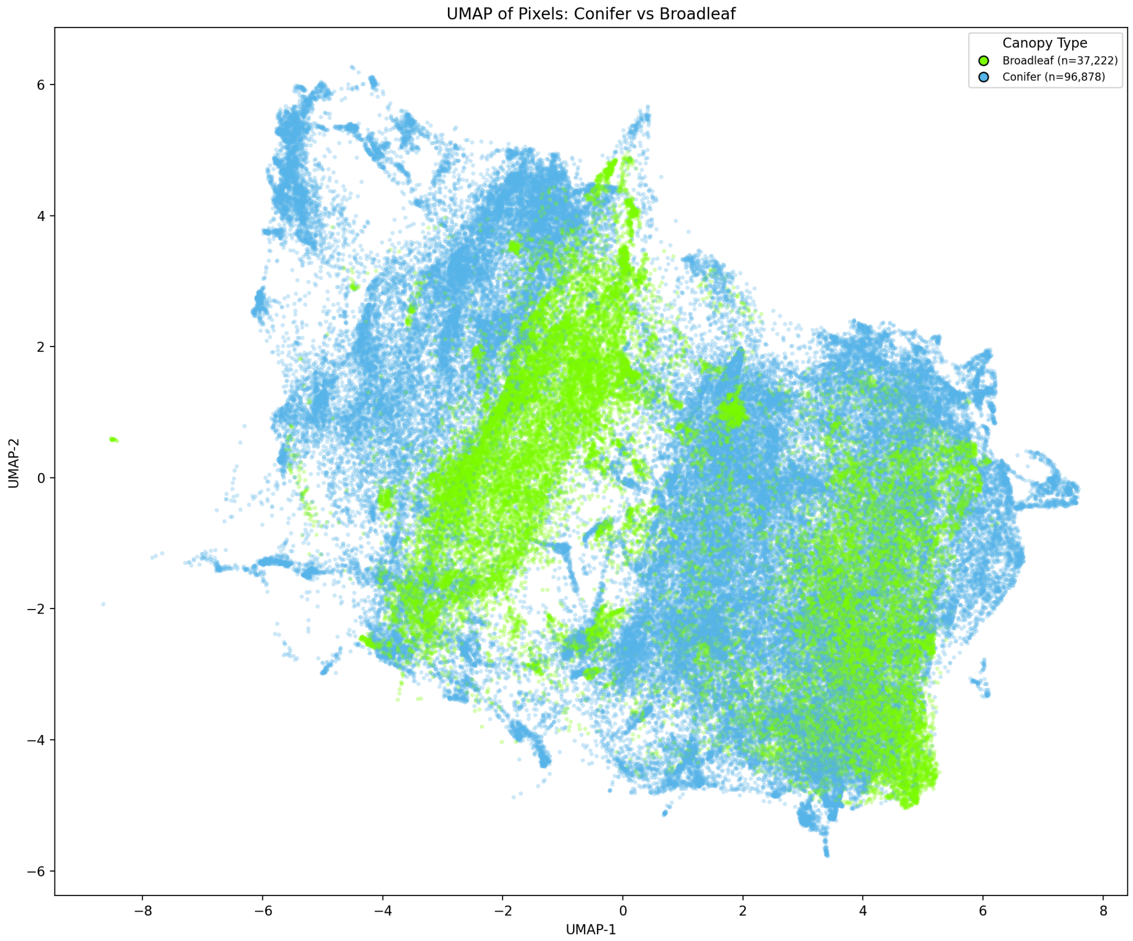

Figure S.18: UMAP projection of AlphaEarth 2018 embeddings coloured by conifer/broadleaf functional type. AlphaEarth likewise separates conifers from broadleaves, though with a somewhat different manifold geometry compared to Tessera (Figure S.17).

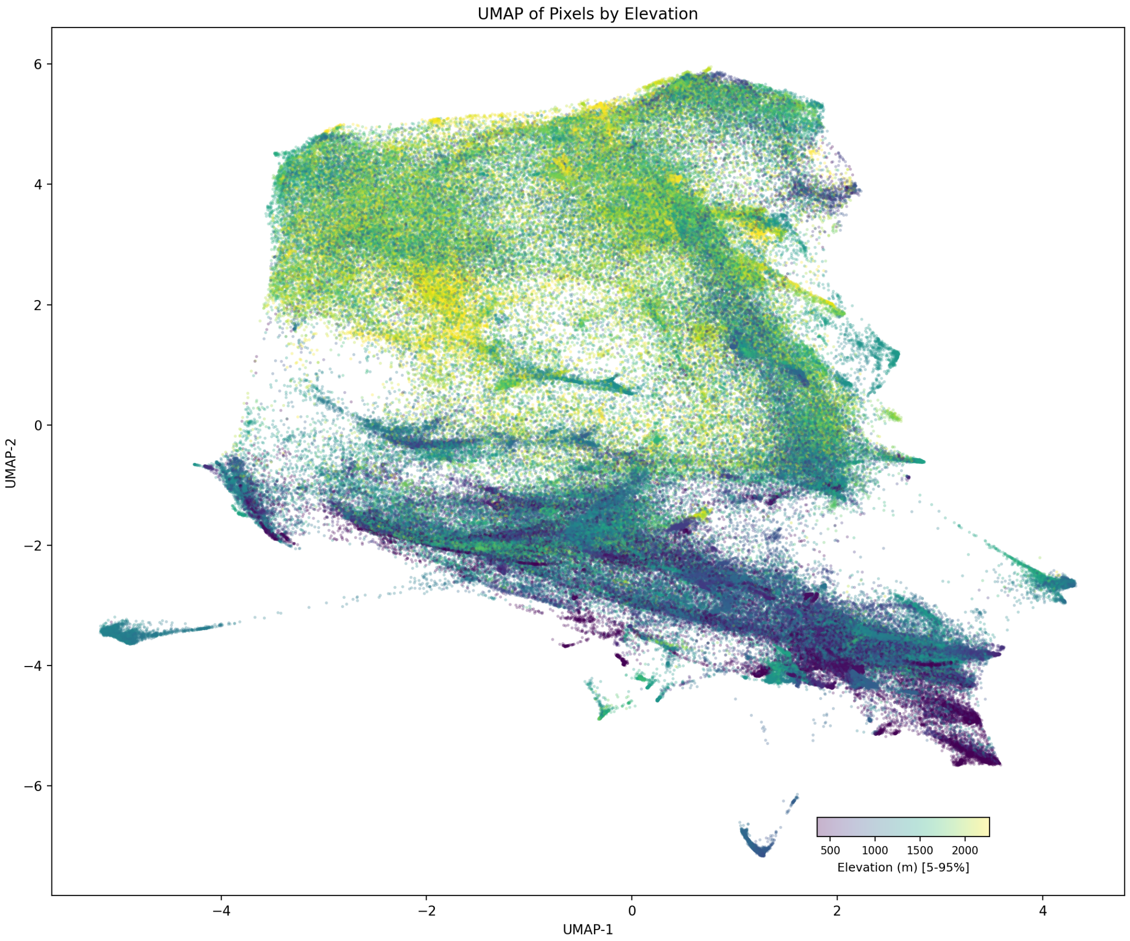

Figure S.19: UMAP projection of Tessera 2018 embeddings coloured by elevation. Elevation emerges as a dominant organising axis of the embedding space, consistent with the strong elevational zonation of tree species in Trentino.

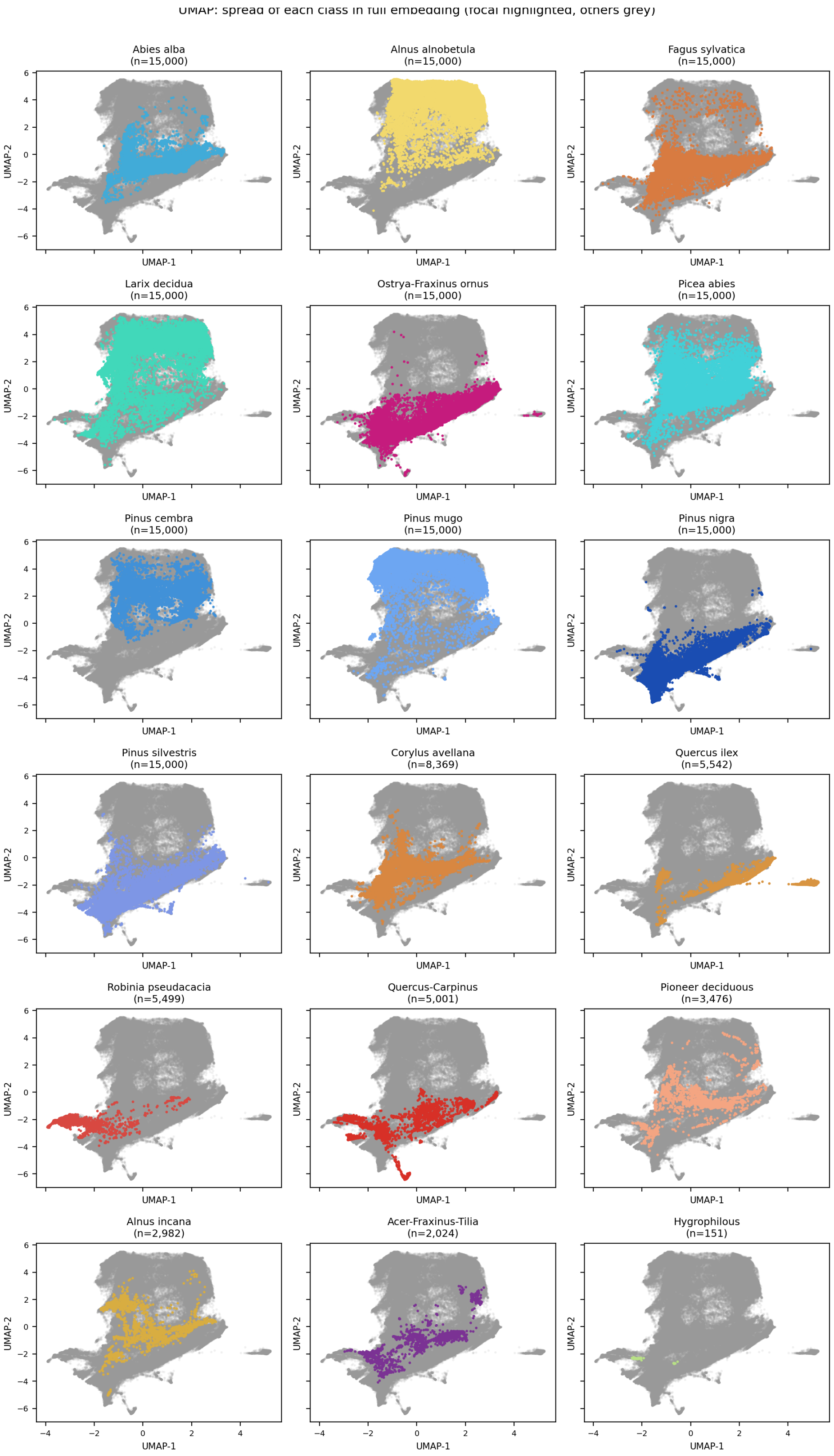

Figure S.20: Per-class spread in the Tessera 2018 UMAP embedding space, using the 18 final classes (Table S.3). Each panel highlights pixels belonging to one class (coloured) against the full embedding (grey), with sample sizes shown. Abundant conifers such as *Picea abies* and *Larix decidua* occupy large, well-separated regions, whereas rare classes such as Hygrophilous and Acer-Fraxinus-Tilia are confined to small, localised clusters.

##### S.7.3 Temporal transfer

To assess whether foundation-model embeddings generalise across years, we trained classifiers on 2018 embeddings and evaluated on 2019 embeddings (and vice versa) without retraining. This tests whether the learned representations are temporally stable despite interannual variation in phenology and acquisition conditions.

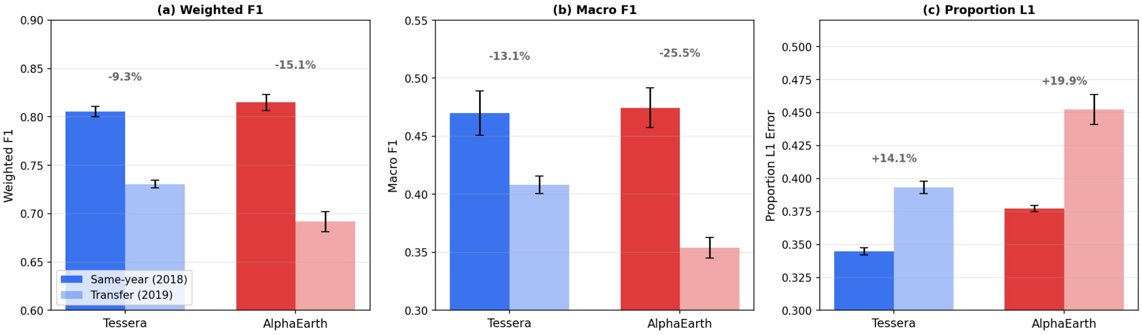

Figure S.21: Temporal transfer performance: classifiers trained on one year's embeddings and evaluated on another. Foundation-model embeddings exhibit smaller cross-year degradation than conventional composites, suggesting more temporally stable representations.

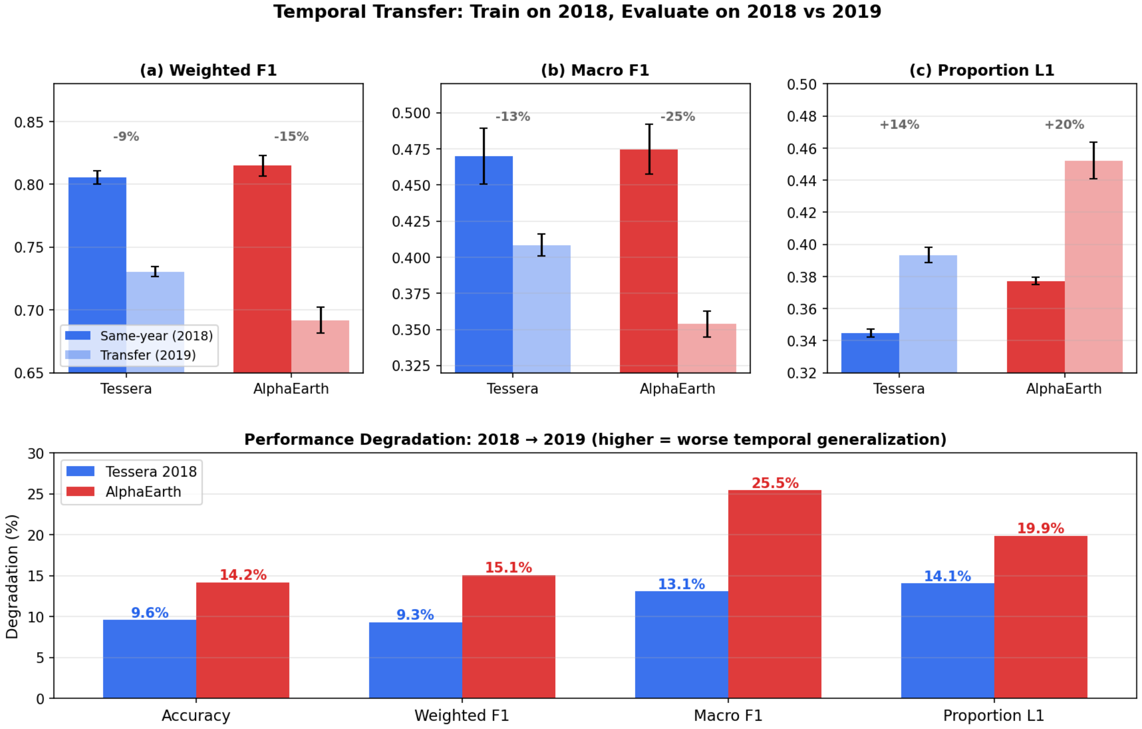

Figure S.22: Combined temporal transfer analysis across all representations and metrics, showing within-year vs. cross-year performance.

##### S.7.4 Sampling strategy comparison

The main-text fraction curves (Section S.7.1) use stratified random sampling at the parcel level to select training subsets. To explore whether more informed selection strategies can improve label efficiency – particularly in the extreme low-data regime relevant to few-shot species mapping – we repeated the fraction-curve analysis under five alternative sampling strategies:

(i) **Random stratified**: class-balanced random parcel selection (baseline, identical to the main-text experiments).

(ii) **Coreset K-means**: parcels whose mean embedding is closest to the $K$-means cluster centres in embedding space, selecting parcels that are maximally representative of the learned feature distribution.

(iii) **Altitude stratified**: parcels sampled proportionally across elevation bands, ensuring coverage of the full altitudinal gradient.

(iv) **Spatially diverse**: parcels selected to maximise geographic spread using farthest-point sampling in coordinate space.

(v) **Active learning**: iterative uncertainty-based selection, where an initial model identifies the parcels on which predictions are most uncertain, and these are added to the training set.

All strategies used the same hyperparameters, cross-validation folds, and evaluation protocol as the main-text fraction curves. Each configuration was evaluated across 3 seeds $\times$ 5 folds = 15 runs per fraction.

Table S.14 reports weighted F1 and macro F1 at selected training fractions for both foundation models. Coreset K-means selection consistently outperforms random stratified sampling, with the largest advantages at the lowest data fractions: +1.3 percentage points (pp) in weighted F1 at 1% for Tessera, and +1.4 pp for AlphaEarth. By 50% the differences are negligible, and all strategies converge at 100% as expected. Notably, spatially diverse and active-learning strategies perform *below* random in the low-label regime, suggesting that representativeness in embedding space is more important than spatial coverage or decision-boundary refinement when training data are scarce.

Figures S.23 and S.24 visualise these fraction curves for each model.

Table S.14: Sampling strategy comparison: weighted F1 and macro F1 at selected training-data fractions for Tessera and AlphaEarth embeddings. Values are mean ± std across 15 runs (3 seeds × 5 folds). Bold indicates the best strategy at each fraction (excluding 100%, where all converge). 100% corresponds to ∼45,200 training parcels.

| | | Training fraction | | | | |
|---|---|---|---|---|---|---|
| Model | Strategy | 0.1% | 1% | 5% | 10% | 100% |
| *Weighted F1* | | | | | | |
| Tessera | Random stratified | .622±.017 | .720±.011 | .754±.010 | .769±.009 | .805±.006 |
| | **Coreset K-means** | **.655**±.023 | **.733**±.009 | **.770**±.008 | **.782**±.007 | .805±.006 |
| | Altitude stratified | .619±.027 | .721±.015 | .753±.008 | .769±.008 | .805±.006 |
| | Spatially diverse | .569±.034 | .641±.018 | .710±.012 | .741±.010 | .805±.006 |
| | Active learning | .622±.017 | .680±.016 | .706±.012 | .728±.012 | .805±.006 |
| AlphaEarth | Random stratified | .604±.025 | .708±.012 | .746±.007 | .765±.005 | .814±.003 |
| | **Coreset K-means** | **.611**±.024 | **.722**±.008 | **.763**±.005 | **.776**±.005 | .814±.003 |
| | Altitude stratified | .605±.024 | .709±.012 | .747±.009 | .761±.005 | .814±.003 |
| | Spatially diverse | .529±.035 | .630±.024 | .678±.011 | .717±.008 | .814±.003 |
| | Active learning | .605±.024 | .676±.014 | .688±.017 | .709±.017 | .814±.003 |
| *Macro F1* | | | | | | |
| Tessera | Random stratified | .216±.020 | .319±.014 | .382±.019 | .406±.015 | .485±.012 |
| | **Coreset K-means** | **.223**±.012 | **.332**±.013 | **.399**±.013 | **.431**±.010 | .485±.011 |
| | Altitude stratified | **.259**±.019 | **.344**±.018 | .389±.014 | .411±.011 | .485±.011 |
| | Spatially diverse | .205±.018 | .265±.011 | .331±.020 | .356±.014 | .485±.011 |
| | Active learning | .216±.020 | .279±.013 | .356±.011 | .399±.010 | .485±.011 |
| AlphaEarth | Random stratified | .190±.018 | .291±.009 | .359±.015 | .393±.010 | .483±.013 |
| | **Coreset K-means** | .165±.010 | **.294**±.014 | **.379**±.015 | **.406**±.015 | .483±.013 |
| | Altitude stratified | **.224**±.011 | **.298**±.013 | .365±.012 | .391±.012 | .483±.013 |
| | Spatially diverse | .162±.019 | .245±.017 | .297±.020 | .328±.015 | .483±.013 |
| | Active learning | .192±.019 | .261±.016 | .312±.019 | .350±.018 | .483±.013 |

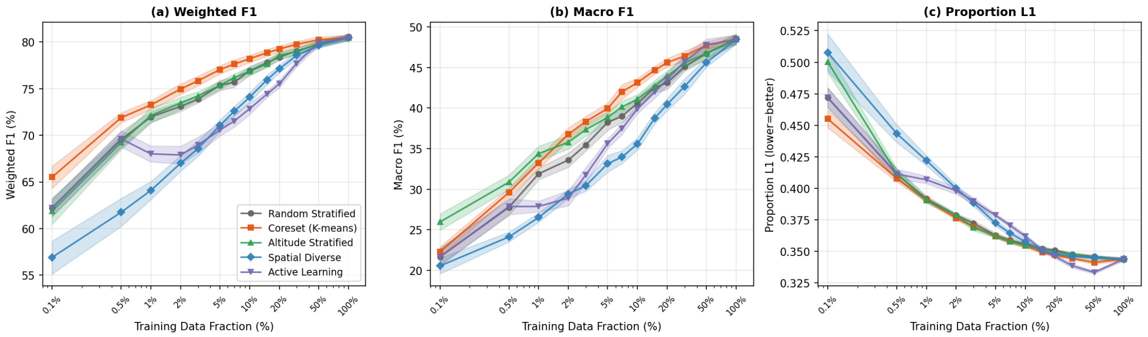

Figure S.23: Fraction curves under alternative sampling strategies for Tessera embeddings. Coreset K-means selection (embedding-guided) consistently outperforms random stratified sampling, while spatially diverse and active-learning strategies underperform in the low-data regime. Shaded regions show ±1 standard deviation across 15 runs.

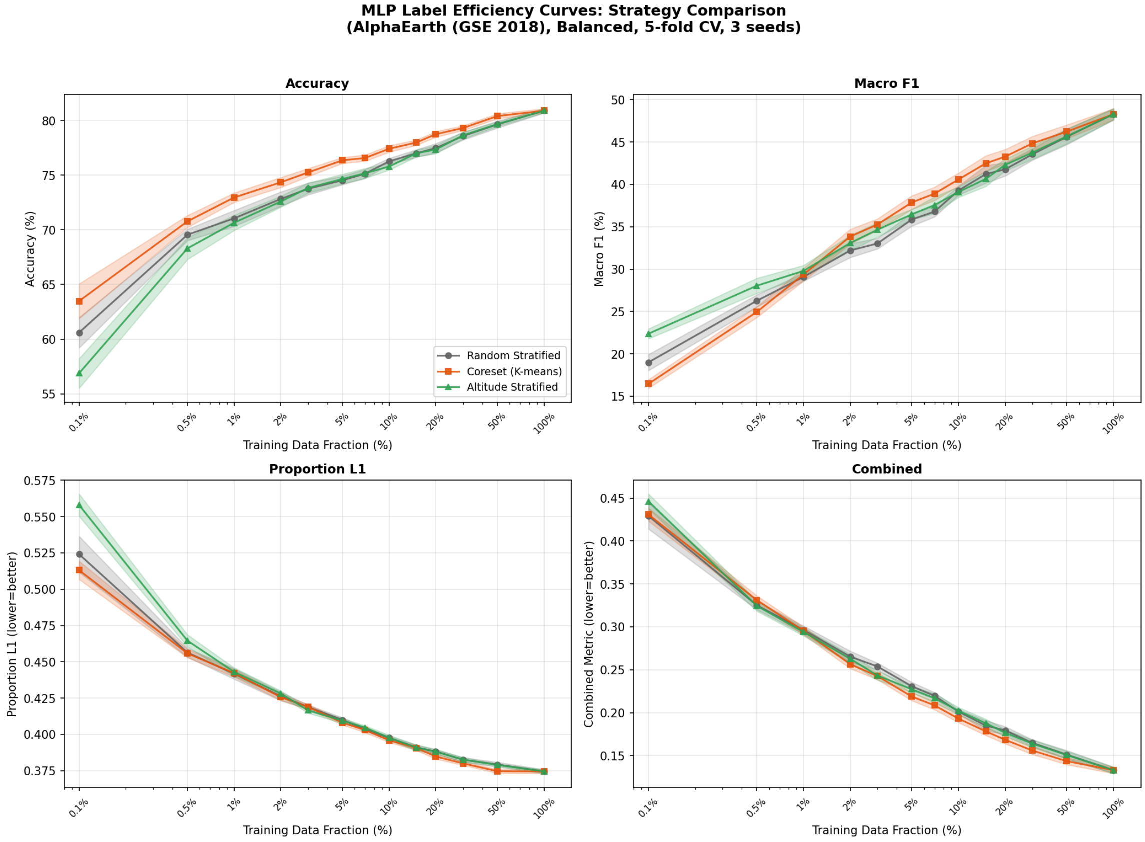

Figure S.24: Fraction curves under alternative sampling strategies for AlphaEarth embeddings. The same pattern holds: coreset K-means is the strongest strategy in the low-data regime, while spatial diversity and active learning offer no advantage over random selection.
